# Descriptions of four new species of *Minyomerus* Horn, 1876 sec. Jansen & Franz, 2018 (Coleoptera: Curculionidae), with notes on their distribution and phylogeny

**DOI:** 10.1101/383091

**Authors:** M. Andrew Jansen, Nico M. Franz

## Abstract

This contribution adopts the taxonomic concept approach, including the use of *taxonomic concept labels* (name sec. [*according to*] source) and Region Connection Calculus (RCC-5) articulations and alignments. Prior to this study, the broad-nosed weevil genus *Minyomerus* Horn, 1876 sec. Jansen & Franz, 2015 (Curculionidae [non-focal]: Entiminae [non-focal]: Tanymecini [non-focal]) contained 17 species distributed throughout the desert and plains regions of North America. In this review of *Minyomerus* sec. Jansen & Franz, 2018, we describe the following four species as new to science: *Minyomerus ampullaceus* sec. Jansen & Franz, 2018 (henceforth: [JF2018]), **new species**, *Minyomerus franko* [JF2018], **new species**, *Minyomerus sculptilis* [JF2018], **new species**, and *Minyomerus tylotos* [JF2018], **new species**. The four new species are added to, and integrated with, the preceding revision, and an updated key and phylogeny of *Minyomerus* [JF2018] are presented. A cladistic analysis using 52 morphological characters of 26 terminal taxa (5/21 outgroup/ingroup) yielded a single most-parsimonious cladogram (Length = 99 steps, Consistency Index = 60, Retention Index = 80). The analysis reaffirms the monophyly of *Minyomerus* [JF2018] with eight unreversed synapomorphies. The species-group placements, possible biogeographic origins, and natural history of the new species are discussed in detail.

## INTRODUCTION

This phylogenetic study follows Jansen & Franz (2015) in the use of the taxonomic concept approach; see Franz & Peet (2009), Franz et al. (2016a,b). Accordingly:

1. *Taxonomic concept labels* – i.e., the taxonomic name sec. (*according to*) author or source (year) – are used whenever we identify one specific usage of the taxonomic name. Examples: *Minyomerus* Horn, 1876 sec. Jansen & Franz, 2015 (henceforth: [JF2015]) and *Minyomerus* Horn, 1876 sec. Jansen & Franz, 2018 (henceforth: [JF2018]). We also employ this convention to express nomenclatural relationships.
2. Solely the taxonomic name – without the sec. annotation – is used to refer to the cumulative history (origin to present) of taxonomic concept labels in which that name participates. Example: *Minyomerus* Horn, 1876.
3. The annotation [non-focal] is added to taxonomic names whose meanings are not under scrutiny in the present context; such as names for higher-level weevil groups and associated plants (exempting common names). Example: Tanymecini Lacordaire, 1863 [non-focal].

The weevil genus *Minyomerus* Horn, 1876 [JF2018] remains currently assigned to the tribe Tanymecini Lacordaire, 1863 [non-focal], subtribe Tanymecina Lacoirdaire, 1863 [non-focal] (Curculionidae [non-focal]: Entiminae [non-focal] – higher-level classification in accordance with Alonso-Zarazaga & Lyal 1999 and Bouchard et al. 2011). A recent phylogenetic revision of the genus *Minyomerus* [JF2015] recognized a total of 17 described species, distributed throughout the desert and plains regions of North America (Jansen & Franz 2015).

Members of the genus *Minyomerus* [JF2018] are phytophagous, and may be found on a variety of host plants, especially the creosote bush *Larrea tridentata* (DC) Coville [non-focal] (Zygophyllaceae [non-focal]), broomweed *Gutierrezia* Lagasca [non-focal] (Asteraceae [non-focal]), sagebrush *Artemisia* Linnaeus [non-focal] (Asteraceae [non-focal]), and occasionally on other various members of Asteraceae [non-focal] (Jansen & Franz 2015). While many species appear to be generalists, the adults are consistently observed on the leaves and branches of the host, feeding on the leaf tissue. All other life stages remain unknown. Species of *Minyomerus* [2018] are commonly found in deserts throughout western North America; including the Mojave, Sonoran, Chihuahuan, and Great Basin Deserts. However, their distributional range extends throughout the semi-arid regions of the Great Plains, the Colorado Plateau, and Baja California, Mexico (O’Brien & Wibmer 1982, Jansen & Franz 2015). The adults are flightless, as the hind wings and associated flight structures of all species are either greatly reduced or not readily apparent in dissection.

*Minyomerus* [JF2018] belongs to the broad-nosed weevils, subfamily Entiminae [non-focal], on the basis of having a short, broad rostrum and dehiscent mandibular process (Marvaldi 1997, Anderson 2002, Oberprieler et al. 2007, 2014, Marvaldi et al. 2014). The adults are clothed in appressed, circular scales, generally in earth-tones from white to dark brown, with sub-recumbent to erect, interspersed setiform scales (“setae”) arranged in rows on the elytral intervals. Their body length can range from 2.8 mm to 6.0 mm (Jansen & Franz 2015). The genus has been classified in the tribe Tanymecini [non-focal] based on the presence of post-ocular vibrissae that project anteriorly from the anterior prothoracic margin, although the exact placement and sister taxa of this genus within the tribe are currently unknown (Howden 1959, 1970, 1982, Jansen & Franz 2015).

*Minyomerus* [JF2015] was circumscribed by a unique combination of synapomorphic traits, described by Jansen & Franz (2015) as follows:

1. The integument is covered by appressed scales that are sub-circular and overlap posteriorly.
2. The nasal plate is present as a broad, scale-covered, chevron-shaped ridge demarcating the epistoma.
3. A sulcus posteriad of nasal plate is present.
4. The scrobe is sub-equal in length to the funicle and club combined.
5. The head is directed slightly ventrally.
6. The metatibial apex lacks setiform bristles yet displays bristles that are shorter to sub-equal in length to the surrounding setae and conical to lamelliform.
7. The mesotarsi are slightly shorter than the mesotibiae.
8. All tarsi lack pads of setiform setae but have stout, spiniform setae.

The following additional characters are useful for identifying members of *Minyomerus* [JF2018], especially when differentiating the former from other genera of Tanymecini [non-focal] that may be found together in the same desert habitats; viz. *Isodrusus* Sharp, 1911 [non-focal], *Isodacrys* Sharp, 1911 [non-focal], and *Pandeleteinus* Champion, 1911 [non-focal] (see also Anderson 2002):

1. The intercoxal process of the prosternum is medially divided into two halves, with the procoxae apparently contiguous in most.
2. The elytral humeri are rounded rather than angled and protruding.
3. The profemora are not dilated and lack spines.
4. The protibiae are ventrally excavated by a longitudinal groove or concavity.
5. A distinct scrobe is present and directed ventrad of the eye, with a more or less apparent tooth formed by an overhang of the dorsal margin.

Following the publication of a monographic revision of *Minyomerus* [JF2015], we have discovered four additional, undescribed species. These are known to us only from limited numbers of specimens, yet are well circumscribed by – i.e., intensionally included in (see Franz & Peet 2009) – the recent generic delimitation of *Minyomerus* [JF2015]. In other words, the addition of these new species *has not* required altering the intensional, property-based definition of the genus-level concept as circumscribed in Jansen & Franz (2015) (see **Phylogenetic Results**). Our RCC-5 alignments (see **RCC-5 Alignments**) reflect this genus-level concept congruence while also showing which classificatory and phylogenetic structures have changed (Figs. 32-34). The precise use of the taxonomic concept labels in accordance with either [JF2015] or [JF2018] is meant to minimize the creation of new taxonomic concept labels (to counter label “inflation”; see Franz & Peet (2009)), while reflecting explicitly *which* taxonomic concepts we consider as relevantly new and unique to the present study.

Here we describe the four newly found species of *Minyomerus* [JF2018] and provide images of the holotypes and of dissected genitalia for the purpose of identification. We additionally conduct a morphological phylogenetic analysis of the genus to clarify the placement of these new taxa within *Minyomerus* [JF2018], based on the analysis provided in our previous work. An emended identification key to the species of *Minyomerus* [JF2018] is given, along with an updated species checklist. Where possible, we make note of host-plant records, and briefly discuss the geographic distributions of the herein described species. A more extensive discussion of the habits, distribution, and delimitation of the genus *Minyomerus* [JF2015] and all of its constituent species is provided in Jansen & Franz (2015).

## MATERIALS AND METHODS

The methods used in this manuscript are generally consistent with Jansen & Franz (2015). Relevant updates are detailed below. In particular, we retain the format for the species descriptions, emphasizing only those characters that vary significantly from the generic circumscription of *Minyomerus* [JF2015].

### Acquisition of museum specimens

The set of specimens used in Jansen & Franz (2015) was supplemented with material from the following collections, using the codens of Arnett Jr. et al. (1993):

**CMNC** Canadian Museum of Nature Collection, Ottawa, Ontario, Canada
**TAMU** Texas A & M University, College Station, Texas, USA
**USNM** National Museum of Natural History, Washington, D.C., USA

Georeferencing of localities was performed with Google Earth (Google Inc. 2018), following the WGS 84 standard, and reported in decimal degrees. Taxonomic names for associated host plants, as noted following each species account, are used in accordance with Munz & Keck (1973) and SEINet (2018).

### Morphological analysis

Our systematic and descriptive approach is complementary to Jansen & Franz (2015), which in turn follows Franz (2010*a,b*, 2012). The terminology for exterior morphology is in general accordance with de la Torre-Bueno et al. (1989). Additional morphological terms specific to broad-nosed weevils (Entiminae [non-focal]) were used as follows: Ting (1936) and Morimoto & Kojima (2003) for mouthparts; Thompson (1992) for tibial apices and abdominal segments; and Oberprieler et al. (2014) and Howden (1995) for male and female terminalia.

Measurements were taken with a Leica M205 C stereomicroscope and associated software, Leica Application Suite (LAS), version 4.1.0. Overall body length and width were measured in dorsal view as the maximum distance between the rostral and elytral apices, and the maximum width of both elytra, respectively. Rostral length was measured in dorsal view as the distance between the epistomal apex and the anterior margin of the eyes. Rostral width was measured in dorsal view as the maximum distance between the dorsal margins of the rostrum near the point of antennal insertion. Pronotal length was measured in dorsal view as the length along the midline between the anterior and posterior margins. The width of an individual elytron was measured in dorsal view as the maximum distance between the lateral margin and the elytral suture. Other length and width measurements were also performed in dorsal orientation, using the maximum length and width of the corresponding structure (profemur, protibia, elytron, and aedeagus). Images of mouthparts and terminalia were produced with the Leica microscope equipment, while habitus photographs were created with a Visionary Digital Passport II system using a Canon EOS Mark 5D II camera.

The herein newly recognized species of *Minyomerus* [JF2018] were delimited through application of the phylogenetic species concept *sensu* Wheeler & Platnick (2000). Species descriptions are in alphabetical order, rather than phylogenetic order, for ease of use. As in Jansen & Franz (2015), the species descriptions represent unique, complementary accounts of the character states observed for each species, including their intra-specific variability, but excepting characters invariant within the genus-level concept of *Minyomerus* [JF2015]. Likewise, descriptions of males emphasize characters that are variable and sufficiently different from those of the females to merit recognition. The key to identifying species of *Minyomerus* [JF2018] is arranged with emphasis being placed on the most readily observable diagnostic characters. This manuscript is arranged with the species descriptions appearing first, followed by the key to species, and then by the phylogenetic and RCC-5 alignment results.

### Phylogenetic analysis

The morphological cladistic analysis includes 26 terminal taxa; with 21 ingroup and 5 outgroup terminals. The ingroup terminals were represented by 17 species previously assigned to *Minyomerus* [JF2015] and four newly recognized species. In keeping with our previous analysis, we sampled outgroups fairly broadly while remaining focused on North American lineages that are putative close relatives of the ingroup (Jansen & Franz 2015, Nixon & Carpenter 1993).

Although the tribe Tanymecini [non-focal] is cosmopolitan, the majority of New World species diversity in the tribe may be found in the subtribe Tanymecina [non-focal] (Alonso-Zarazaga & Lyal 1999). Thus, four of the five outgroup terminals are represented by species belonging to separate genera in the Tanymecina [non-focal]; viz. *Isodacrys buchanani* Howden, 1961 [non-focal], *Isodrusus debilis* Sharp, 1911 [non-focal], *Pandeleteinus subcancer* Howden, 1969 [non-focal], and *Pandeleteius cinereus* (Horn, 1876) [non-focal]. Because generic relationships in the Tanymecini [non-focal] remain unresolved, we selected a relatively far-removed taxon to root the cladogram that would nevertheless display states applicable to the ingroup for characters under consideration (Rieppel 2007, Franz 2014). To this end we used the North American species *Sitona californicus* (Fahraeus, 1840) [non-focal], of the tribe Sitonini Gistel, 1856 [non-focal].

The character matrix was edited and phylogenetic results viewed using the WinDada and WinClados interfaces of WinClada, respectively (Nixon et al. 2002). Characters are numbered in accordance with descriptive sequence used in the species accounts. A “–” symbol indicates inapplicable (character, state), whereas a “?” symbol indicates missing information, e.g., due to the unavailability of male specimens or insufficient specimens on hand to permit full dissections. Characters 9, 27, 39,45 - 47,49, and 51 were mapped onto the preferred phylogeny using ACCTRAN optimization (see Agnarsson & Miller 2008), and the remaining characters had an unambiguous optimization. All multi-state characters but one were coded as additive, as explained beneath the description for each character (see **Phylogenetic Results**), based on their alignment with the preferred phylogeny. Each alternative coding scheme was tested both alone and in unison with the other multi-state characters to assess their impact on the topology of the preferred phylogeny.

The most parsimonious tree and character state optimizations were inferred under parsimony using NONA (Goloboff 1999). An unconstrained heuristic search was conducted using the commands: `hold 100001, mult*10 0 0, hold/100`, with `mult*max*` selected. Bootstrap support was inferred in WinClada using the parameters of `1000 replications, hold 1000, hold/100, mult*10, Don’t do max*`, and Save consensus. Finally, Bremer support values (Bremer et al. 1994) and relative fit difference (Goloboff & Farris 2001) were calculated in NONA using the commands: `hold 1001, sub 20, bs` for Bremer support values, and `bs*` for relative fit difference, respectively (Goloboff et al. 2008).

The motivation for providing Bremer support values and relative fit difference comes from their respective interpretations, based on how the measures are calculated, *per* Goloboff & Farris (2001). Both of these indices rely on summation of the number of favorable and contradictory characters when comparing a most-parsimonious tree to a suboptimal tree. If the step length of the *i^th^* character (*I*) of *n* total characters on the most-parsimonious tree (*L_MPT_*) is less than its corresponding step length on the suboptimal tree (*L_SUB_*), the character is designated as favorable (*f_i_*), but if the opposite is true, the character is designated as contradictory (*c_i_*), and expressed formally:

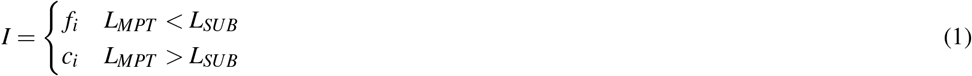

Where the number of favorable (*F*) and contradictory (*C*) characters are defined, respectively, as:

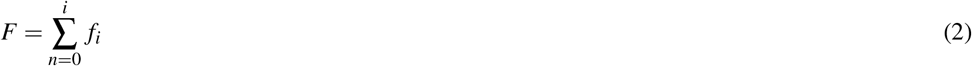

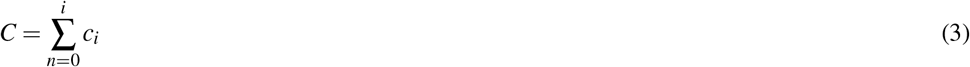

Bremer support values (bsv) and relative fit difference (rfd) are then calculated simply as:

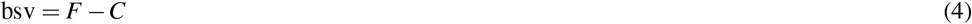

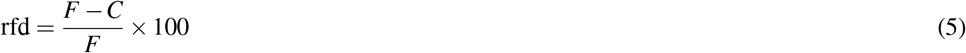

The Bremer support value for a node thus indicates how many more characters support a node than contradict it, while the relative fit difference indicates what proportion of the favorable characters are represented by the Bremer support value. Whereas the Bremer support value is as large as the number of characters supporting the node, in excess of the contradicting characters, the relative fit difference can only vary from 0 to 100, as a proportion of the number of supporting characters. By providing both measures, one may quickly discriminate, for example, between a node supported by 4 characters but contradicted by 1 character (bsv = 3, rfd = 75), and a node supported by 10 characters but contradicted by 7 characters (bsv = 3, rfd = 30).

### Taxonomic annotations and RCC-5

In accordance with Jansen & Franz (2015), we use the symbol “=“ to indicate nomenclatural synonymy (objective/subjective); and the RCC-5 symbols {==, >, <, ><, !} indicate taxonomic concept articulations. The annotations (INT) and (OST) indicate intensional and ostensive readings of articulations, and AND is used to connect multiple simultaneously recognized provenance relationships. Two *intensional* alignments are produced as part of this review, i.e., one that captures the non-/congruence of *Minyomerus* [JF2018] versus *Minyomerus* [JF2015] represented as rank-only classifications (Fig. 32), and another that represents these as fully bifurcated phylogenies with newly assigned clade concept labels, shown in whole-concept resolution (Fig. 33) and in split-concept resolution (Fig. 34); see Franz et al. (2018).

A detailed breakdown of our alignment approach and outcomes using an RCC-5 logic reasoner toolkit (Chen et al. 2014) is provided in the **Supplemental Information, SI1 to SI4**. For further information, see also Jansen & Franz (2015), Franz et al. (2016a,b).

### Species distribution modeling

We used the modeling program Maxent, Version 3.4, to generate habitat models for the species of *Minyomerus* [JF2018] (Figs. 35-38) based on documented occurrence records (Phillips et al. 2004, 2006, Elith et al. 2011). The default settings were adjusted to `Max number background points = 100, 000` and `Iterations = 10`. Cross-validation was used to leverage all available locality data; however, no models could be created for species with two or fewer documented localities. We selected 19 bioclimatic variables and elevation as Environmental Layers in Maxent, obtained from WorldClim (Hijmans et al. 2005). The layers were downloaded by tile (zones 11-13 and 21-23), with a 30 arc-second resolution (projected using WSG 84) to provide adequate coverage of the full distribution of the genus. Layerwise assembly of tiles was done using QGIS, Version 2.18.16 ‘Las Palmas’, creating composite maps of six tiles each to use in species distribution modeling (Quantum GIS Development Team 2018).

The rasterized predictive probabilities were imported into QGIS, where each file was designated a specific color. Each pixel in the raster was assigned a linearly interpolated saturation of that color, with increasing saturation denoting an increased probability of successful prediction of species presence at that point. Pixels with a value below 0.50 were rendered transparent so that the maps only show regions with a greater than 50% chance of successful prediction. The raster files were clipped to remove extraneous predicted regions based on: (1) predictive probability (i.e., removing large areas with only transparent pixels) and (2) geographic extent (accounting for endemicity). For example, a species endemic to the Snake River Valley of Idaho does not require a predictive model for bioclimatically similar habitats in the Chihuahuan Desert. Documented occurrence records are laid over the modeled habitat ranges as colored circles on their respective maps (Figs. 36-38), along with vector layers of country (white) and state (gray) borders (Hijmans et al. 2012).

### Nomenclature

The electronic version of this article in Portable Document Format (PDF) will represent a published work according to the International Commission on Zoological Nomenclature (ICZN), and hence the new names contained in the electronic version are effectively published under that Code from the electronic edition alone. This published work and the nomenclatural acts it contains have been registered in ZooBank, the online registration system for the ICZN. The ZooBank LSIDs (Life Science Identifiers) can be resolved and the associated information viewed through any standard web browser by appending the LSID to the prefix http://zoobank.org/. The LSID for this publication is: urn:lsid:zoobank.org:pub:0AEE5733-06D1-401F-88C9-0D5232FBFC7A. The online version of this work is archived and available from the following digital repositories: PeerJ, PubMed Central and CLOCKSS.

Minyomerus ampullaceus: Minyomerus franko: Minyomerus sculptilis: Minyomerus tylotos:

## DESCRIPTIONS OF NEW SPECIES

### *Minyomerus ampullaceus* Jansen & Franz sec. Jansen & Franz, 2018; sp. n

urn:lsid:zoobank.org:act:24943E17-F20E-4E3C-A3A1-A1D4D907B48E Figures 1-6

**Figure 1.**
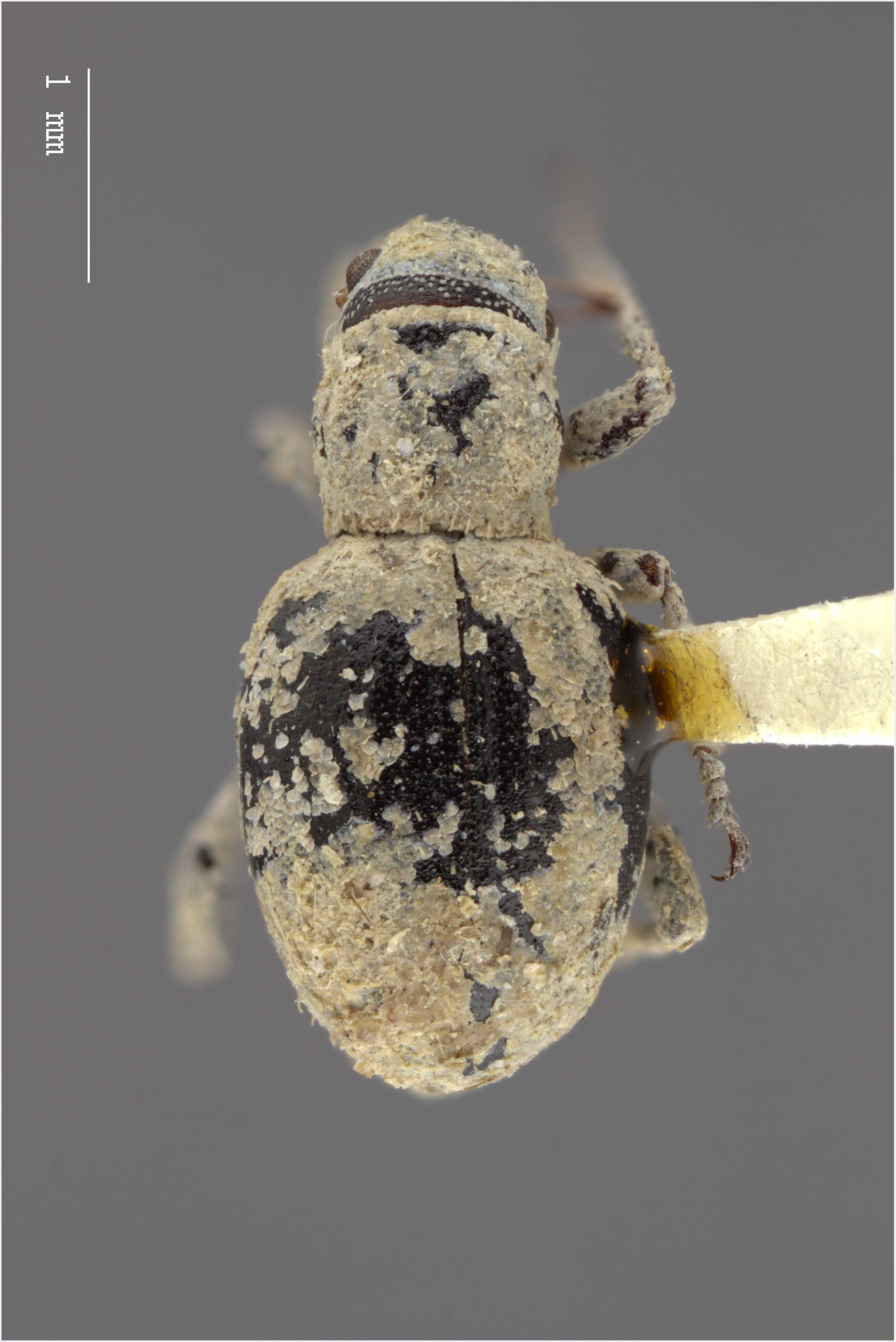
Dorsal habitus of *M. ampullaceus* [JF2018]. Image of female (♀) holotype.

**Figure 2.**
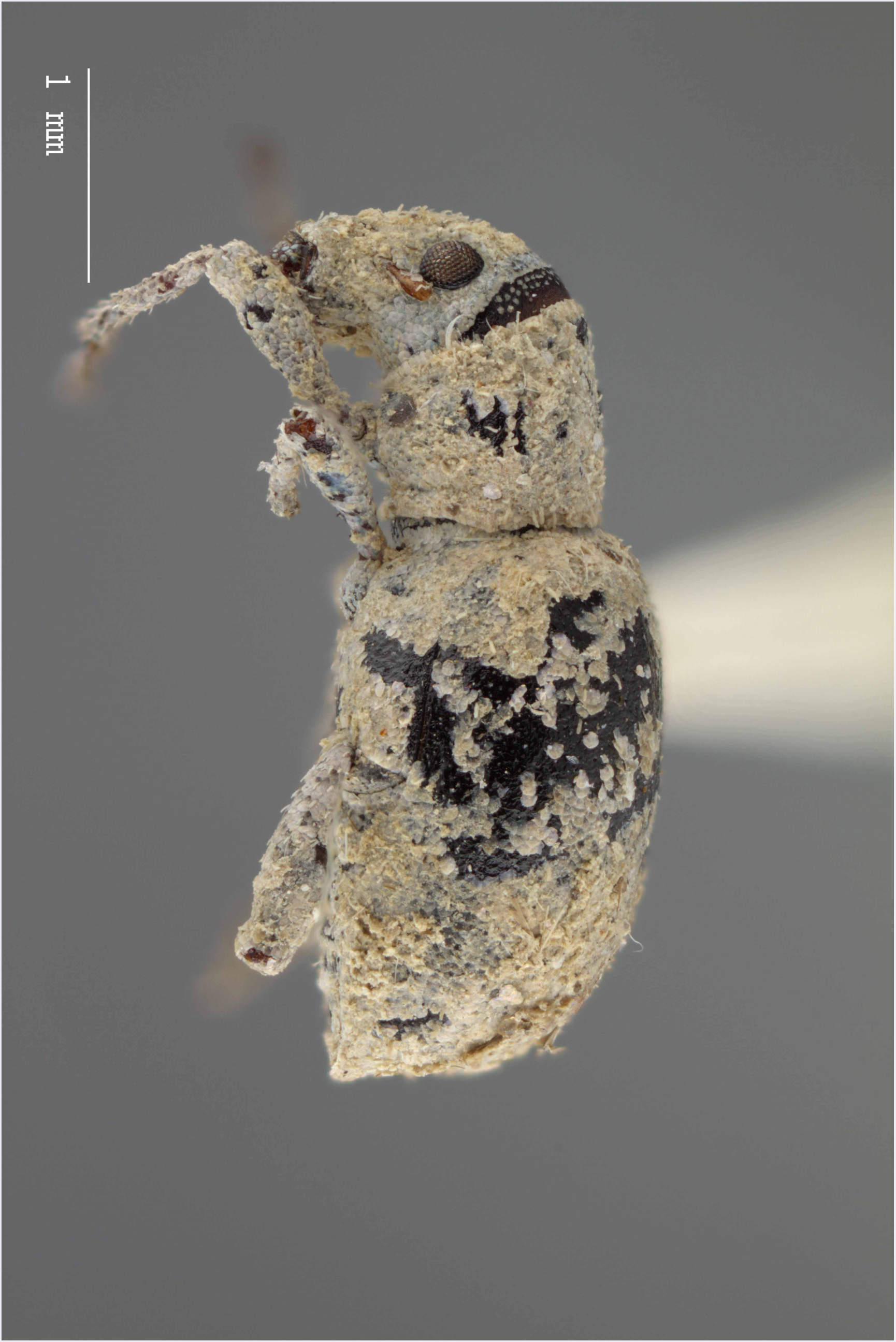
Lateral habitus of *M. ampullaceus* [JF2018]. Image of female (♀) holotype.

**Figure 3.**
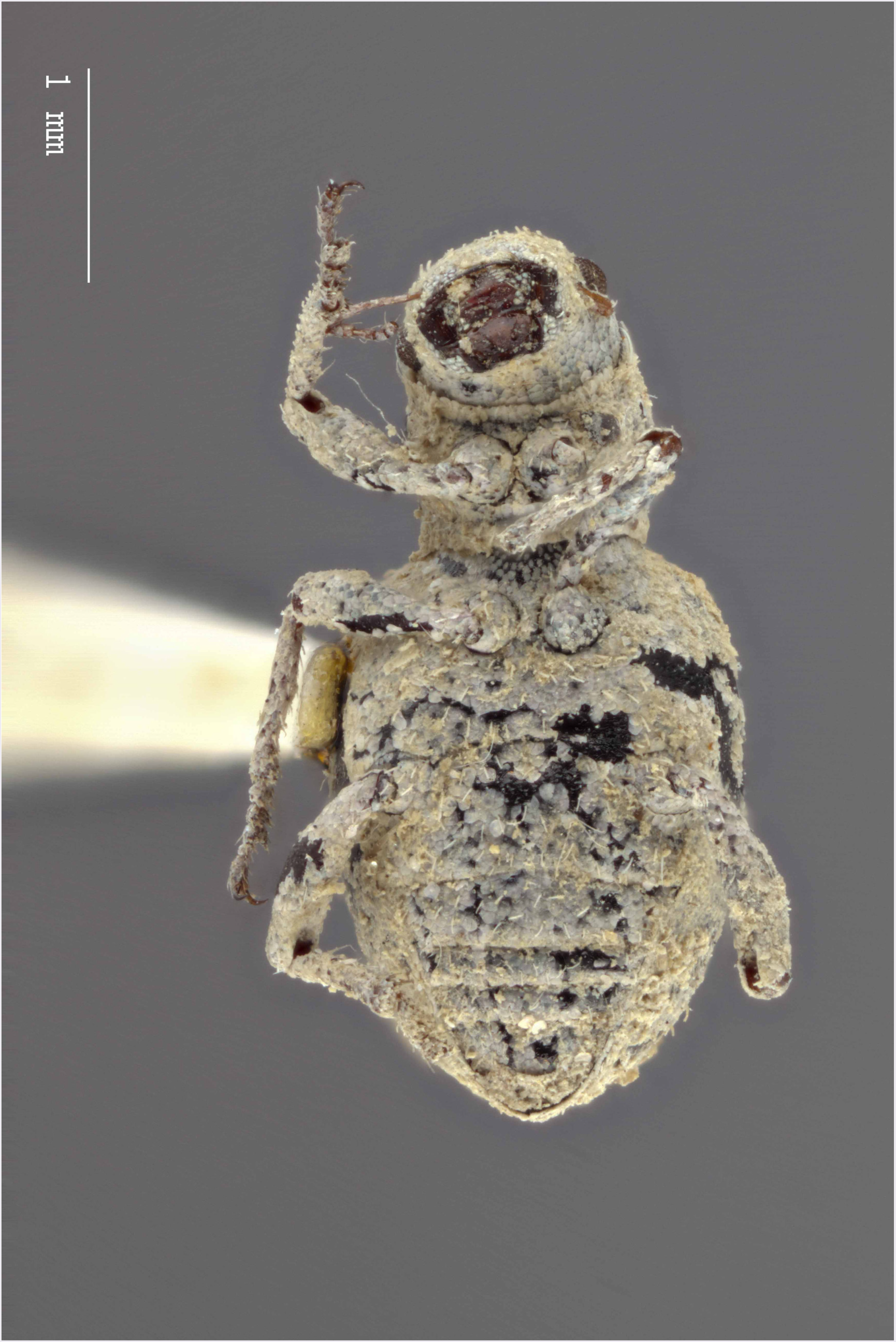
Ventral habitus of *M. ampullaceus* [JF2018]. Image of female (♀) holotype.

**Figure 4.**
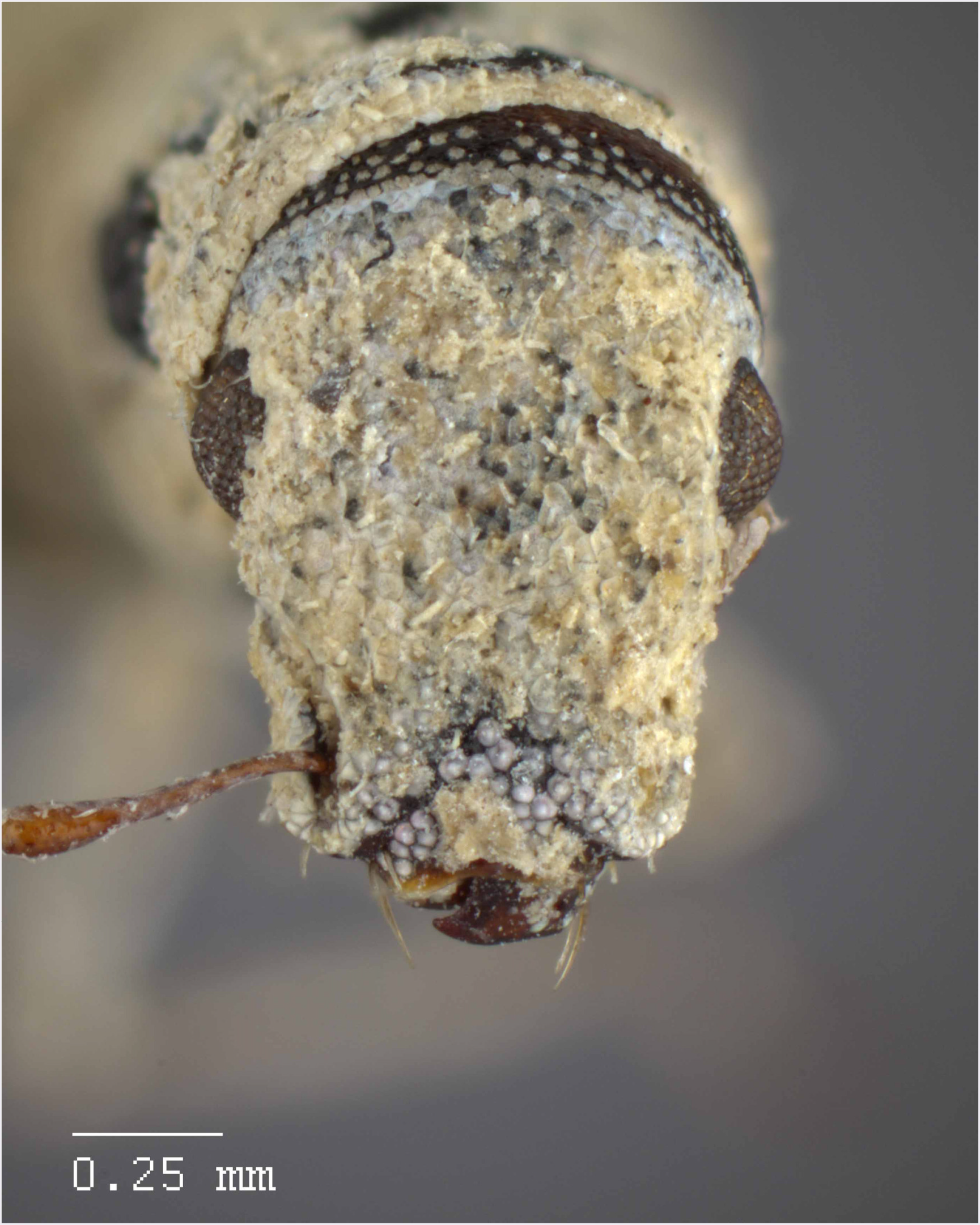
Head and rostrum of *M. ampullaceus* [JF2018]. Frontal view of female (♀) holotype.

**Figure 5.**
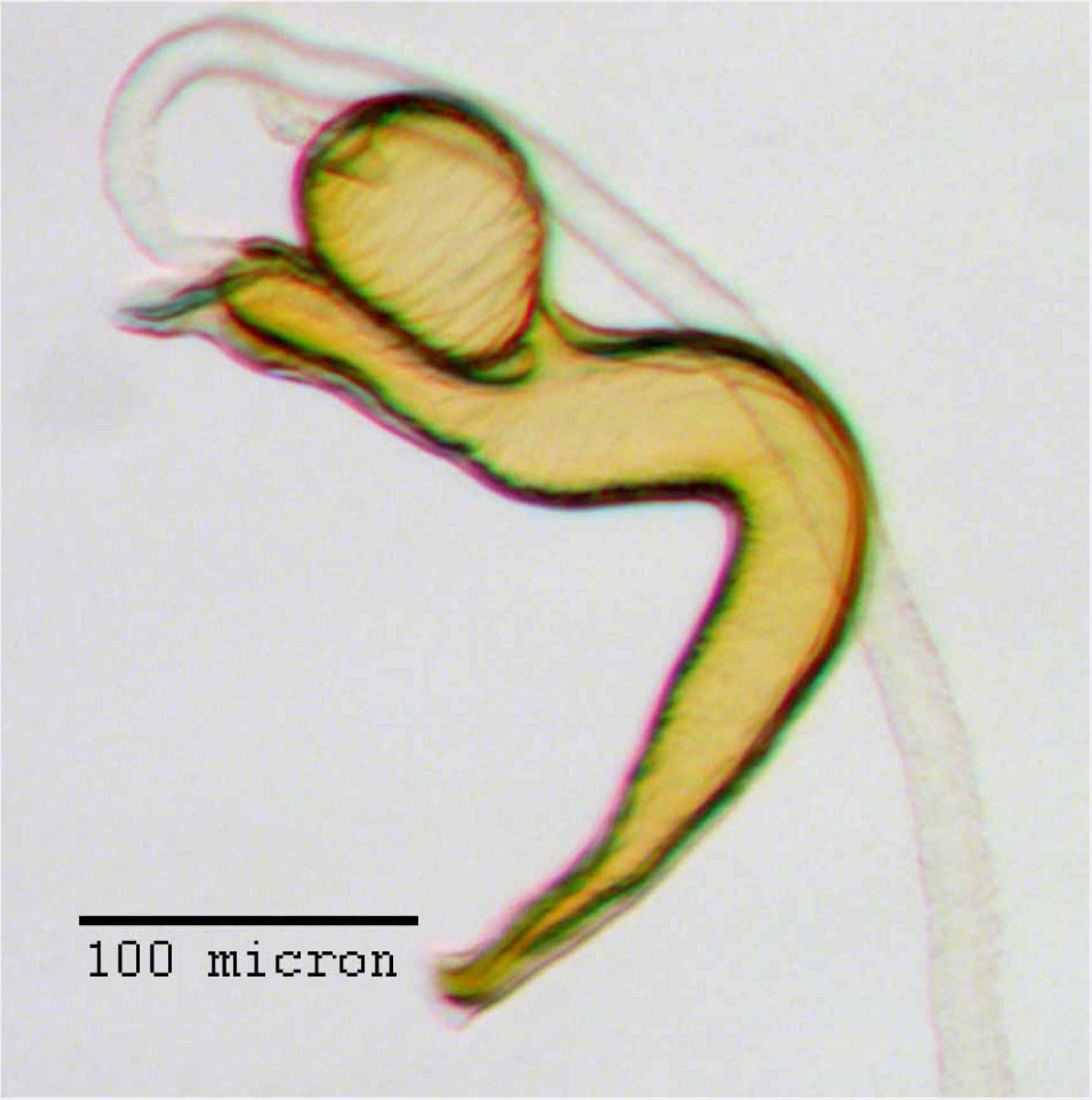
Spermatheca of *M. ampullaceus* [JF2018]. Genitalia of female (♀) holotype.

**Figure 6.**
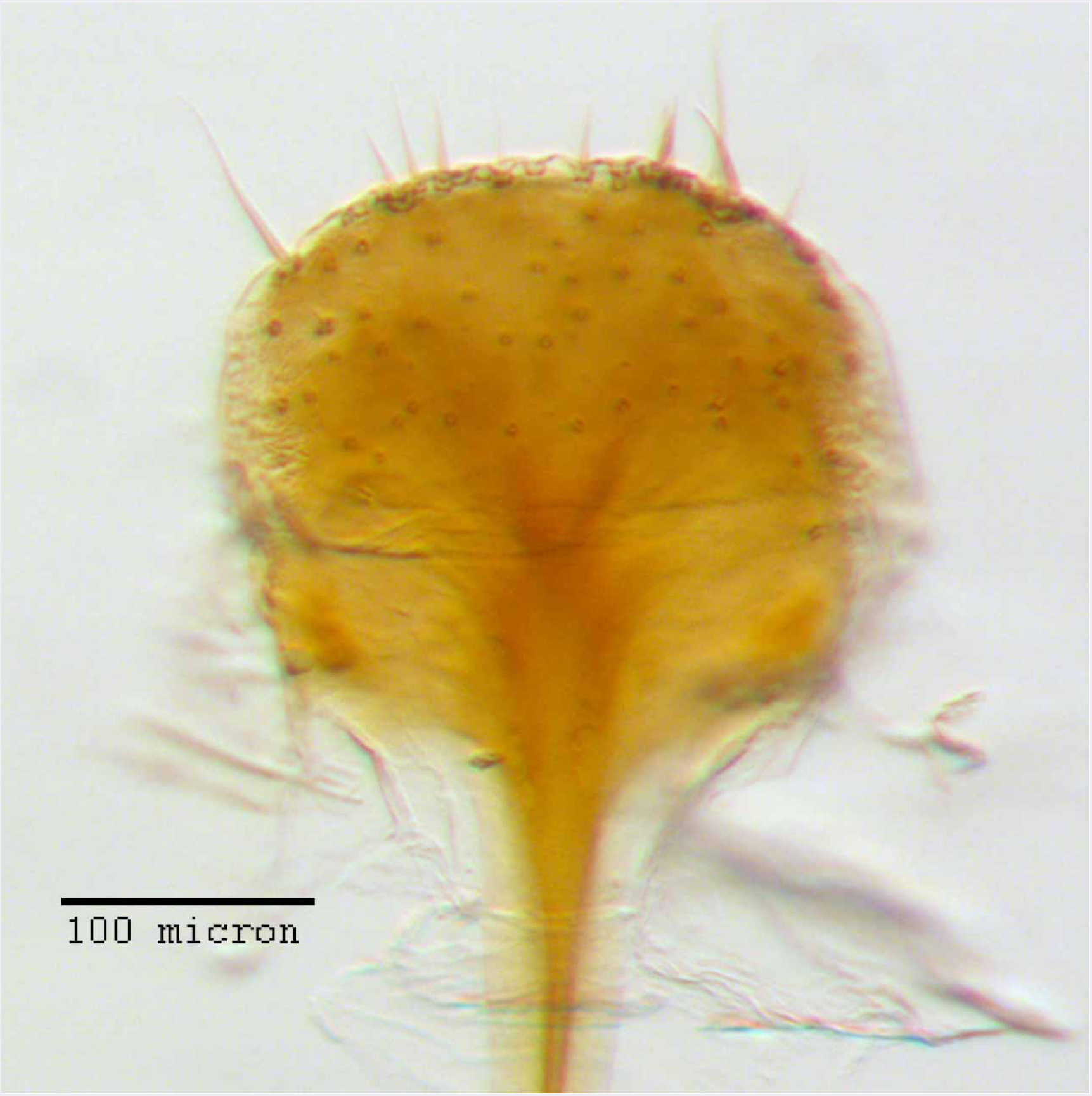
Lamina of spiculum ventrale of *M. ampullaceus* [JF2018]. Sternum VIII of female (♀) holotype.

#### Diagnosis

*Minyomerus ampullaceus* [JF2018] is best differentiated from other congenerics by its unique body shape, which most prominently features a strongly constricted, sub-cylindrical pronotum and greatly protuberant elytra; this combination gives the species a distinctly flask- or bottle-shaped appearance. Due to the relatively poor condition of the scales and setae of the holotype, color and setation cannot be reliably used for identification. However, the elytra themselves are unique in shape, and diagnostic, together nearly 2 × the width of the pronotum at their widest point, and nearly 3/4 × as wide as long in dorsal view. In lateral view the anterior and posterior declivities of the elytra are strongly abrupt, and nearly vertical; most notably, the anterior margin of the elytra projects strongly and characteristically dorsad of its articulation with the posterior pronotal margin. The spermatheca is also quite distinct, having a highly elongate projection of the corpus aligned with midline of the ramus, which is basally tapered and angled at nearly 45° to the corpus.

#### Description of female

##### Habitus

Length 3.76 mm, width 1.76 mm, length/width ratio 2.14, widest at anterior 1/3 of elytra. Integument orange-brown to black. Scales with variously interspersed colors ranging from slightly off-white to beige to yellow. Setae recumbent to sub-recumbent, white to brown in color.

##### Mandibles

Partially covered with white, slightly opalescent scales, with 3 longer setae, and 1 shorter seta between these.

##### Rostrum

Length 0.54 mm, anterior portion 1.5-2× broader than long, rostrum/pronotum length ratio 0.57, rostrum length/width ratio 1.10. Separation of rostrum from head generally obscure. Dorsal outline of rostrum nearly square, anterior half of dorsal surface mesally concave, posterior half coarsely but shallowly punctate to rugose. Rostrum in lateral view nearly square; apical margin broadly bisinuate and emarginate, with 2 pairs of large vibrissae. Nasal plate defined by Y-shaped, impressed lines, convex, integument partially covered with white scales. Margins of mandibular incision directed ca. 15ú outward dorsally in frontal view. Ventrolateral sulci strongly defined, beginning as a narrow sulcus posteriad of insertion point of mandibles, running parallel to scrobe, terminating in a ventral fovea.

##### Antennae

Small tooth formed by overhanging dorsal margin of scrobe directly ventrad of margin of eye. Scape extending to posterior 1/3 of eye. Funicular segments V-VII and club missing.

##### Head

Eyes globular, anterodorsal margin of each eye feebly impressed, posterior margin elevated from lateral surface of head; eyes separated in dorsal view by 4× their anterior-posterior length, set off from anterior prothoracic margin by 1/3 of their anterior-posterior length. Head without any transverse post-ocular impression.

##### Pronotum

Length/width ratio 0.88; widest near midpoint. Anterior margin slightly arcuate, lateral margins curved and widening into a bulge just anteriad of midpoint of pronotum, posterior margin straight, with a slight mesal incurvature. Pronotum in lateral view with setae that reach beyond anterior margin by 1/2 of their length; these setae becoming evenly longer and more erect laterally, reaching a maximum length equal to 1/2 of length of eye. Anterolateral margin with a reduced tuft of 6-7 post-ocular vibrissae present, emerging near ventral 1/2 of eye, and stopping just below ventral margin of eye; vibrissae subequal in length at 1/3 of anterior-posterior length of eye, except for three vibrissae achieving a maximum length similar to anterior-posterior length of eye.

##### Scutellum

Exposed, margins straight.

##### Pleurites

Metepisternum hidden by elytron.

##### Thoracic sterna

Mesocoxal cavities separated by 1/4× width of mesocoxal cavity. Metasternum with transverse sulcus not apparent; metacoxal cavities widely separated by ca. 2× their width.

##### Legs

Profemur/pronotum length ratio 1.04; profemur with distal 1/5 produced ventrally as a rounded projection covering tibial joint; condyle of tibial articulation occupying 4/5 of distal surface and 1/5 length of femur. Protibia/profemur length ratio 0.93; protibial apex with ventral setal comb recessed in an incurved groove; mucro present as a large, black, sub-triangular, medially-projected tooth, which is approximately equilateral and whose sides are sub-equal in length to surrounding setae. Protarsus with tarsomere III 1.25 × as long as II; wider than long. Metatibial apex with almond shaped convexity ringed by 10 short, spiniform setae.

##### Elytra

Length/width ratio 2.66; widest at anterior 1/3; anterior margins jointly almost 2 × wider than posterior margin of pronotum and strongly produced dorsally from margin of pronotum; lateral margins evenly rounded until posterior 1/3, more strongly rounded and converging thereafter. Posterior declivity angled at nearly 85° to main body axis. Elytra with 10 complete striae; striae shallow; punctures faint beneath appressed scales, separated by 5-7 × their diameter; intervals very slightly elevated.

##### Abdominal sterna

Ventrite III anteromesally incurved around a fovea located mesally on anterior margin, posterior margin elevated and set off from IV along lateral 1/3s of its length. Sternum VII mesally 1/2× as long as wide; anterior margin weakly curved.

##### Tergum

Pygidium (tergum VIII) sub-conical; posterior margin emarginate; medial 1/3 of anterior 3/5 of pygidum less sclerotized.

##### Sternum VIII

Anterior laminar edges each incurved forming a 115° angle with lateral margin, this angle distinctly sclerotized; posterior 1/2 of lamina porose throughout, laminar arms more sclerotized medially; posterior edge evenly, moderately arcuate.

##### Ovipositor

Coxites in dorsal view slightly longer than broad, with a medial region that is weakly sclerotized.

##### Spermatheca

Comma-shaped; collum expanded to form a long, cylindrical projection, sub-equal in length to ramus, 1/3 × width of corpus, angled at 45° to corpus, apically with a reduced hood-shaped projection; ramus elongate, bulbous, slightly wider than thickness of corpus, basally constricted to form a short stalk; corpus not greatly swollen; cornu sub-equal in length to corpus and collum, recurved distally to form in inner angle of 60° to corpus, straight and gradually narrowing along basal 2/3, with apical 1/3 abruptly narrowed, angled at 45° to coprus, and tapering to a slight knob.

#### Description of male

Male not available or known.

#### Comments

Due to the limited number of specimens of this species, dissections of mouthparts could not be performed.

#### Etymology

Named in reference to the shape of the body in dorsal view, which appears bottle-shaped due to the large elytra and comparatively cylindrical pronotum – *ampullaceus* = “flasklike”; Latin adjective (Brown 1956).

#### Material examined

##### Holotype

♀ “Carlsbad, N.M.; Geococcyx calif; 144640” (**USNM**).

#### Distribution

This species is known only from Carlsbad, New Mexico (USA), from an unspecified locality; the location of the city is shown in Fig. 36.

#### Natural history

No host plant associations have been documented. The label indicates “Geococcyx calif”; this is presumably a reference to *Geococcyx californianus* (Lesson, 1829) [non-focal] (Cuculidae [non-focal]), the Greater Roadrunner. We had initially believed that this indicated a specimen found in a roadrunner nest; however, according to our reviewers, the USNM frequently assisted with the identification of insect specimens retrieved from the stomach contents of birds, and thus the specimen was most likely retrieved from the gut contents of a roadrunner. This seems quite likely given the poor external condition of the specimen. It is unknown whether this species is parthenogenetic.

### *Minyomerus franko* Jansen & Franz sec. Jansen & Franz, 2018; sp. n

urn:lsid:zoobank.org:act:F8C0153E-DF0E-40E0-AF31-EBEA7075D06D Figures 7-15

**Figure 7.**
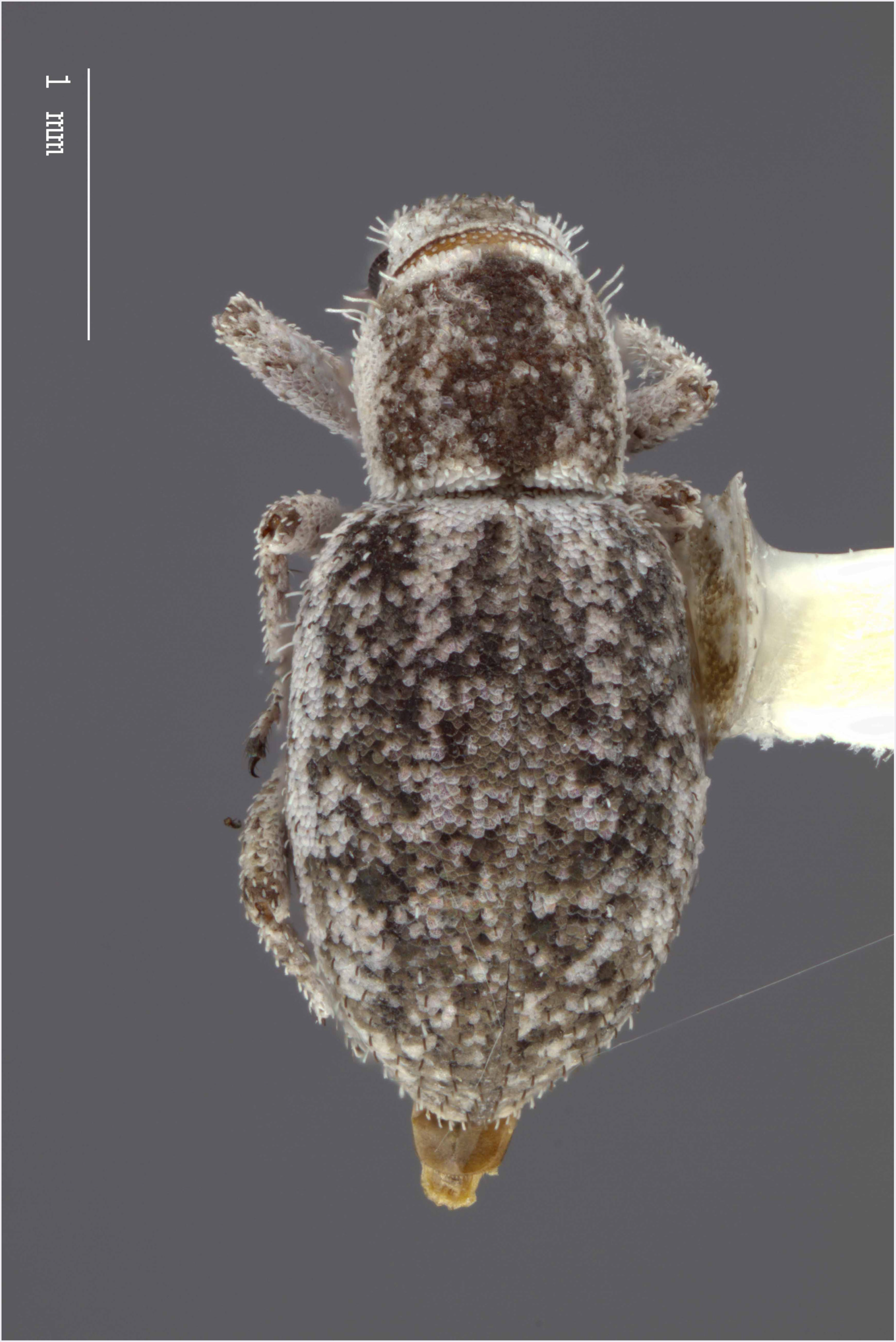
Dorsal habitus of *M. franko* [JF2018]. Image of female (♀) holotype.

**Figure 8.**
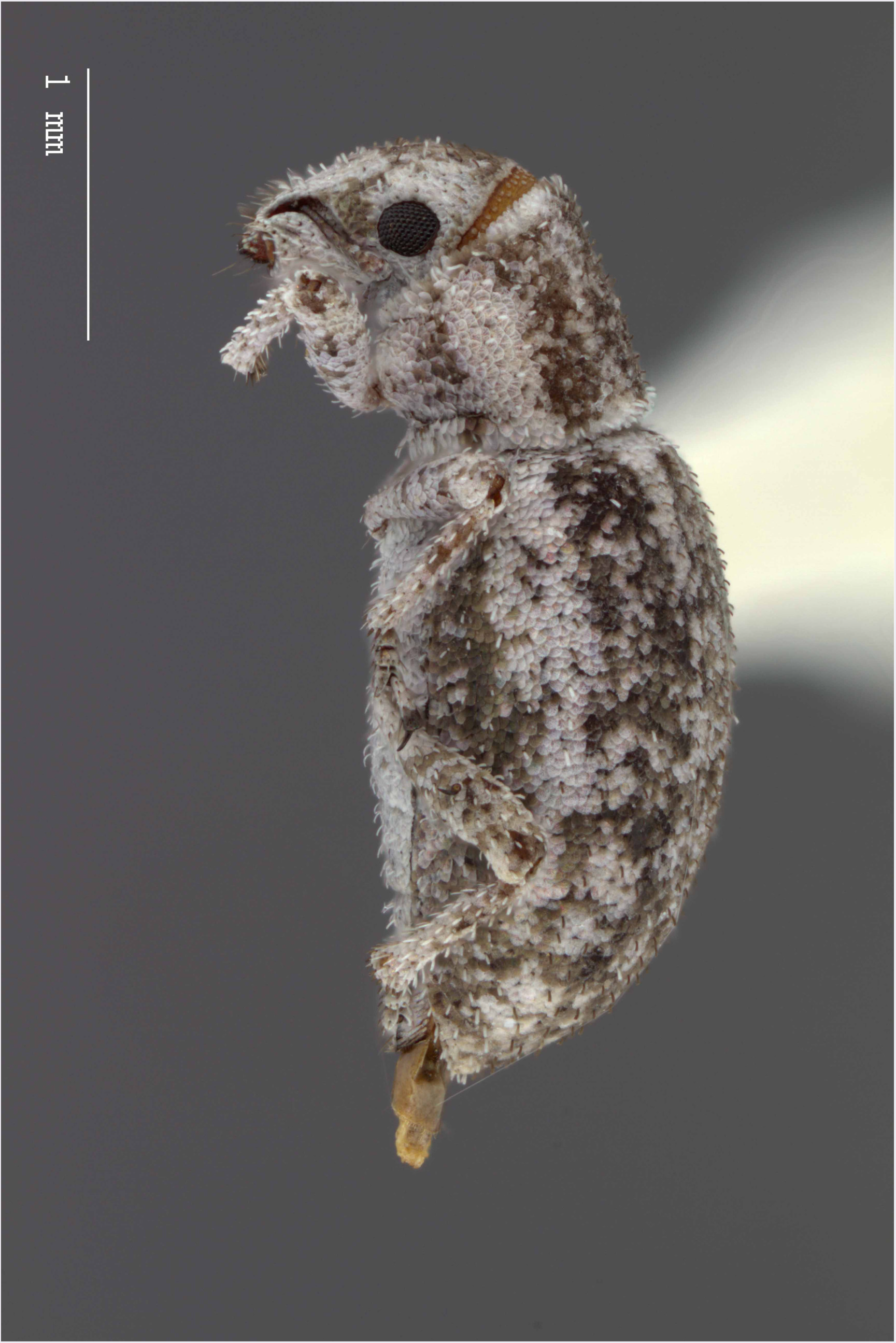
Lateral habitus of *M. franko* [JF2018]. Image of female (♀) holotype.

**Figure 9.**
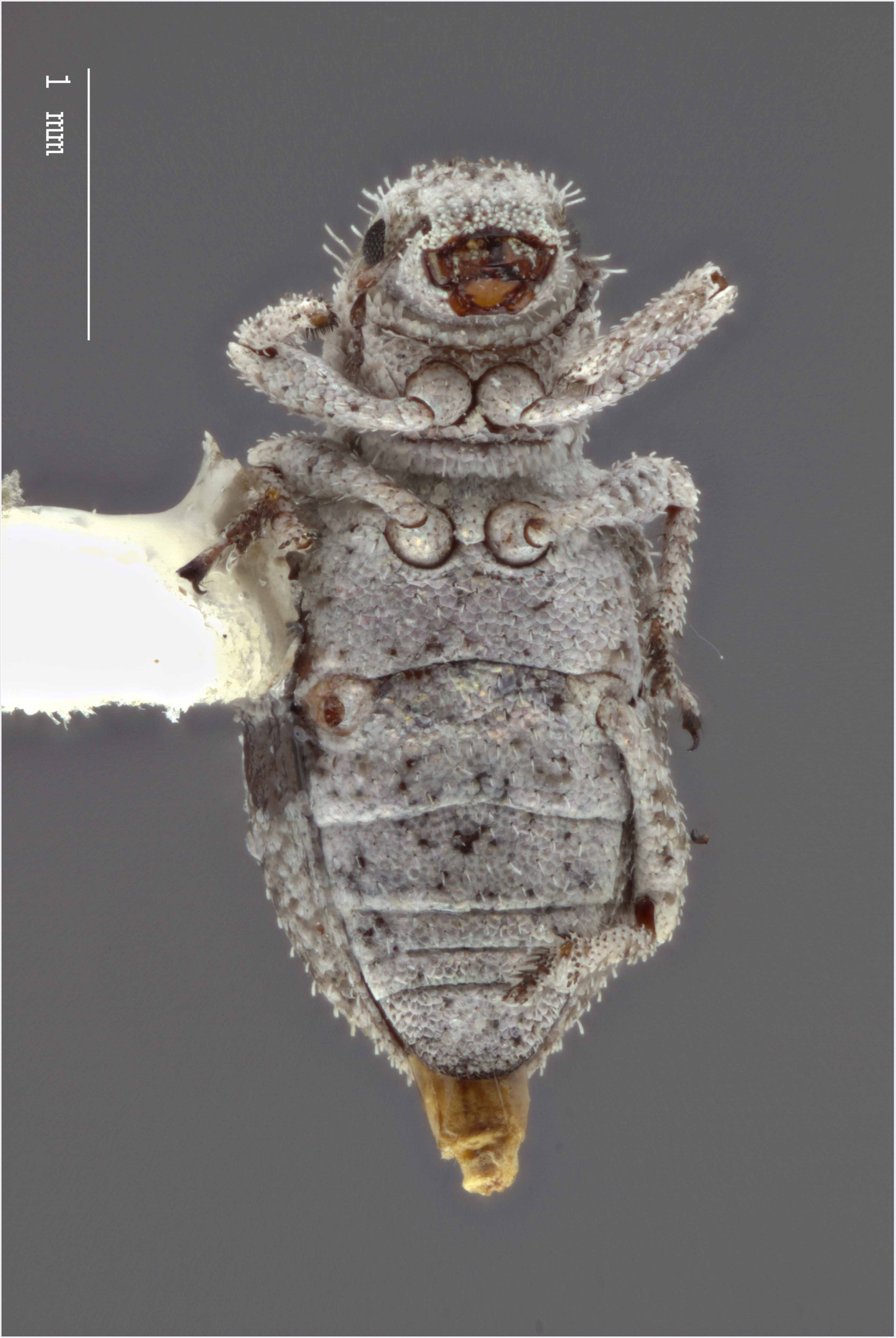
Ventral habitus of *M. franko* [JF2018]. Image of female (♀) holotype.

**Figure 10.**
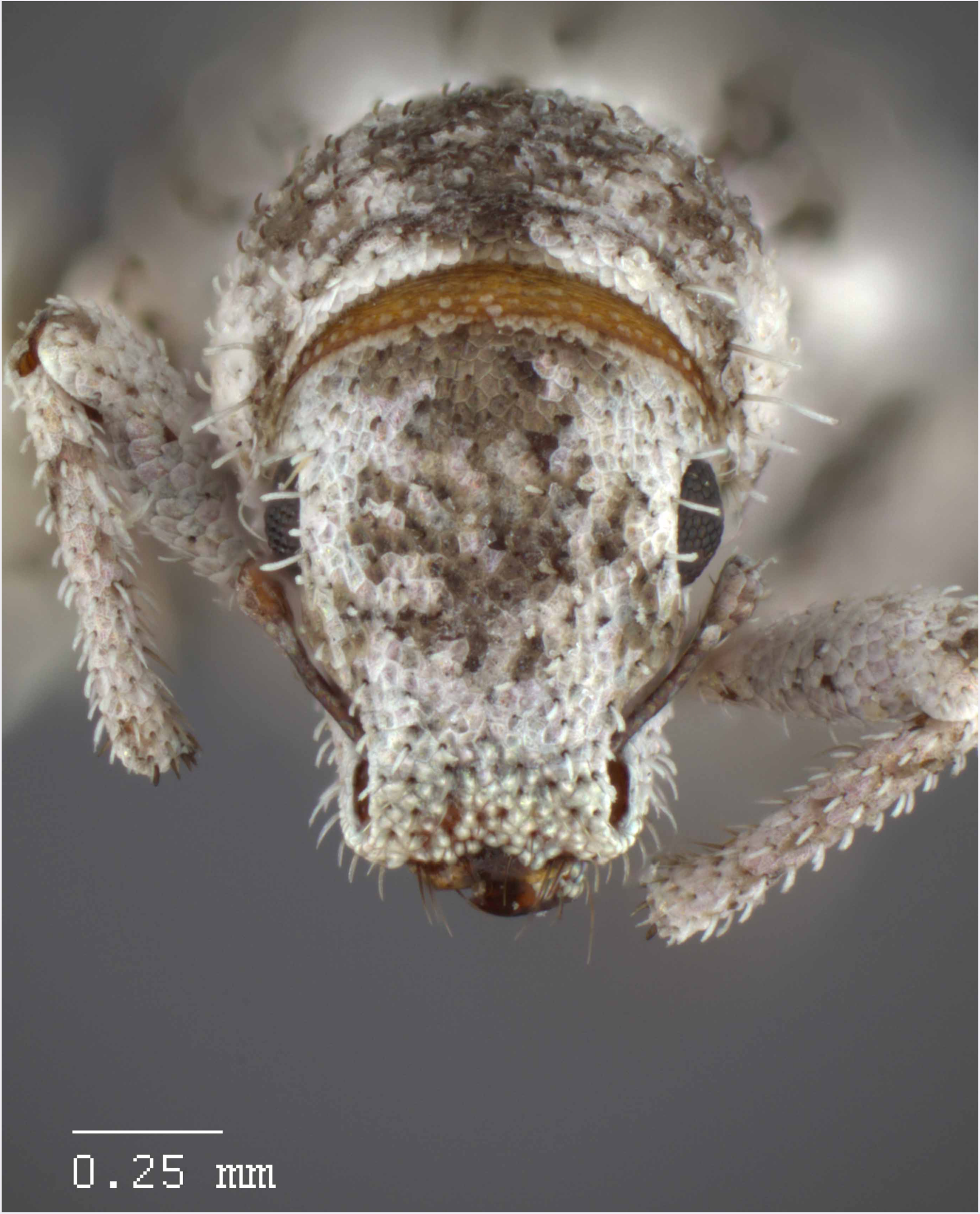
Head and rostrum of *M. franko* [JF2018]. Frontal view of female (♀) holotype.

**Figure 11.**
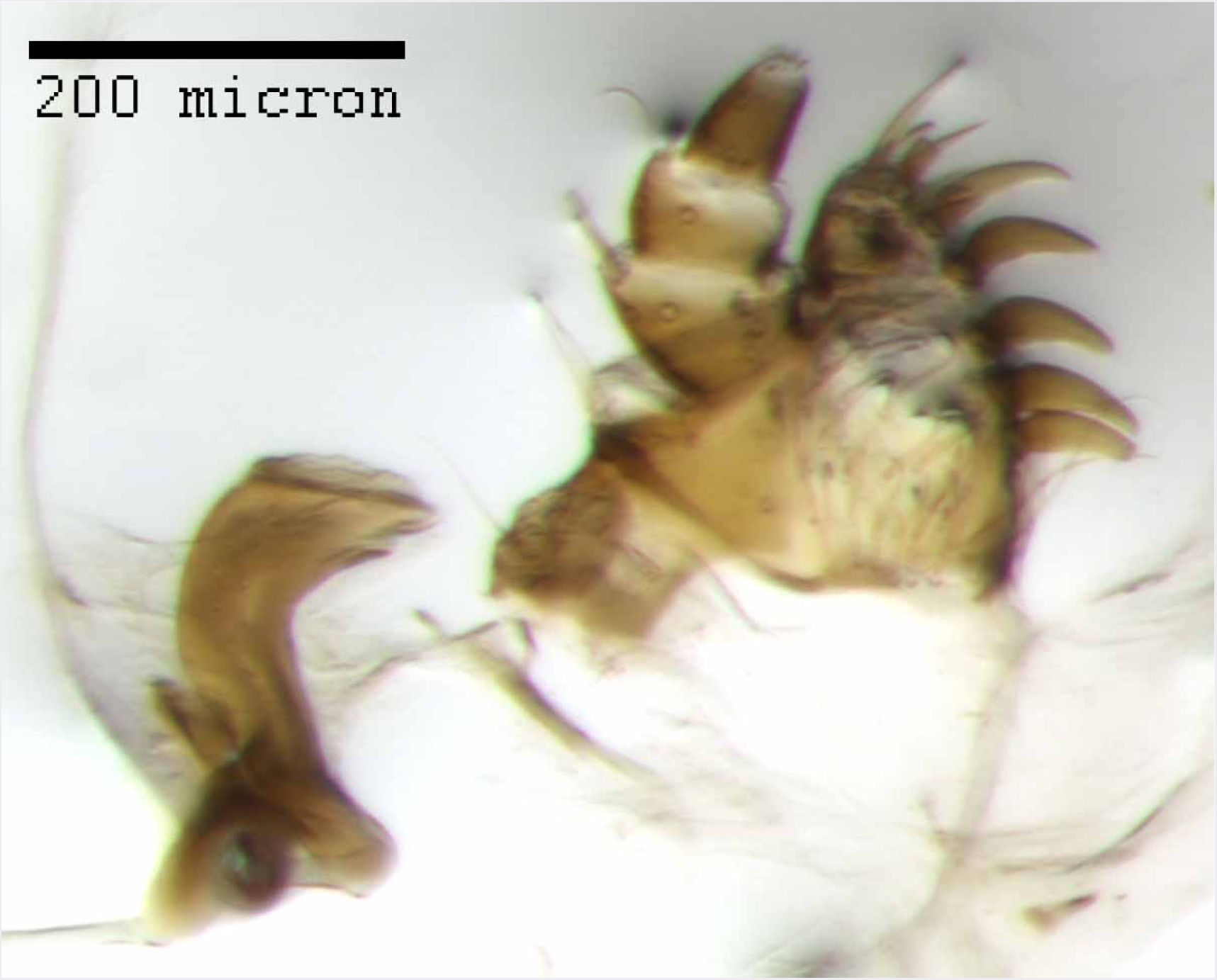
Maxilla of *M. franko* [JF2018]. Dextral maxilla of female (♀) paratype.

**Figure 12.**
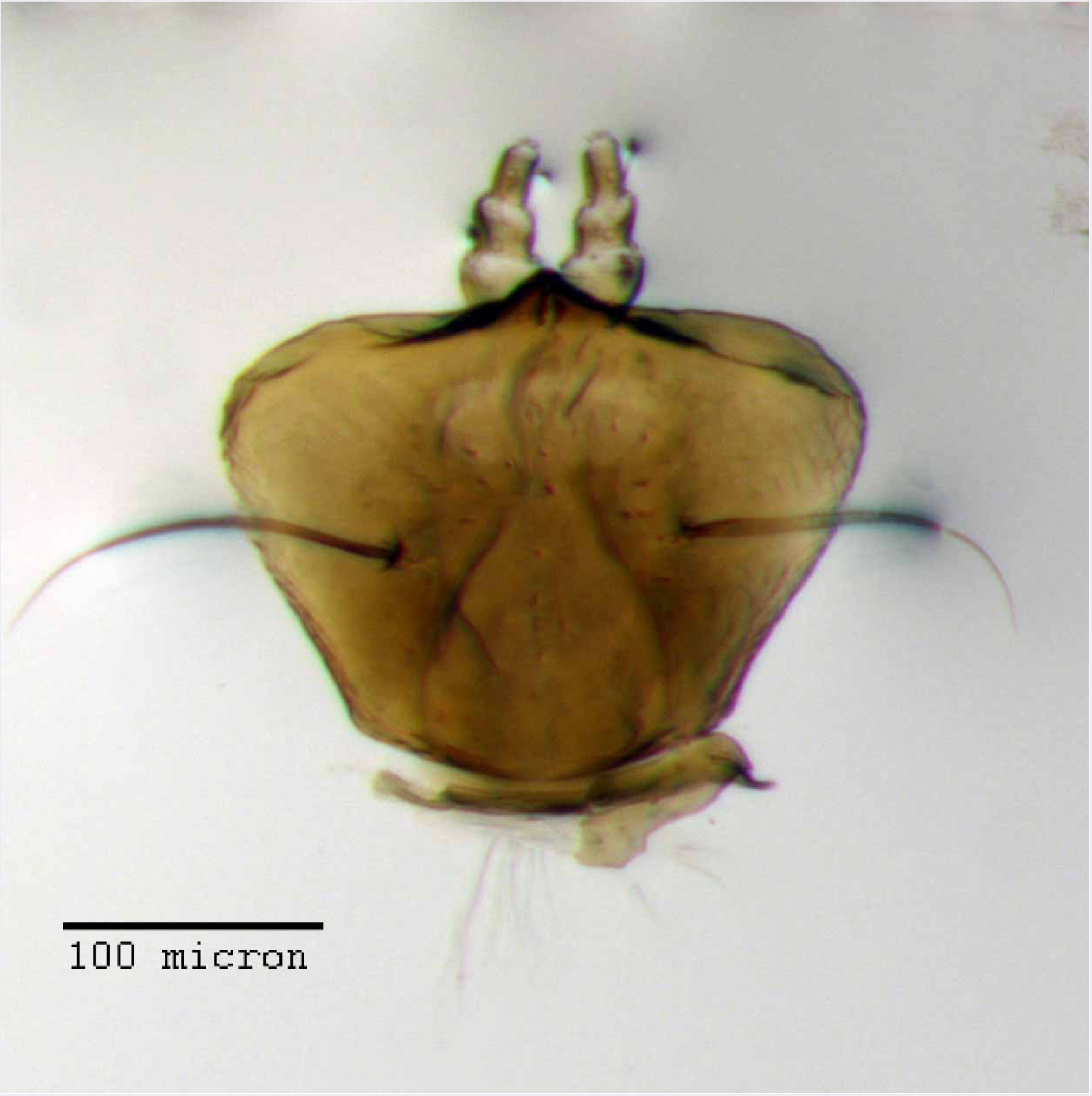
Prementum of *M. franko* [JF2018]. Labium of female (♀) paratype.

**Figure 13.**
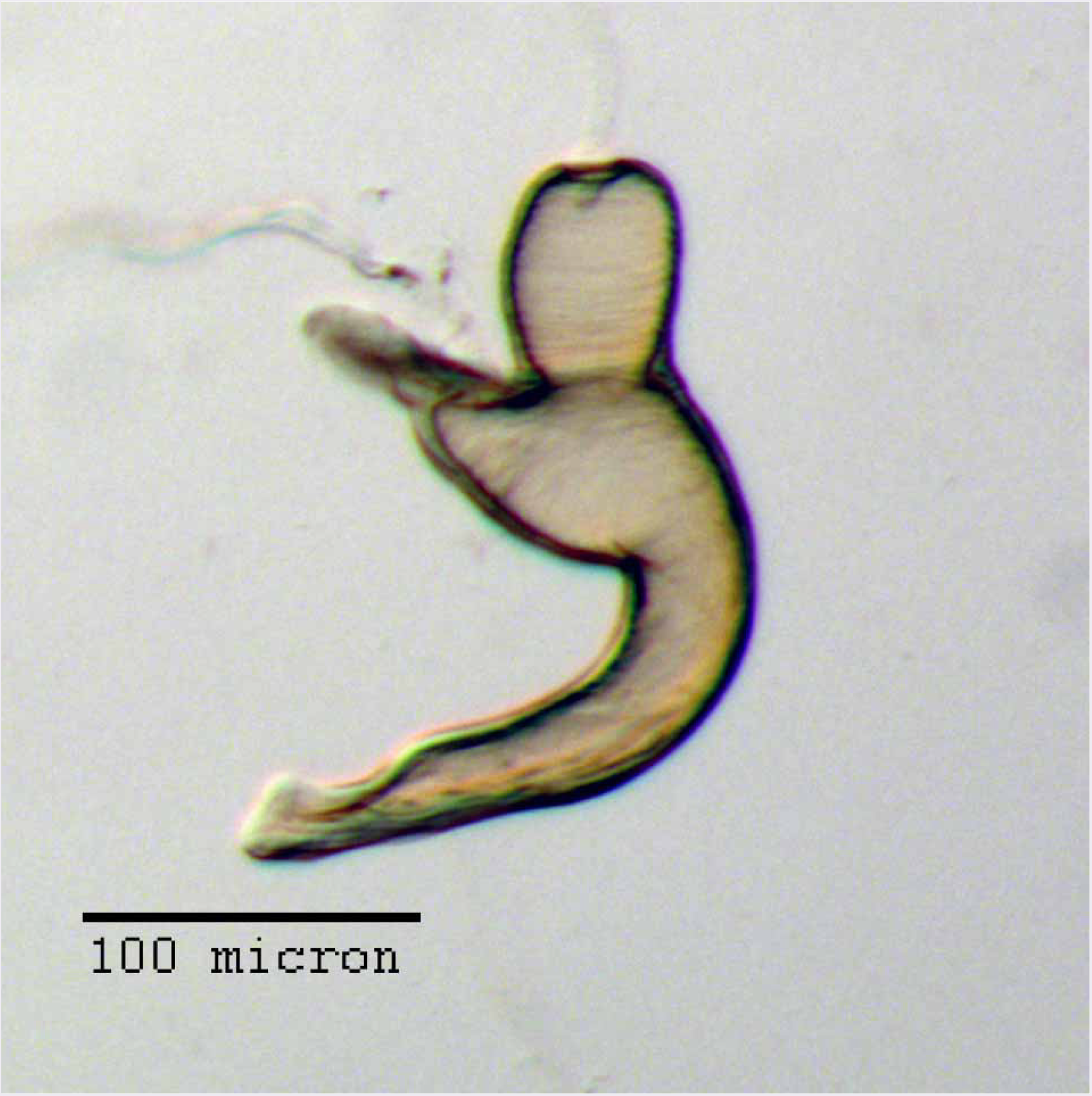
Spermatheca of *M. franko* [JF2018]. Genitalia of female (♀) paratype.

**Figure 14.**
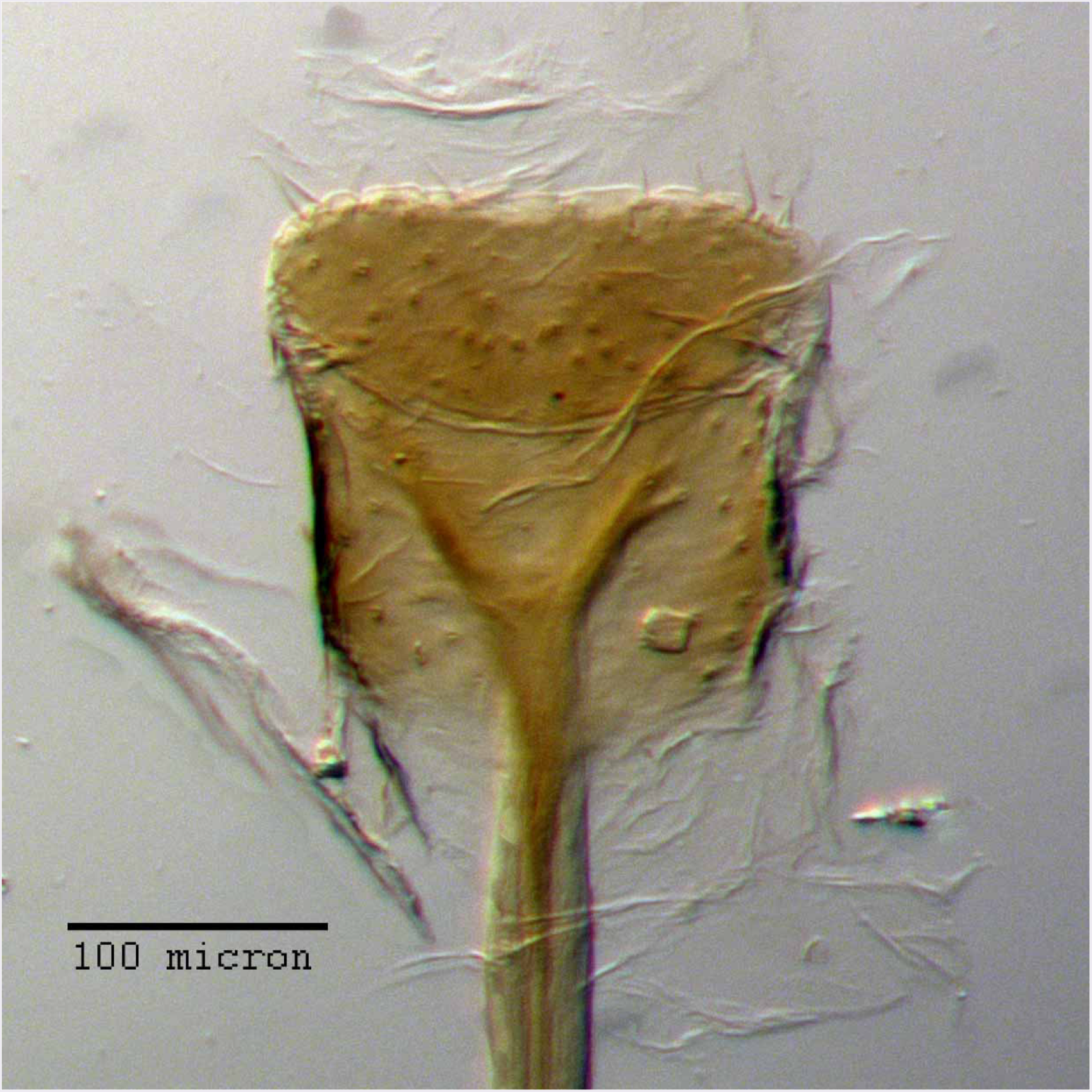
Lamina of spiculum ventrale of *M. franko* [JF2018]. Sternum VIII of female (♀) paratype.

**Figure 15.**
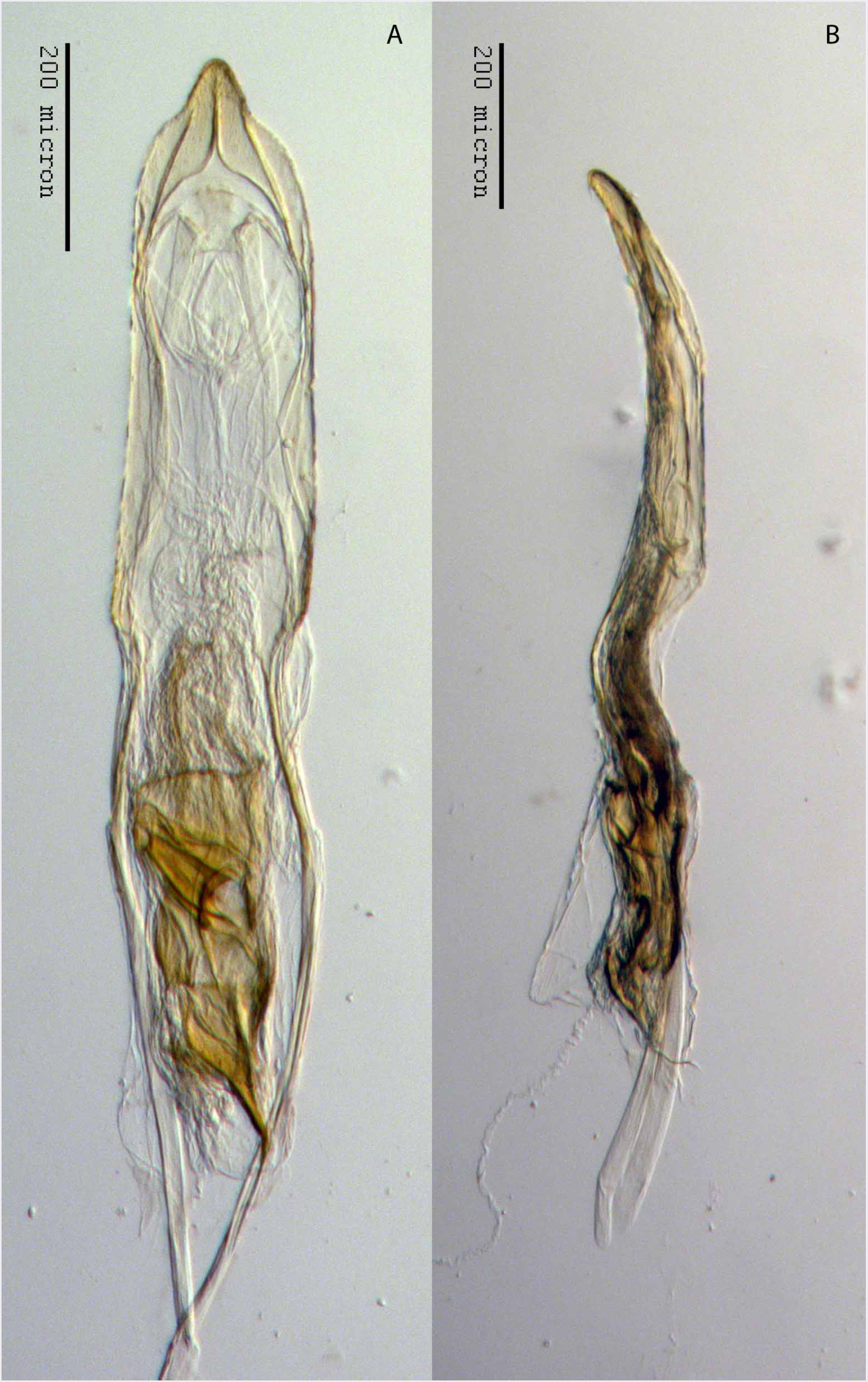
Aedeagus of *M. franko* [JF2018]. Genitalia of male (♂) paratype in (A) dorsal view and (B) lateral view.

#### Diagnosis

*Minyomerus franko* [JF2018] is readily distinguished from other congenerics by the strikingly long setae of the anterior margin of the pronotum, which project laterally up to 80° from the longitudinal axis of the body and achieve a maximum length at least equaling the diameter of the eye. In addition, the setae lining the dorsal margin of the ocular impression are elongate and reach a length equal to 1/2 - 3/4 × the diameter of the eye. The spermatheca has a short, somewhat bulbous corpus, with the ramus sub-equal in size and perpendicular to the corpus, and the collum is strongly recurved along the basal 1/3 of its length. The aedeagus is relatively short and wide, and is abruptly constricted in the apical 1/5 of its length, thereafter tapered to a rounded point.

#### Description of female

##### Habitus

Length 3.10-3.30 mm, width 1.38-1.44 mm, length/width ratio 2.25-2.29, widest at anterior 1/3-1/4 of elytra. Integument orange-brown to black. Scales with variously interspersed colors ranging from slightly off-white or beige to manila/tan to dark coffee brown, in some specimens appearing semi-translucent (in others opaque). Setae linear to slightly apically explanate, appearing minutely spatulate, sub-recumbent to sub-erect, white or brown in color.

##### Mandibles

Covered with white scales, with 3 longer setae, and 1-2 shorter setae between these.

##### Maxillae

Cardo bifurcate at base with an inner angle typically between 90–120°, arms of equal length, inner (mesal) arm nearly 1.5 × thicker than outer arm, both arms of bifurcation equal in length to apically outcurved arm, glabrous. Stipes sub-quadrate, roughly equal in length to each bifurcation of cardo, with a single lateral seta. Galeo-lacinial complex nearly extending to apex of maxillary palpomere II; complex mesally membranous, laterally sclerotized, with sharp demarcation of sclerotized region separating palpiger from galeo-lacinial complex; setose in membranous area just adjacent to sclerotized region, setae covering 2/3 of dorsal surface area; dorsally with 7 apicomesal lacinial teeth; ventrally with 4 reduced lacinial teeth. Palpiger with a single lateral seta, otherwise glabrous and evenly sclerotized throughout.

##### Maxillary palps

I apically oblique, apical end forming a 45° angle with base, with 2 apical setae; II sub-cylindrical, with 1 apical seta.

##### Labium

Prementum roughly trapezoidal; apical margins angulate, ventral margin gently sinuate, dorsal margin straight; lateral margins feebly incurved near posterior margin; basal margin arcuate. Labial palps 3-segmented, I with apical 2/3 projecting beyond margin of prementum, exceeding apex of ligula; III slightly longer than II.

##### Rostrum

Length 0.46-0.48 mm, anterior portion 1.75-2.25× broader than long, rostrum/pronotum length ratio 0.58-0.59, rostrum length/width ratio 1.21-1.26. Separation of rostrum from head generally obscure. Dorsal outline of rostrum sub-rectangular, anterior half of dorsal surface feebly impressed, posterior half coarsely but shallowly punctate to rugose. Rostrum in lateral view nearly square; apical margin bisinuate and emarginate, with 2 large vibrissae. Nasal plate defined by broad, V-shaped, shallowly impressed lines, anteromesally slightly convex, integument partially covered with white scales. Margins of mandibular incision directed ca. 15° outward dorsally in frontal view. Ventrolateral sulci weakly defined (or entirely absent in some specimens) as a broad concavity dorsad of insertion point of mandibles, running parallel to scrobe, becoming flatter posteriorly and disappearing ventrally. Dorsal surface of rostrum with short, linear, median fovea. Rostrum ventrally lacking sulci at corners of oral cavity.

##### Antennae

Small tooth formed by overhanging dorsal margin of scrobe anterior to margin of eye by 1/5 of length of eye. Scape nearly extending to posterior 1/4 of eye. Terminal funicular antennomere lacking appressed scales, having instead a covering of apically-directed pubescence with interspersed sub-erect setae. Club nearly 3 × as long as wide.

##### Head

Eyes globular to slightly elongate, slanted ca. 35° antero-ventrally; eyes separated in dorsal view by 4× their anterior-posterior length, set off from anterior prothoracic margin by 1/3 of their anterior-posterior length. Head without any transverse post-ocular impression.

##### Pronotum

Length/width ratio 0.84-0.86; widest near anterior 1/3, between anterior constriction and midpoint. Anterior margin arcuate, lateral margins curved and widening into a slight bulge just anteriad of midpoint of pronotum, posterior margin straight, with a slight mesal incurvature. Pronotum in lateral view with setae that reach just beyond anterior margin, angled laterally at 45-80° to longitudinal axis, and strikingly long; these setae becoming evenly longer and more angled laterally, reaching a maximum length nearly equal to length of eye. Anterolateral margin with a reduced tuft of 5 post-ocular vibrissae present, emerging near ventral 1/2 of eye, and stopping just below ventral margin of eye; vibrissae sub-equal in length at 1/3× anterior-posterior length of eye, except for one vibrissa achieving a maximum length similar to anterior-posterior length of eye.

##### Scutellum

Narrowly exposed, with visible area approximately equal to length of appressed scales, margins straight.

##### Pleurites

Metepisternum nearly hidden by elytron except for triangular extension.

##### Thoracic sterna

Mesocoxal cavities separated by 1/3 × width of mesocoxal cavity. Metasternum with transverse sulcus not apparent; metacoxal cavities widely separated by ca. 2× their width.

##### Legs

Profemur/pronotum length ratio 1.01-1.02; profemur with distal 1/5 produced ventrally as a sub-rectangular projection covering tibial joint; condyle of tibial articulation occupying 4/5 of distal surface and 1/5 length of femur. Protibia/profemur length ratio 0.86-0.89; protibial apex with ventral setal comb recessed in a subtly incurved groove; mucro present as a large, black, sub-triangular, medially-projected tooth, which is approximately equilateral and whose sides are sub-equal in length to surrounding setae. Protarsus with tarsomere III 2× as long as II; wider than long. Metatibial apex with almond shaped convexity ringed by 8-9 short, spiniform setae.

##### Elytra

Length/width ratio 3.08-3.20; widest at anterior 1/3-1/4; anterior margins jointly 1.5 × wider than posterior margin of pronotum; lateral margins sub-parallel to slightly rounded after anterior 1/3, more strongly rounded and converging in posterior 1/3. Posterior declivity angled at 70-85° to main body axis. Elytra with 10 complete striae; striae shallow; punctures faint beneath appressed scales, separated by 5-7 × their diameter; intervals very slightly elevated.

##### Abdominal sterna

Ventrite III anteromesally incurved around a fovea located mesally on anterior margin, posterior margin elevated and set off from IV along lateral 1/3s of its length. Sternum VII mesally 1/2× as long as wide; setae darkening, lengthening, and becoming more erect in posterior 2/3; anterior margin weakly curved.

##### Tergum

Pygidium (tergum VIII) sub-cylindrical; medial 1/3 of anterior 2/3 of pygidum less sclerotized.

##### Sternum VIII

Anterior laminar edges each incurved forming a 140° angle with lateral margin; slightly less sclerotized medially between arms of bifurcation; posterior edge subtly incurved medially.

##### Ovipositor

Coxites 1.5 × as long as broad, glabrous; styli 1/2× as long as coxites. Genital chamber apically sclerotized.

##### Spermatheca

Comma-shaped; collum short, apically with a large, hood-shaped projection angled at ca. 60° to ramus, nearly equal in length and contiously aligned with curvature of bulb of ramus; collum sub-contiguous with, and angled at 90° to ramus; ramus elongate, sub-cylindrical to slightly bulbous, 4/5 × thickness of corpus; corpus swollen, 1.25 × thicknes of ramus and 1.5 × thickness of cornu; cornu elongate, strongly recurved in basal 1/3, nearly straight thereafter and narrowing apically, abruptly narrowed in apical 1/3 with apex angled at 30° to corpus.

#### Description of male

Similar to female, except where noted.

##### Habitus

Length 2.47-2.81 mm, width 0.99-1.24 mm, length/width ratio 2.27-2.49. Rostrum length 0.30-0.42 mm, rostrum/pronotum length ratio 0.44-0.53, rostrum length/width ratio 1.00-1.08. Pronotum length/width ratio 0.91-1.00. Profemur/pronotum length ratio 0.87-0.90, protibia/profemur length ratio 0.87-0.97. Elytra length/width ratio 3.00-3.10.

##### Elytra

Elytral declivity more angulate than female on average, forming an 80° angle to main body axis, but otherwise as in female.

##### Abdominal sterna

Sternum VII 2/5-1/2× as long as wide, posterior margin arcuate mesally.

##### Tergum

Pygidium (tergum VIII) with posterior 1/3 punctate; anterior 2/3 rugose.

##### Sternum IX

Spiculum gastrale 2× length of aedeagal pedon. Laminar alae located on lateral 1/4 of posterior margin.

##### Aedeagus

Length/width ratio 2.78-3.16; lateral margins very slightly converging posteriorly, abruptly constricted and more strongly converging in apical 1/5. Pedon in lateral view becoming gradually narrower posteriorly in anterior 1/2, ventral margins in posterior 1/2 abruptly curving to meet dorsal margins at a rounded apical point. Flagellum with large, elonage, tortuous apical sclerite, sclerite nearly as long as pedon, with complex, asymmetrical interior structure.

#### Etymology

Named in reference to the long, somewhat unkempt, erect setae on the anterior margin of the pronotum–*franko* = “free”; Old High-German adjective (Brown 1956).

#### Material examined

##### Holotype

♀ “MEX: S.L.P 1 km N.; Entronque El Huizache; 1493 m 2.VI.87; R. Anderson, <u>Sphaeralcea; hastula</u> A. Gray” [non-focal] (**CMNC**).

##### Paratypes

Same label information as female holotype (**CMNC**: 1 ♀, 1 ♂; **TAMU**: 2 ♂); “MEXICO: S.L.P; 19.6 mi. n. Huizache; July 25, 1976; Peigler, Gruetzmacher,; R&M Murray, Schaffner” (**CMNC**: 1 ♂); “MEXICO: San Luis Potosi; Entronque el Hulzache; 2 June 1987; R. Turnbow” (**USNM**: 1 ♀; **CMNC**: 1 ♂); “MEXICO: Tamaulipas; 8.8 mi. ne. Jaumave; October 10, 1973; Gaumer & Clark” (TAMU: 2♀); “9 mi east Santo; Domingo, S.L.P.,; Mexico XI-14-68; Veryl V. Board” (TAMU: 2 ♂).

#### Distribution

This species has been found in San Luis Potosí and Tamaulipas (Mexico). It is likely to be found throughout the Chihuahuan Desert and arid regions of south-central Mexico based on habitat similarity (Fig. 37).

#### Natural history

Associated with spear globemallow *Sphaeralcea hastulata* A. Gray [non-focal] (Malvaceae [non-focal]). The indication of “Sphaeralcea hastula A. Gray” is not a valid name and appears to be a misspelling of *Sphaeralcea hastulata* [non-focal].

### *Minyomerus sculptilis* Jansen & Franz sec. Jansen & Franz, 2018; sp. n

urn:lsid:zoobank.org:act:EA0B1AD9-68F2-4409-A0F8-903B0DA0FFF9 Figures 16-22

**Figure 16.**
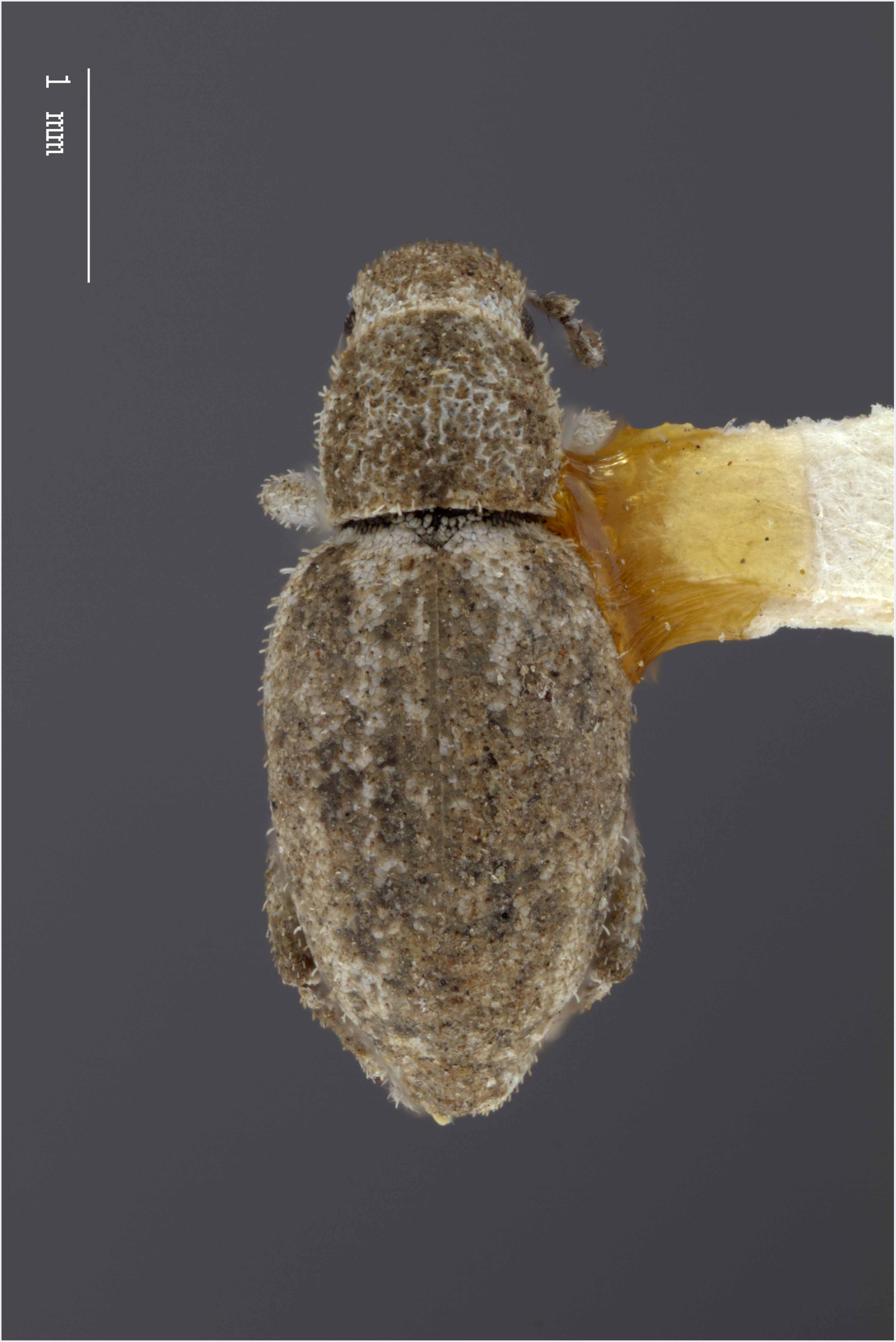
Dorsal habitus of *M. sculptilis* [JF2018]. Image of female (♀) holotype.

**Figure 17.**
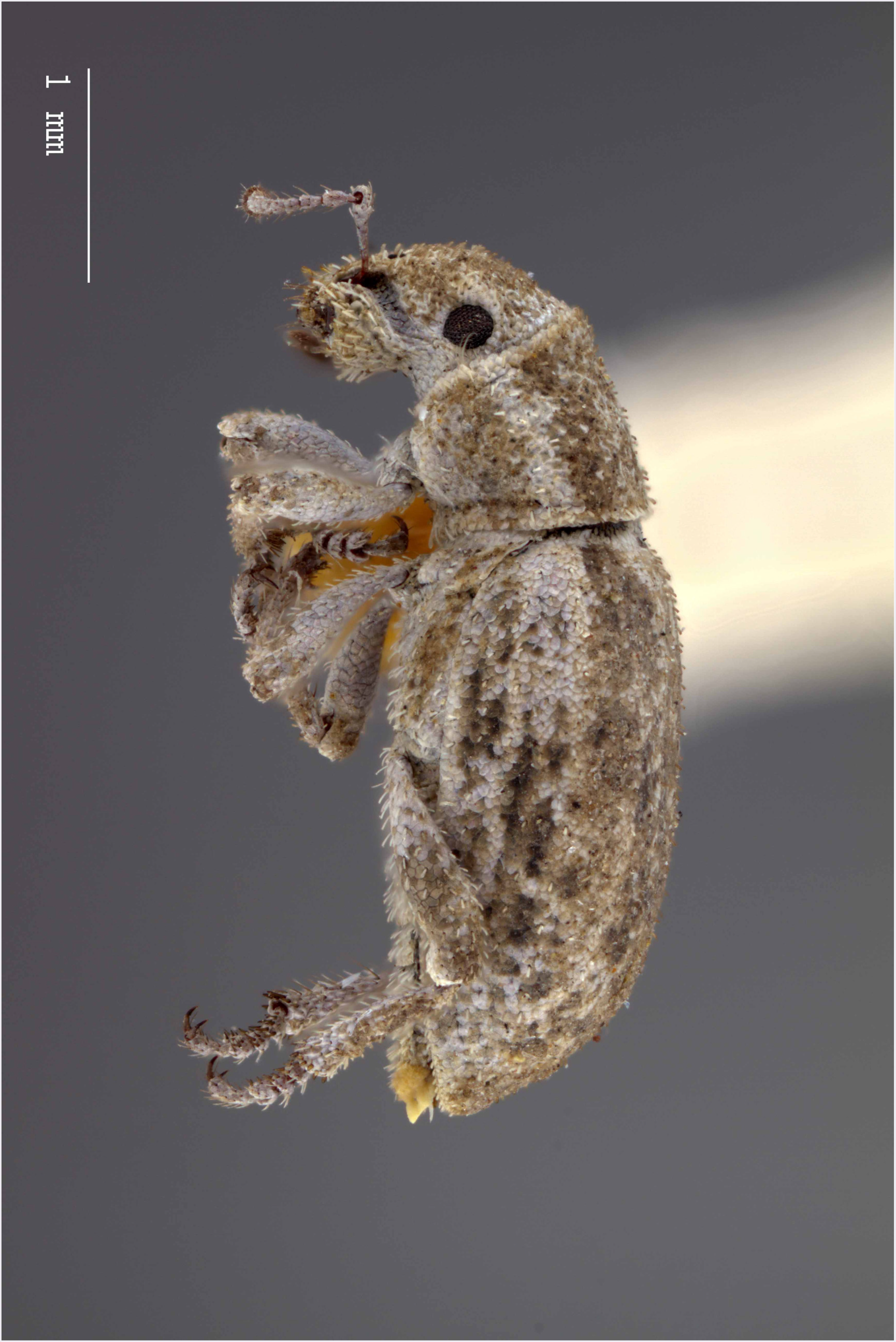
Lateral habitus of *M. sculptilis* [JF2018]. Image of female (♀) holotype.

**Figure 18.**
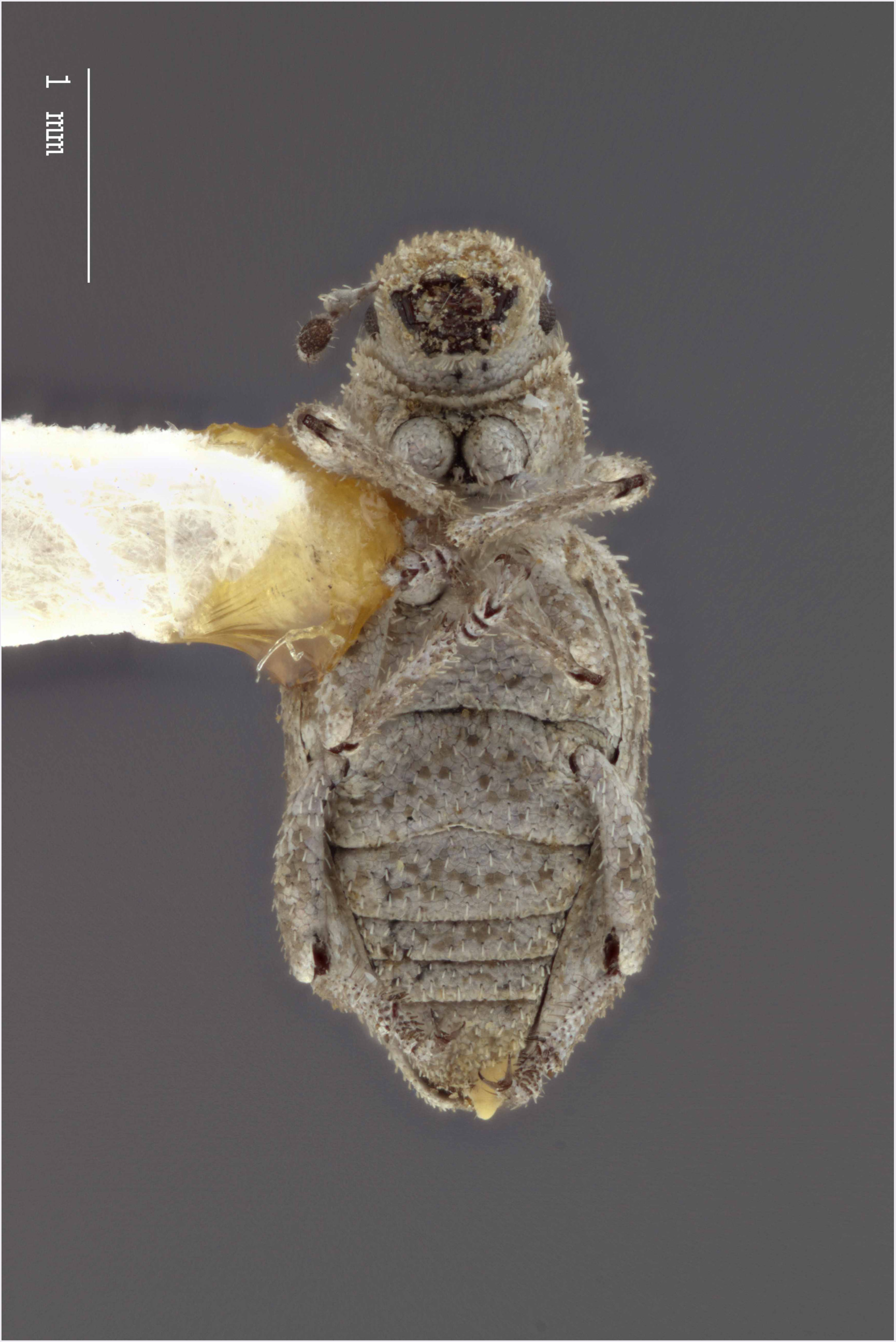
Ventral habitus of *M. sculptilis* [JF2018]. Image of female (♀) holotype.

**Figure 19.**
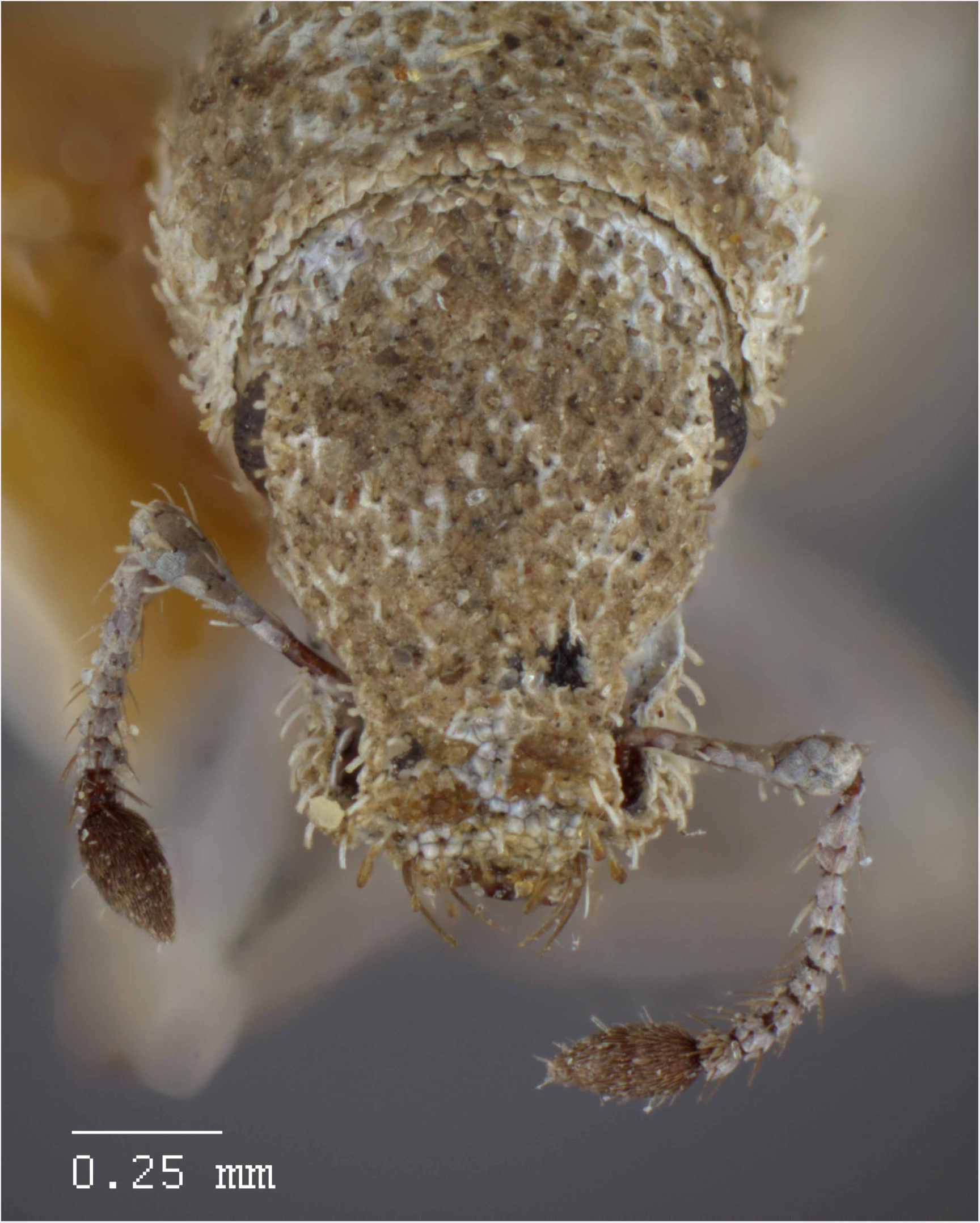
Head and rostrum of *M. sculptilis* [JF2018]. Frontal view of female (♀) holotype.

**Figure 20.**
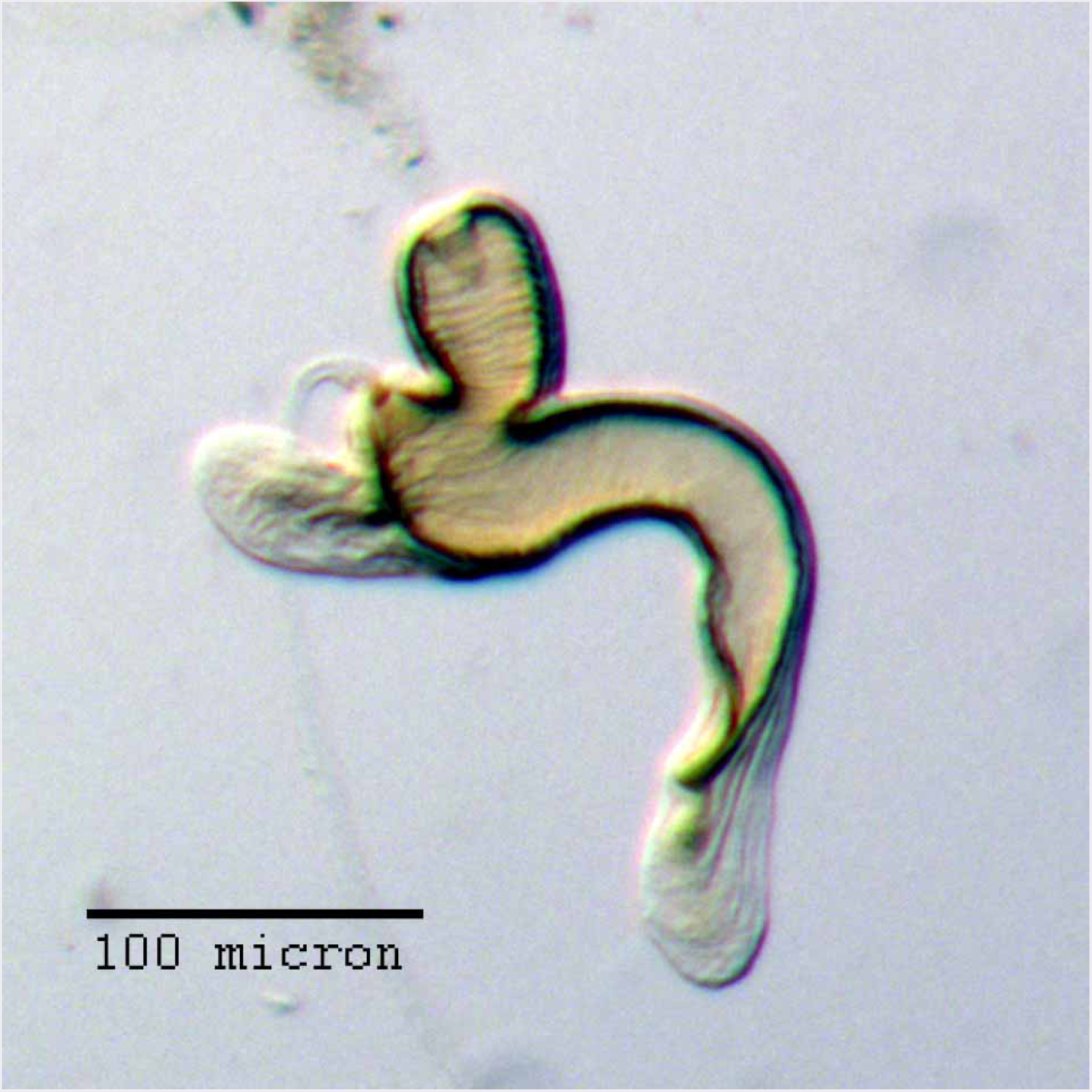
Spermatheca of *M. sculptilis* [JF2018]. Genitalia of female (♀) paratype.

**Figure 21.**
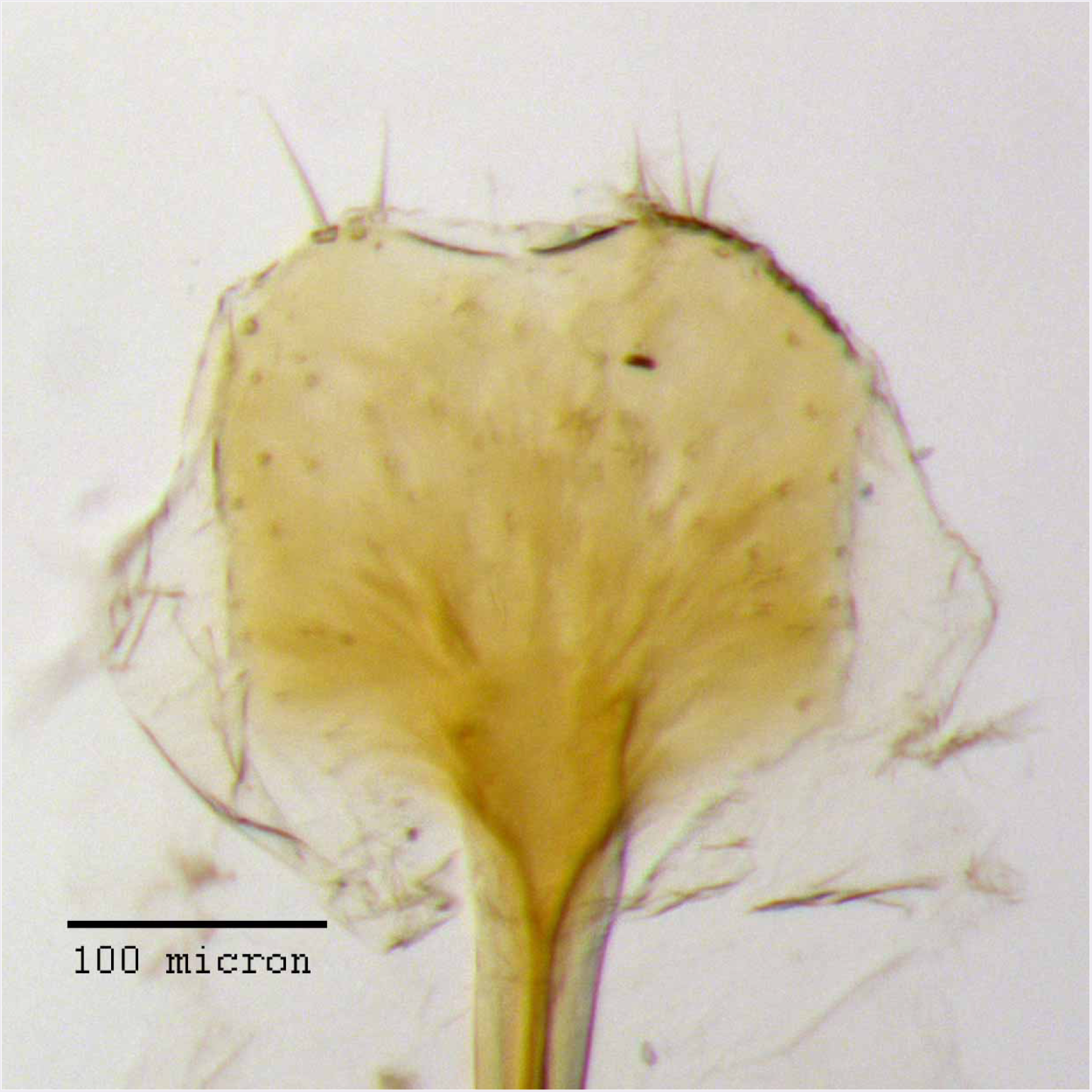
Lamina of spiculum ventrale of *M. sculptilis* [JF2018]. Sternum VIII of female (♀) paratype.

**Figure 22.**
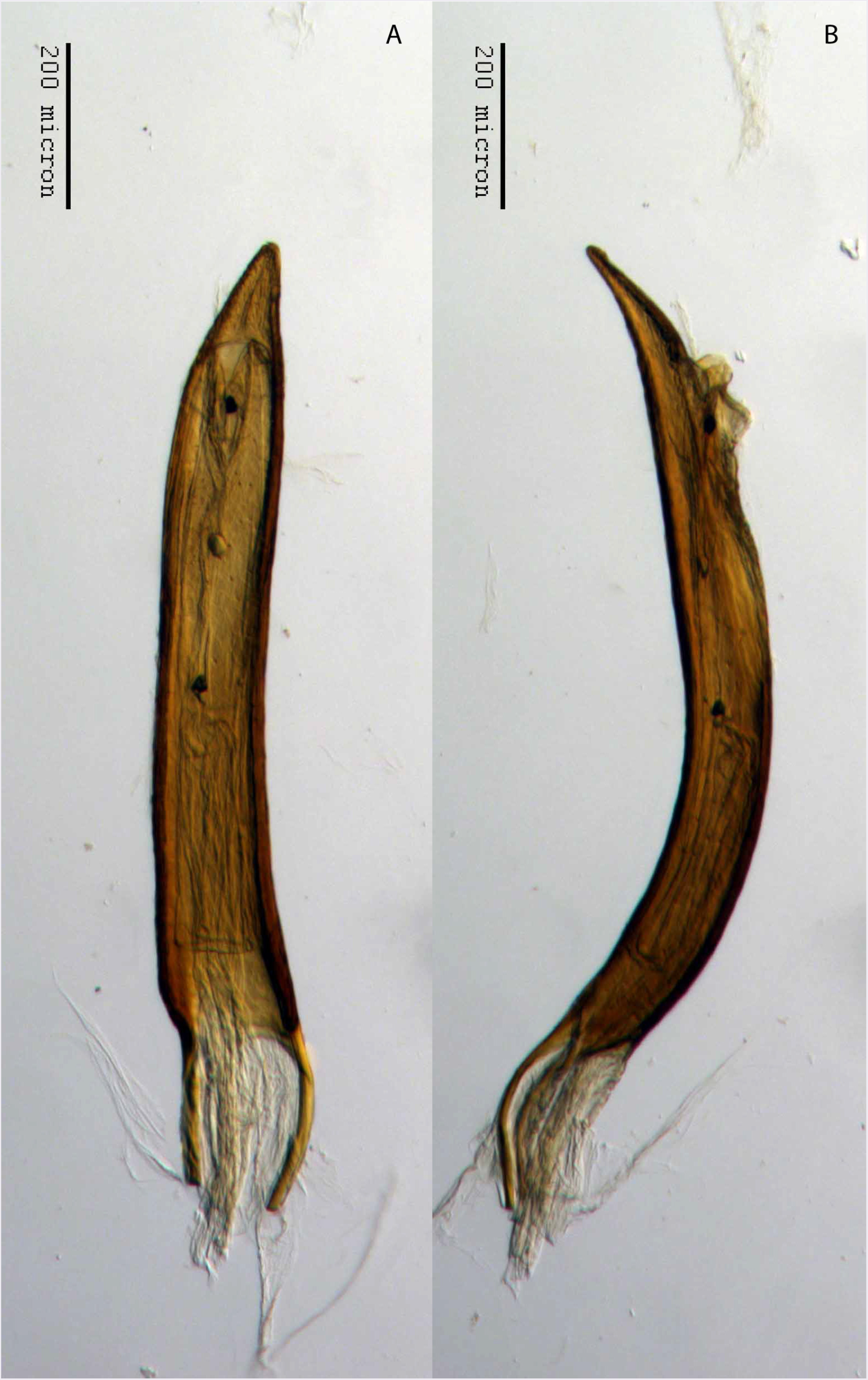
Aedeagus of *M. sculptilis* [JF2018]. Genitalia of male (♂) paratype in (A) dorsal view and (B) lateral view.

#### Diagnosis

*Minyomerus sculptilis* [JF2018] is best distinguished from other congenerics, especially *Minyomerus imberbus* Jansen & Franz, 2015 [JF2015], by a combination of characters, as follows. The interspersed setae on the body are linear and either brown or white. The anterior margin of the pronotum bears a reduced tuft of post-ocular vibrissae. The head is barely elevated between the eyes. The ventrolateral sulci of the rostrum are well defined. The lateral face of each elytron has the intervals raised and well sculpted in appearance. The spermatheca is distinct and has an elongate, annulate, basally tapered ramus, which is slightly thinner than corpus. The cornu is strongly recurved in the basal half, giving it a uniquely sinuate appearance. Both the corpus and cornu terminate in large, hood-shaped, explanate projections equal in size to the ramus. The aedeagus is elongate, acutely angulate, and narrowing towards the apex more strongly in the region of the ostium.

#### Description of female

##### Habitus

Length 3.39-3.70 mm, width 1.33-1.58 mm, length/width ratio 2.34-2.55, widest at anterior 1/5 of elytra. Integument orange-brown to black. Scales with variously interspersed colors ranging from slightly off-white or beige to golden brown to dark coffee brown. Setae sub-recumbent to sub-erect, white to brown in color.

##### Mandibles

Covered with white scales, with 3 longer setae, and 1 shorter seta between these.

##### Rostrum

Length 0.50-0.59 mm, anterior portion ca. 1.5 × broader than long, rostrum/pronotum length ratio 0.66-0.67, rostrum length/width ratio 1.43-1.48. Separation of rostrum from head generally obscure. Dorsal outline of rostrum nearly square, anterior half of dorsal surface mesally concave, posterior half coarsely but shallowly punctate to rugose. Rostrum in lateral view nearly square; apical margin bisinuate and emarginate, with 2 pairs of large vibrissae. Nasal plate defined by Y-shaped, impressed lines, convex, integument covered with white scales. Margins of mandibular incision directed ca. 15-20° outward dorsally in frontal view. Ventrolateral sulci strongly defined, beginning as a narrow sulcus posteriad of insertion point of mandibles, running parallel to scrobe, terminating in a ventral fovea.

##### Antennae

Dorsal margin of scrobe overhanging broadly (not forming a minute tooth). Funicle slightly longer than scape. Scape extending to posterior 1/4 of eye. Club nearly 3 × as long as wide.

##### Head

Eyes globular, anterodorsal margin of each eye impressed, posterior margin slightly elevated from lateral surface of head; eyes separated in dorsal view by 5× their anterior-posterior length, set off from anterior prothoracic margin by 1/4 of their anterior-posterior length. Head between eyes rugose and slightly bulging.

##### Pronotum

Length/width ratio 0.85-0.87; widest near anterior 2/5. Anterior margin arcuate, subtly incurved mesally, and somewhat produced dorsally; anterior constriction broad, posterior margin slightly arcuate. Pronotum in lateral view with setae that reach beyond anterior margin; these setae becoming slightly longer and more erect laterally. Anterolateral margin with a reduced tuft of 3-6 post-ocular vibrissae present, emerging near ventral 1/2 of eye, and stopping just below ventral margin of eye; vibrissae varying in length from 1/2 × anterior-posterior length of eye to a maximum length similar to anterior-posterior length of eye.

##### Scutellum

Exposed, margins straight.

##### Pleurites

Metepisternum nearly hidden by elytron except for triangular extension.

##### Thoracic sterna

Mesocoxal cavities separated by 1/3 × width of mesocoxal cavity. Metasternum with transverse sulcus not apparent; metacoxal cavities widely separated by ca. 2× their width.

##### Legs

Profemur/pronotum length ratio 0.92-1.03; profemur with distal 1/6 produced ventrally as a slightly rounded, sub-rectangular projection covering tibial joint; condyle of tibial articulation occupying 4/5 of distal surface and 1/6 length of femur. Protibia/profemur length ratio 0.87-0.93; protibial apex with ventral setal comb recessed in a subtly incurved groove; mucro not apparent. Protarsus with tarsomere III 1.5 × as long as II; wider than long. Metatibial apex with almond shaped convexity ringed by 10-12 short, spiniform setae.

##### Elytra

Length/width ratio 3.12-3.16; widest at anterior 1/5; anterior margins jointly 1.5-2× wider than posterior margin of pronotum; lateral margins gently converging after anterior 1/5, more strongly converging in posterior 1/4. Posterior declivity angled at 65-70° to main body axis. Elytra with 10 complete striae; striae broadly sculpted; punctures faint beneath appressed scales, separated by 5-7 × their diameter; intervals elevated, with every second interval, beginning at elytral suture, more strongly raised than adjacent intervals.

##### Abdominal sterna

Ventrite III anteromesally incurved around a fovea located mesally on anterior margin, posterior margin elevated and set off from IV along lateral 1/3s of its length. Sternum VII mesally 2/3 × as long as wide; anterior margin straight.

##### Tergum

Pygidium sub-cylindrical; medial 1/2 of anterior 3/5 of pygidium less sclerotized.

##### Sternum VIII

Anterior laminar edges of spiculum ventrale each incurved forming a 125° angle with lateral margin; lamina more sclerotized medially; posterior margin medially incurved.

##### Ovipositor

Coxites as long as broad; styli as long as coxites, glabrous.

##### Spermatheca

S-shaped; collum short, apically with a large, hood-shaped projection roughly aligned with central axis of corpus, nearly equal in length to bulb of ramus; collum sub-contiguous with, and angled at 30° to ramus; ramus elongate, sub-cylindrical to slightly bulbous, 3/4× thickness of corpus, with a short stalk oriented at ca. 45° to the corpus; corpus swollen, 1.3 × thicknes of ramus; cornu short, 2.5-3 × length or ramus, recurved and strongly arched in basal 1/2, forming an inner angle of ca. 80°, feebly sinuate thereafter, with apical 1/2 expanded, then abruptly constricted near apical 1/4 to a fine point.

#### Description of male

Similar to female, except where noted.

##### Habitus

Length 3.10 mm, width 1.22 mm, length/width ratio 2.54. Rostrum length 0.53 mm, rostrum/pronotum length ratio 0.65, rostrum length/width ratio 1.66. Pronotum length/width ratio 0.99. Profemur/pronotum length ratio 1.01, protibia/profemur length ratio 0.82. Elytra length/width ratio 3.18.

##### Elytra

Elytral declivity slightly less angulate than female, forming a 60° angle to main body axis, but otherwise as in female.

##### Abdominal sterna

Sternum VII 1/2× as long as wide, posterior margin feebly arcuate mesally.

##### Tergum

Pygidium (tergum VIII) with mesal 1/3 of posterior margin subtly incurved; posterior 2/3 punctate; anterior 1/3 rugose.

##### Sternum VIII

Consisting of 2 sub-triangular sclerites; antero-laterally with a sharply-pointed projection as long as anterior-posterior length of triangular portion of sclerite.

##### Aedeagus

Length/width ratio 7.00; lateral margins parallel, more strongly converging in region of ostium. In lateral view, width of pedon even throughout in anterior 2/3, ventral margins in posterior 1/3 becoming straight towards apex, then curving to meet dorsal margins at a sharp apical point; apex acutely angulate. Flagellum without apparent sclerite.

#### Comments

Due to the limited number of specimens of this species, dissections of mouthparts could not be performed.

#### Etymology

Named in reference to the elevated elytral intervals, which give this species a sculpted appearance – *sculptilis* = ‘‘sculpted”; Latin adjective (Brown 1956).

#### Material examined

##### Holotype

♀ “Burley, Idaho; #7, 5-20-32; <u>A</u>.[*rtemisia*] <u>tridentata</u> [non-focal]; David E. Fox” (**USNM**).

##### Paratypes

“Milner, Idaho; #5a, 7-9-31; S.[alsola] *pestifer*; David E. Fox” (**CMNC**: 1 ♀); “Hazelton, Ida; #10 4/29/30; *N.[orta] altissma”* (**USNM**: 1 ♂)

#### Distribution

This species has been found in three localities along the Snake River in Idaho (USA), and is thought to be endemic to the Snake River Plain (Fig. 38).

#### Natural history

Associated with big sagebrush *Artemisia tridentata* Nutt. [non-focal] (Asteraceae [non-focal]), tumbleweed *Salsola tragus* L. [non-focal] (= *Salsola pestifer* A. Nelson [non-focal]) (Amaranthaceae [non-focal]), and tall tumblemustard *Sisymbrium altissimum* L. [non-focal] (= *Norta altissima* (L.) Britt. [non-focal]) (Brassicaceae [non-focal]).

### *Minyomerus tylotos* Jansen & Franz sec. Jansen & Franz, 2018; sp. n

urn:lsid:zoobank.org:act:10CD3562-5969-4BCF-ACFE-BB0E5E2BF9A6 Figures 23-29

**Figure 23.**
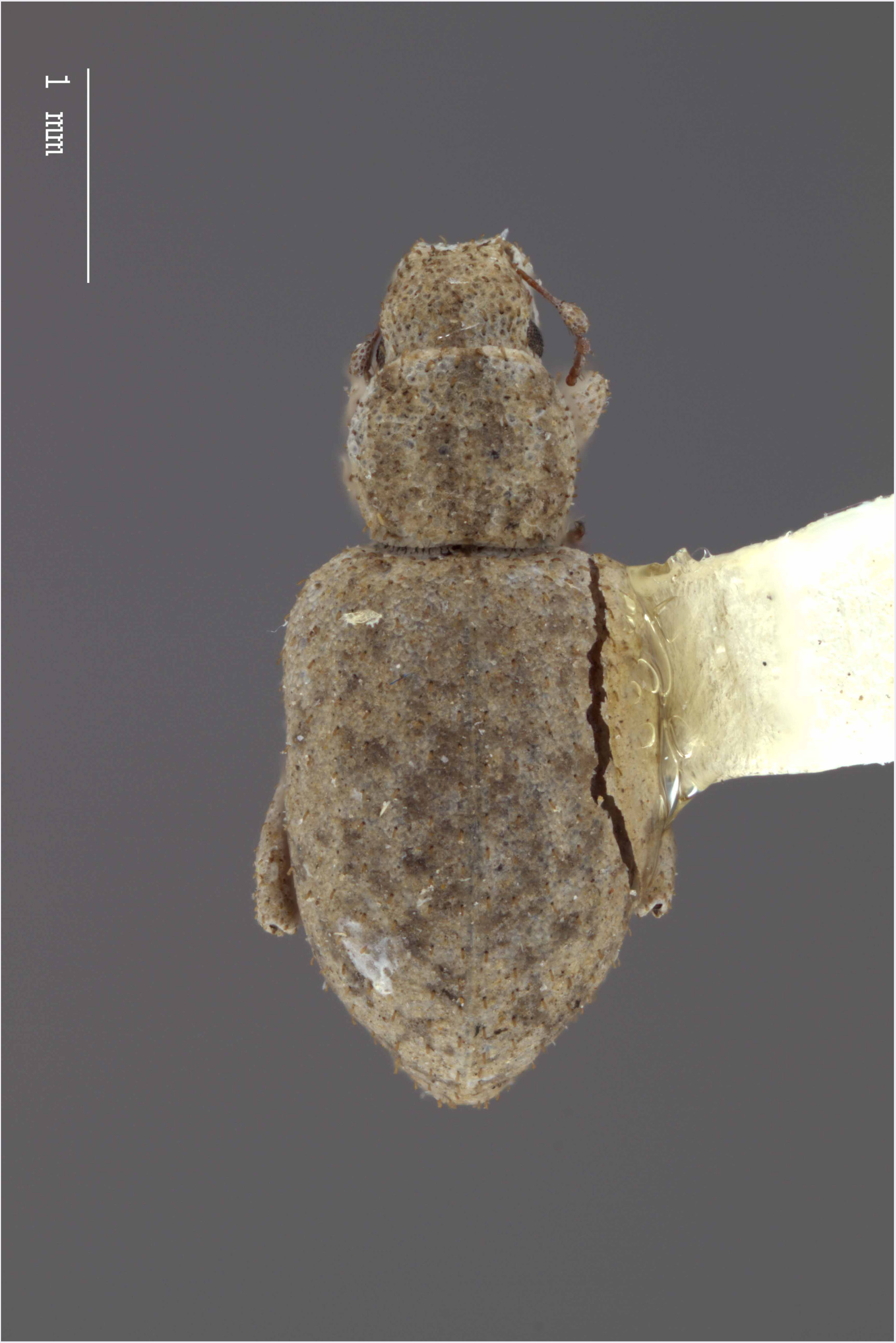
Dorsal habitus of *M. tylotos* [JF2018]. Image of female (♀) holotype.

**Figure 24.**
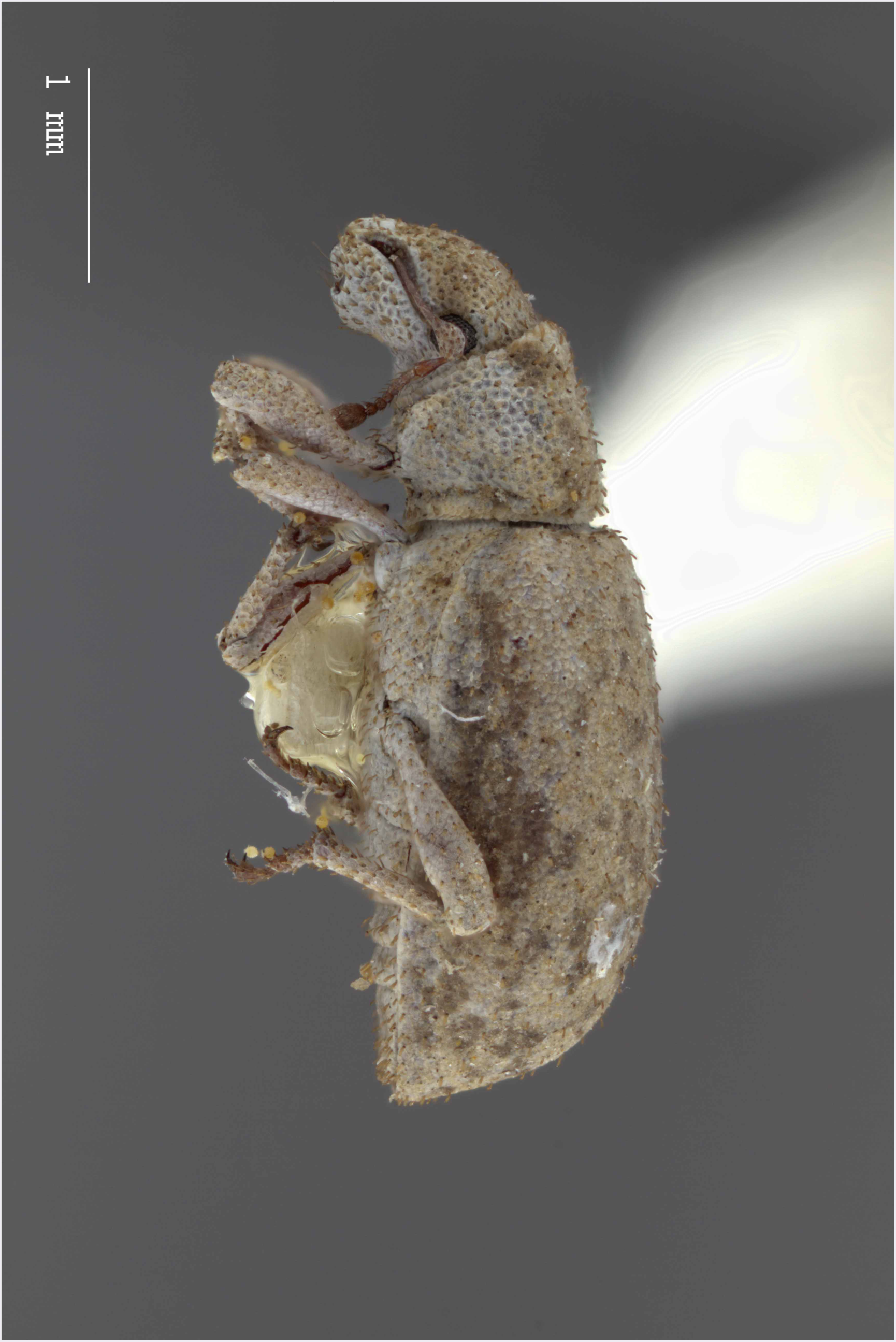
Lateral habitus of *M. tylotos* [JF2018]. Image of female (♀) holotype.

**Figure 25.**
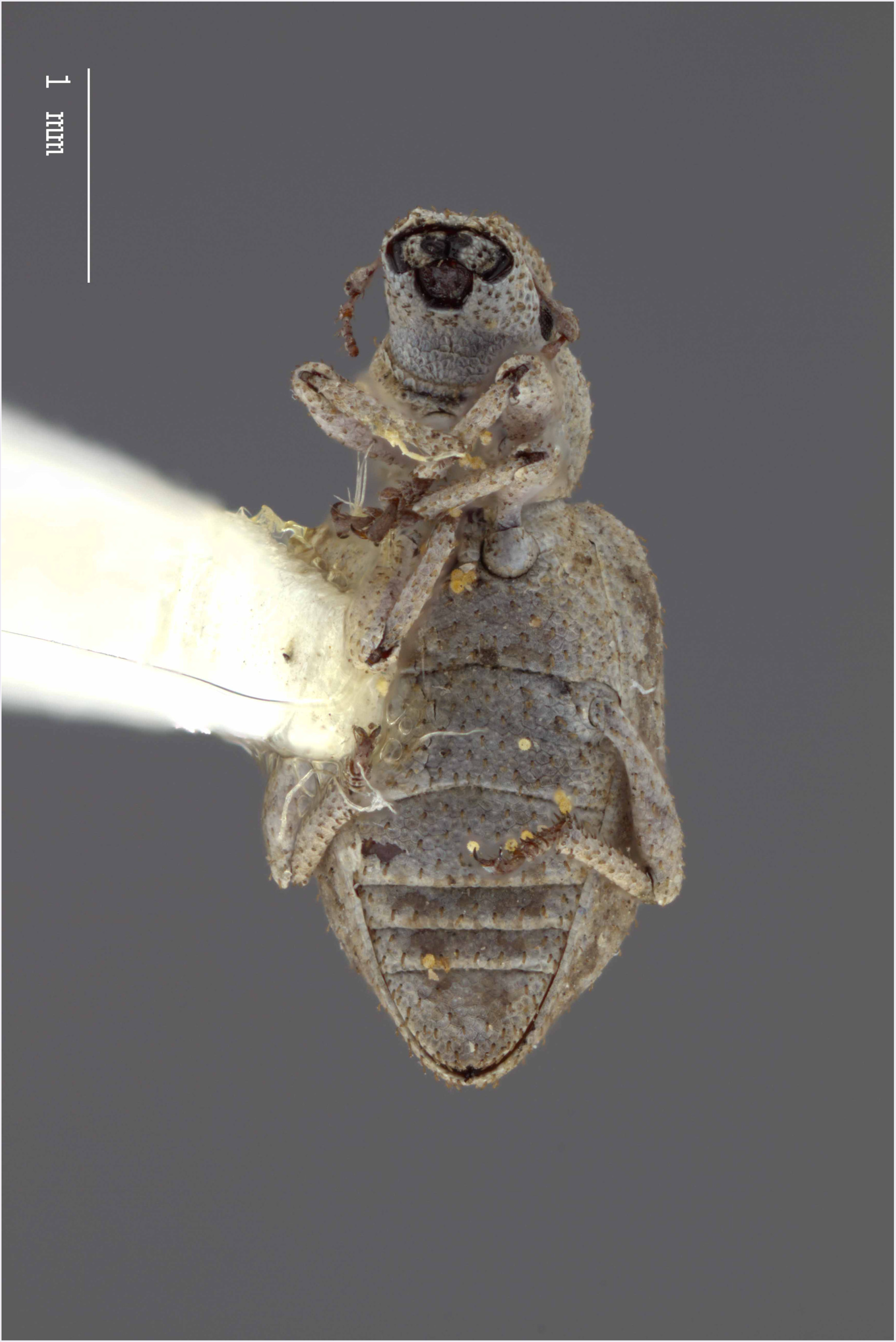
Ventral habitus of *M. tylotos* [JF2018]. Image of female (♀) holotype.

**Figure 26.**
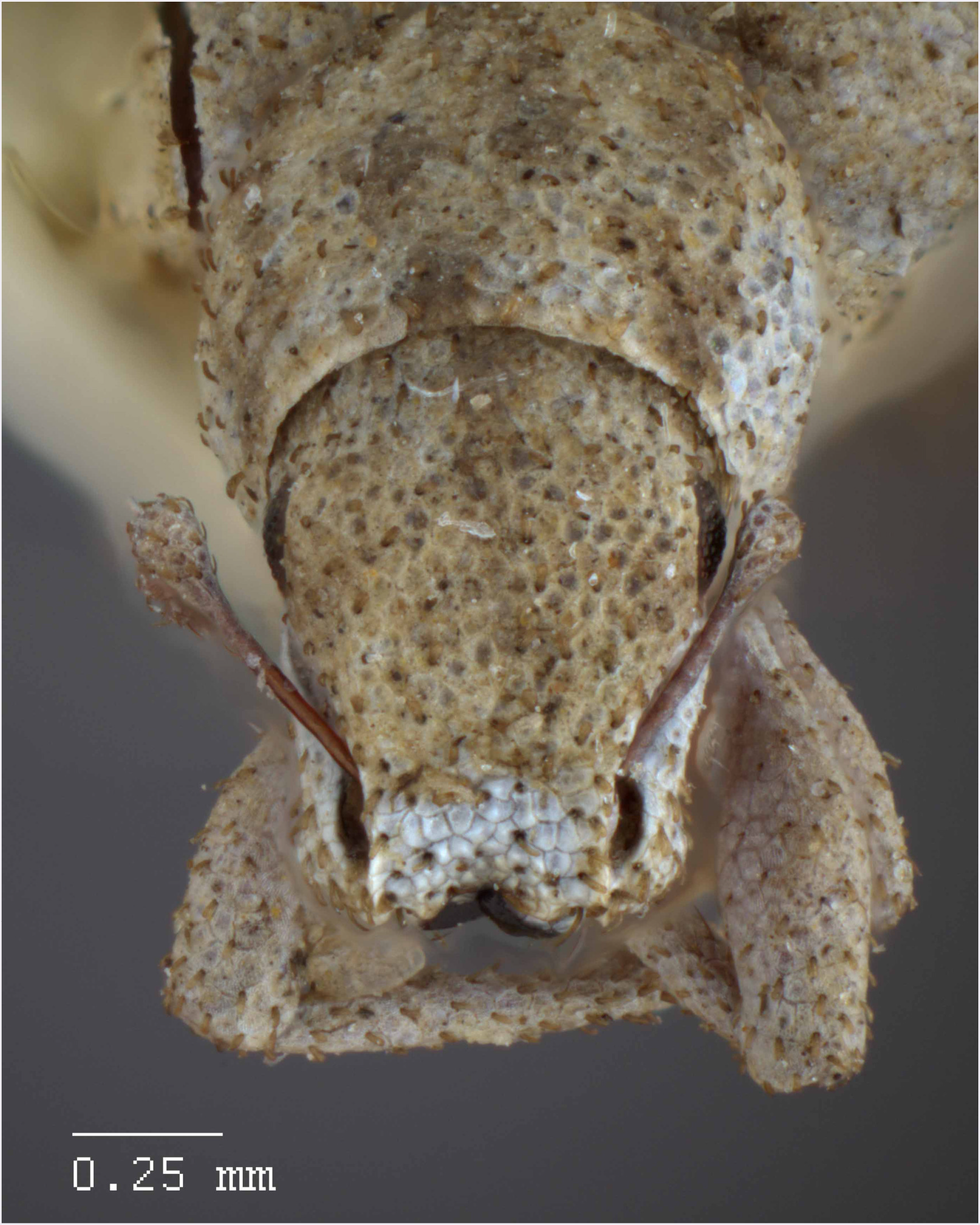
Head and rostrum of *M. tylotos* [JF2018]. Frontal view of female (♀) holotype.

**Figure 27.**
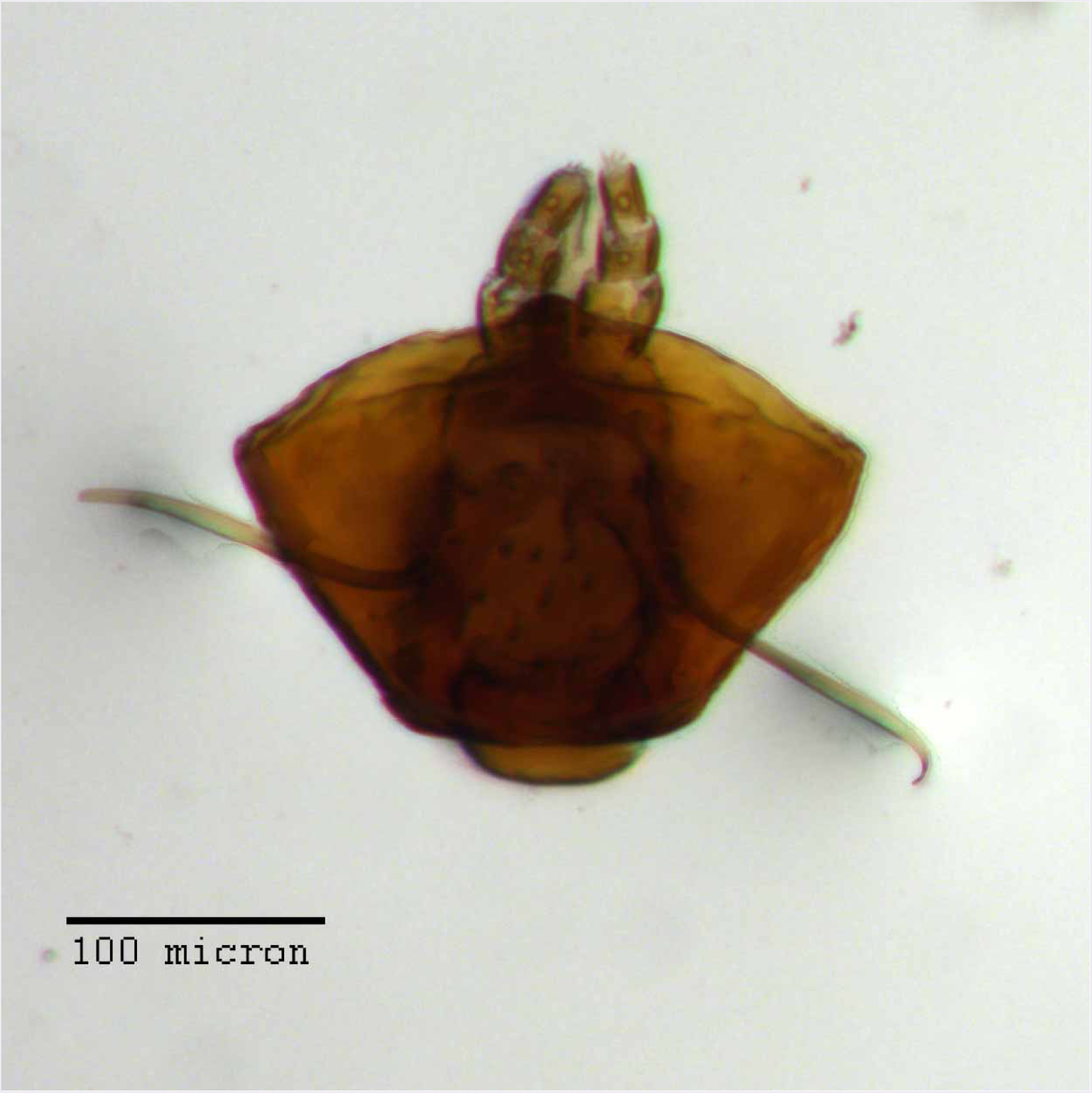
Prementum of *M. tylotos* [JF2018]. Labium of female (♀) paratype.

**Figure 28.**
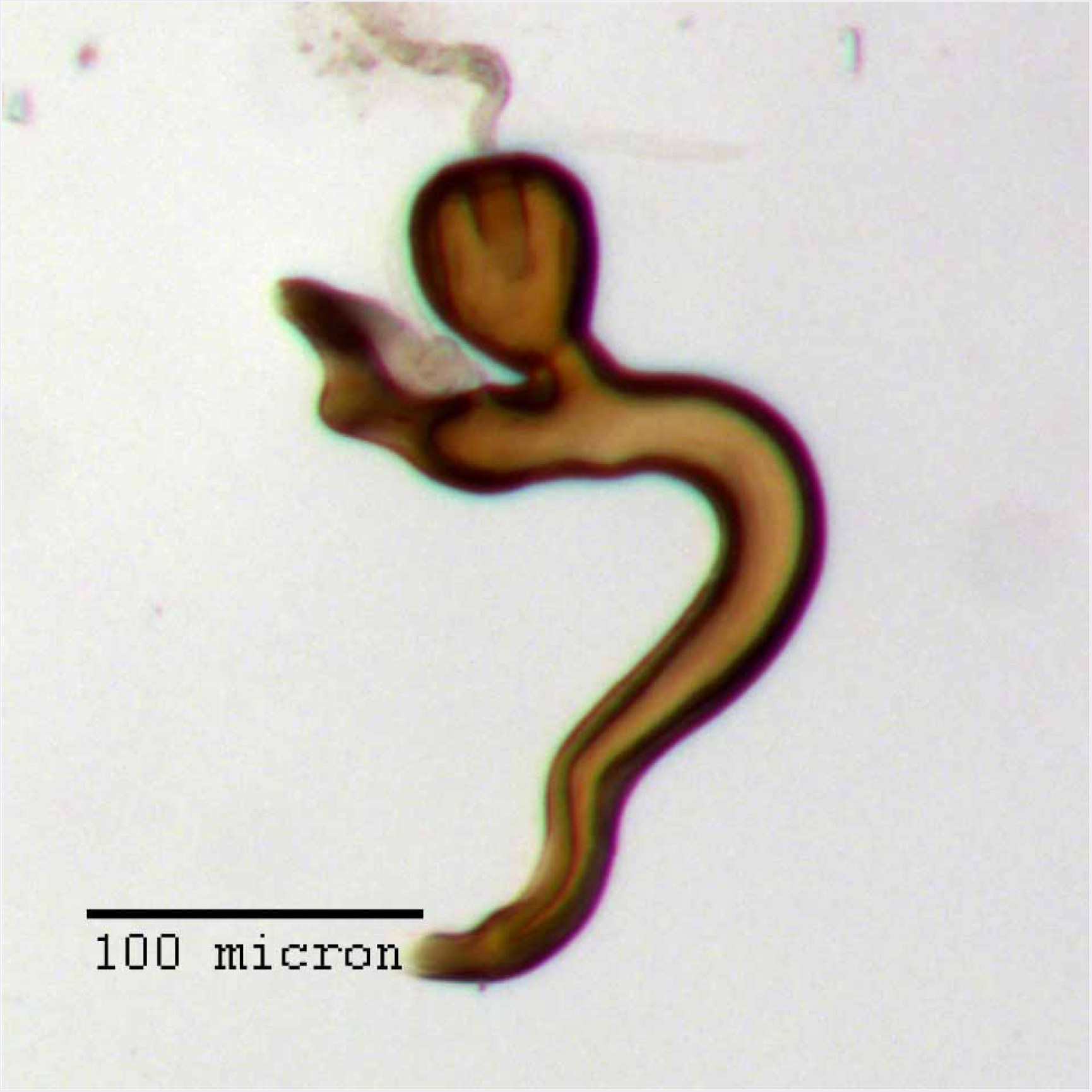
Spermatheca of *M. tylotos* [JF2018]. Genitalia of female (♀) paratype.

**Figure 29.**
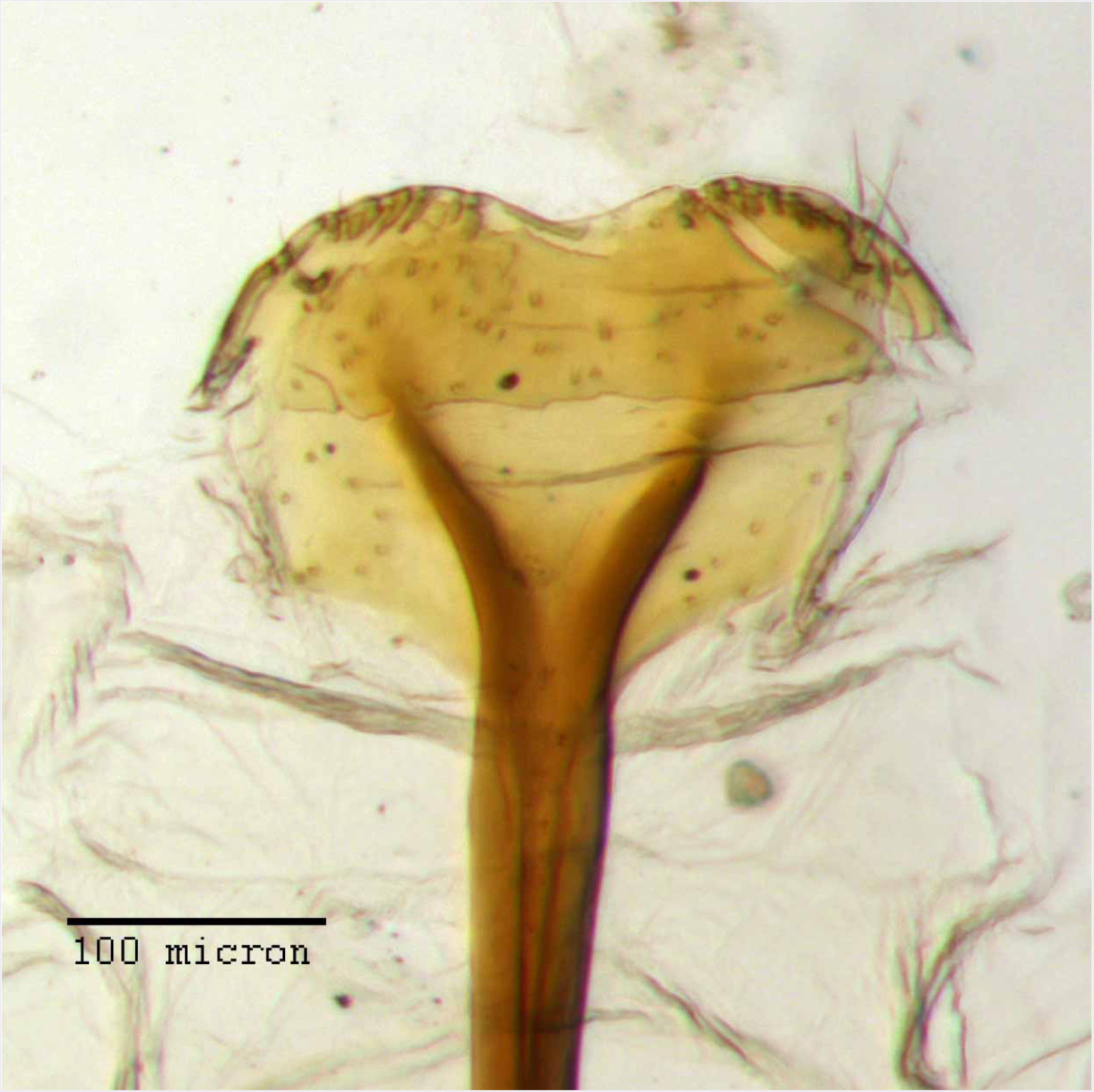
Lamina of spiculum ventrale of *M. tylotos* [JF2018]. Sternum VIII of female (♀) paratype.

**Figure 30.**
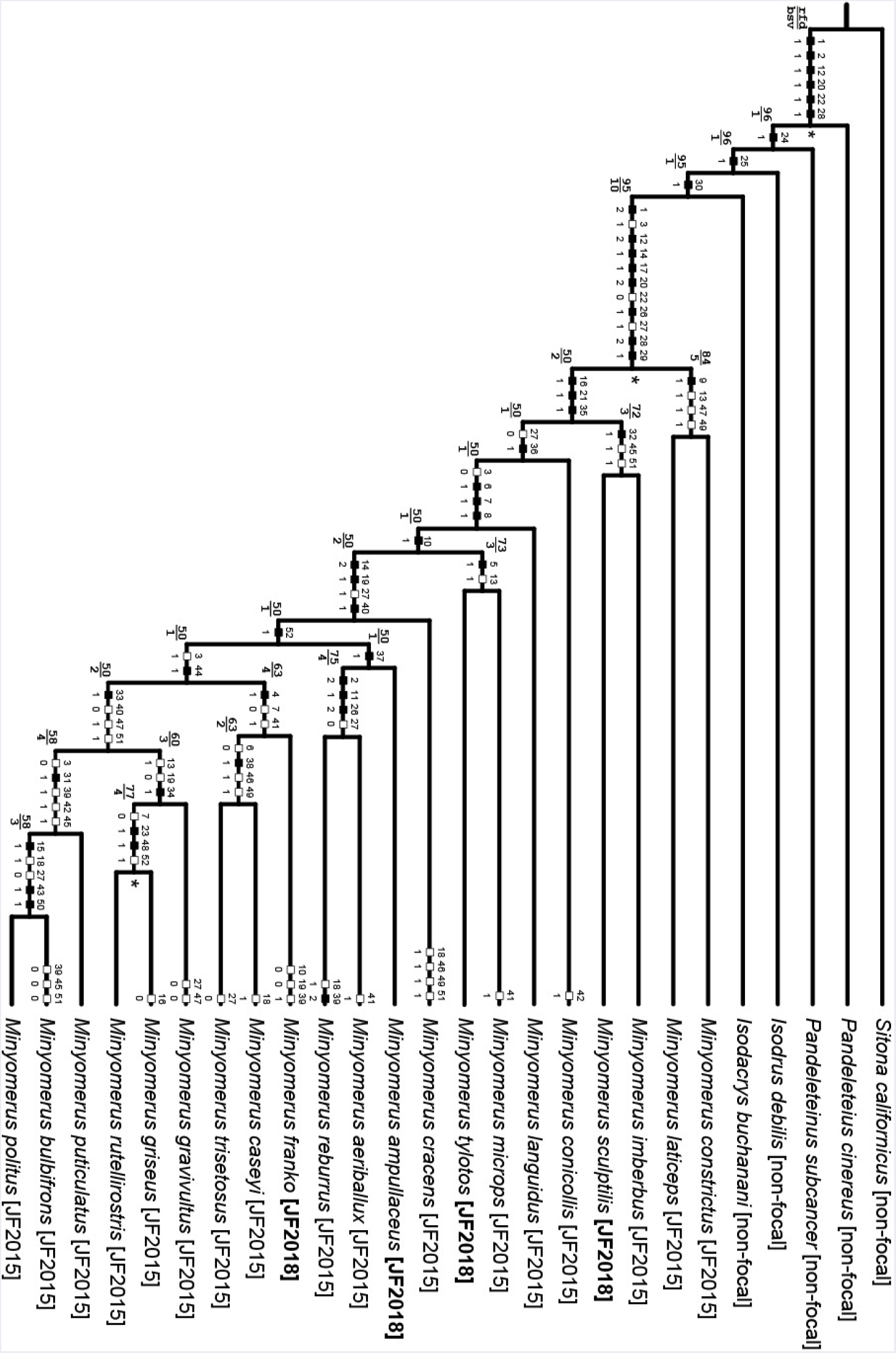
Preferred phylogeny - character transitions and support. Single most parsimonious cladogram representing the preferred phylogeny of species of *Minyomerus* [JF2018], and select outgroup taxa (L = 99, CI = 60, RI = 80). Characters 9, 27, 39, 45 - 47, 49, and 51 are mapped under ACCTRAN optimization; all others are unambiguously optimized. Black squares indicate non-homoplasious character state changes, whereas white squares indicate homoplasious character state changes. The numbers above and below the squares represent character numbers and states, respectively. Bremer support (upper value) and relative fit difference (lower value) values can be found at the left ends of the branches. A “*” symbol at the right end of a branch indicates Bootstrap support greater than 0.95.

**Figure 31.**
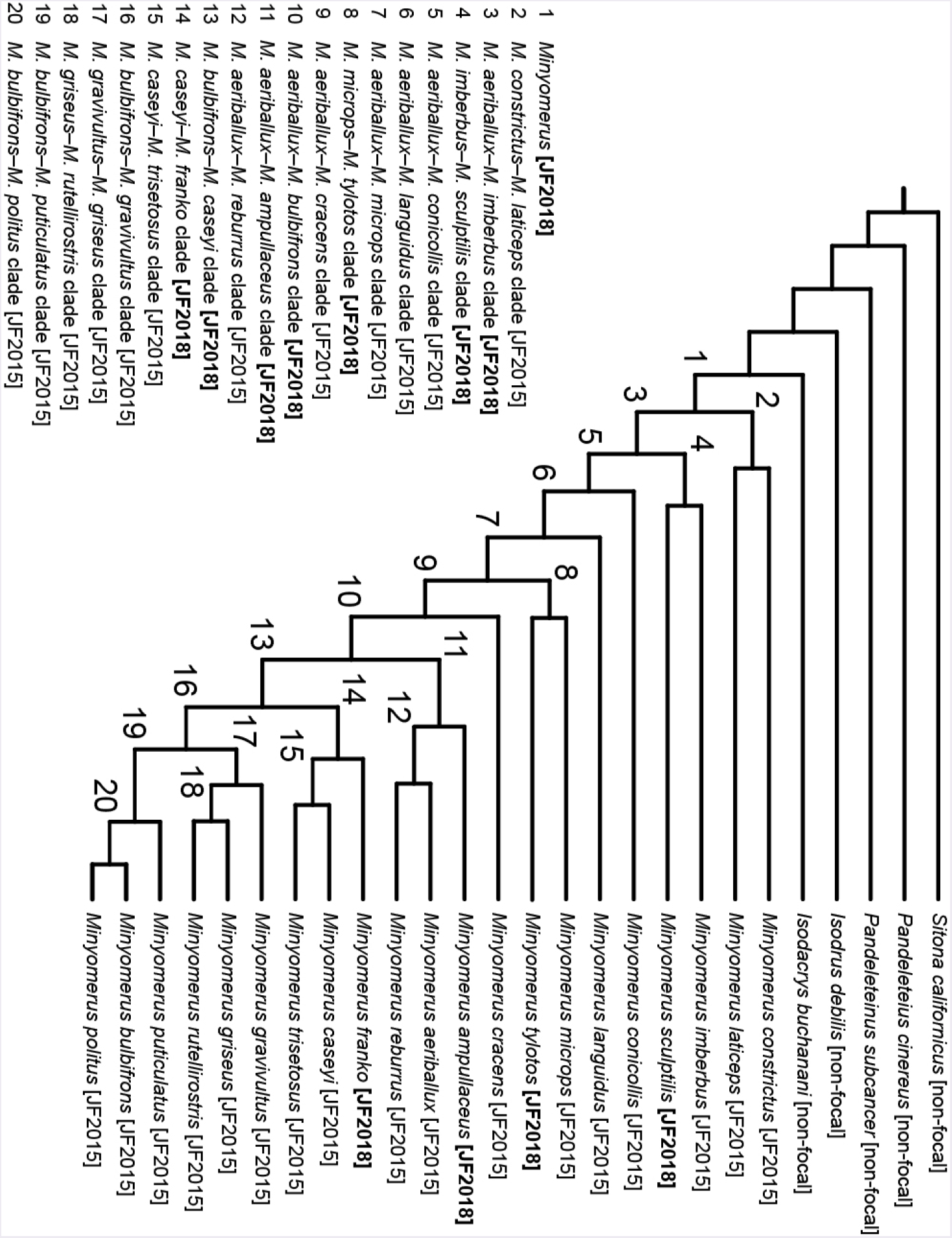
Preferred phylogeny - clade concept labels. Topology and species-level taxonomic concept labels as in Fig. 30. Clade concept labels, numbered 1-20, are consistently generated by using the alphabetically first epithet in each of the bifurcating sister clades. This method safeguards the clade concept labels against changes due simply to reorientation of leaves. Bold-font square brackets indicate new [JF2018] labels. See also RCC-5 Alignments.

**Figure 32.**
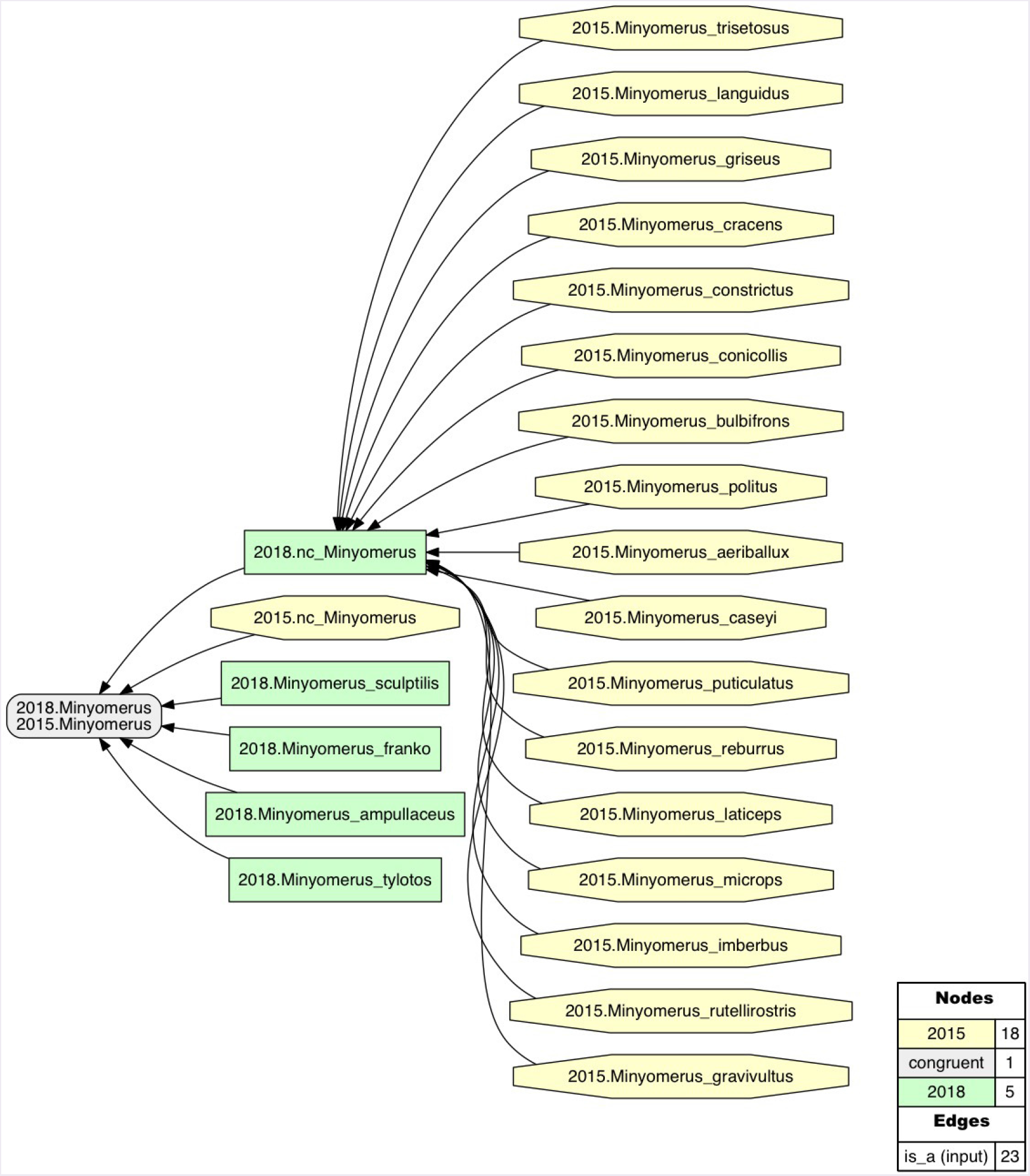
Intensional RCC-5 alignment of the rank-only classifications of *Minyomerus* [JF2018]/[JF2015]See also Jansen & Franz (2015) and Supplemental Information SI2. Taxonomic concept labels such as *Minymerus microps* [JF2015] are abbreviated as “2015. Minyomerus_microps”. Relaxation of the coverage constraint is indicated with the prefix “nc_” (no coverage). Congruent concept regions (T_2_ and T_1_) are shown as grey rectangles, concepts regions unique to the later taxonomy (T_2_) are shown as green rectangles, and concept regions unique to the earlier taxonomy (T_1_) are shown as yellow octagons. Articulations of inverse proper inclusion (<) and overlap (><), where present, are also shown.

**Figure 33.**
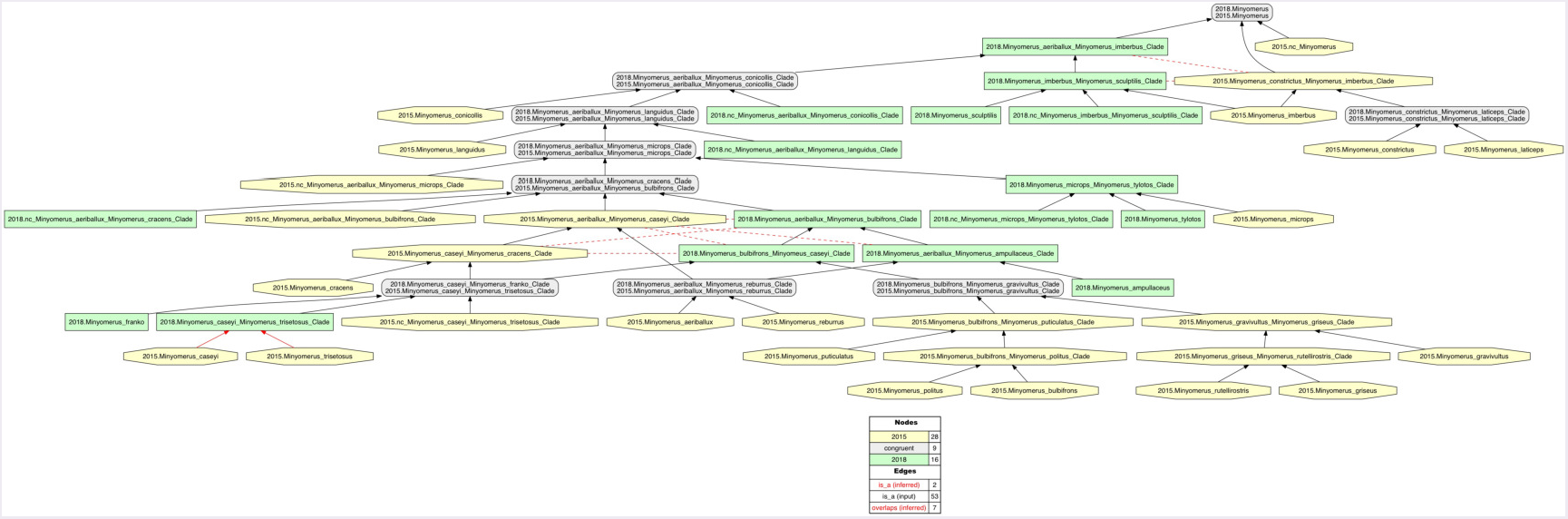
Intensional RCC-5 alignment of the phylogenies of *Minyomerus* [JF2018]/[JF2015] - whole-concept resolution with overlap. See also Supplemental Information SI3. Seven overlapping articulations are inferred. For further discussion, see the RCC-5 Alignments section.

**Figure 34.**
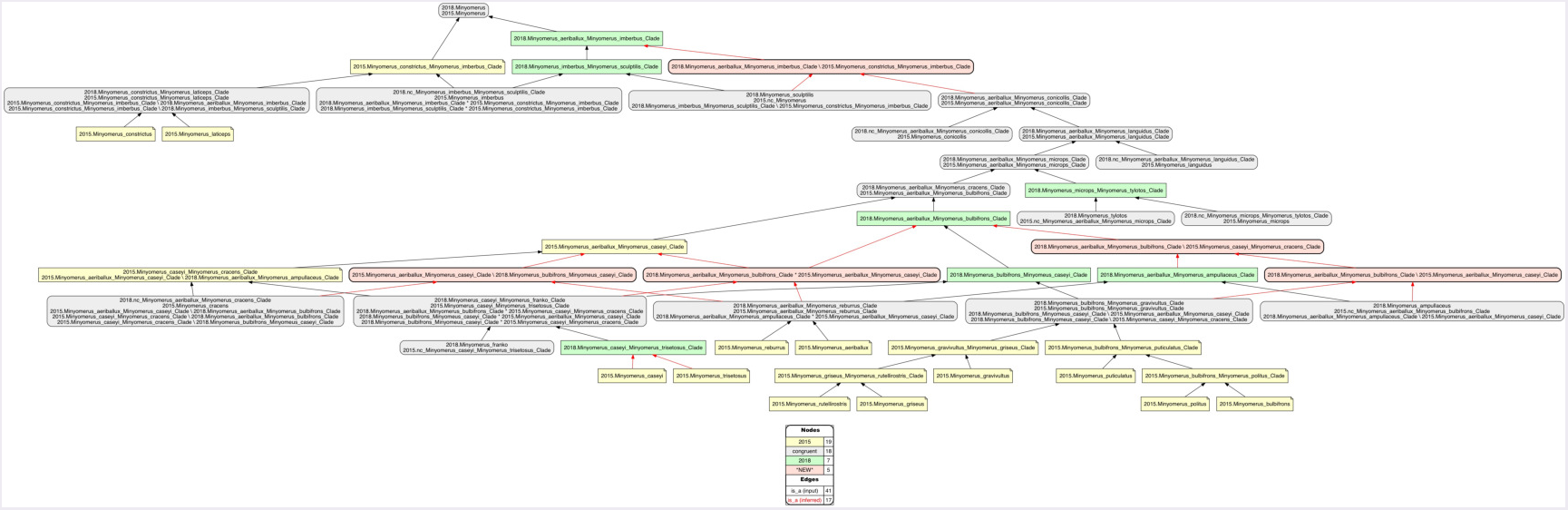
Intensional RCC-5 alignment of the phylogenies of *Minyomerus* [JF2018]/[JF2015] - split-concept resolution. See also Supplemental Information SI4. The seven overlapping articulations of the alignment displayed Fig. 33 are resolved into their constituent split regions. That is, if regions A and B overlap, the three resulting split regions are labeled A\b (“A, *not* b”), A*B (“A *and* B”), and B\a (“B, *not* a”). Five split-concept regions can *only* be named using this convention, and are salmon-colored in the alignment visualization.

**Figure 35.**
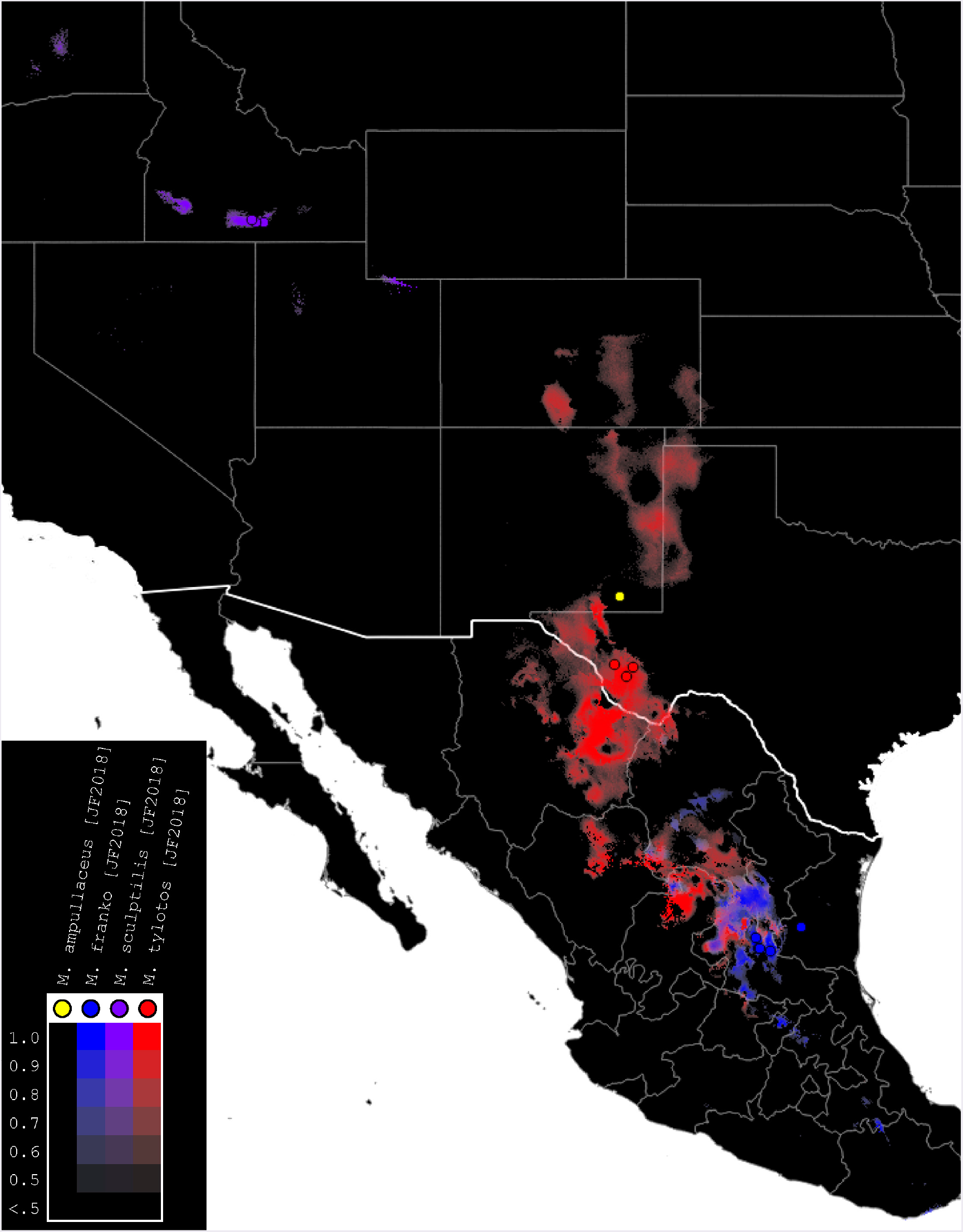
Summary map of distributions of new species of *Minyomerus* [JF2018]. Combined occurrence record and Maxent habitat modeling map for four newly-described species of *Minyomerus* [JF2018], as indicated in the legend.

**Figure 36.**
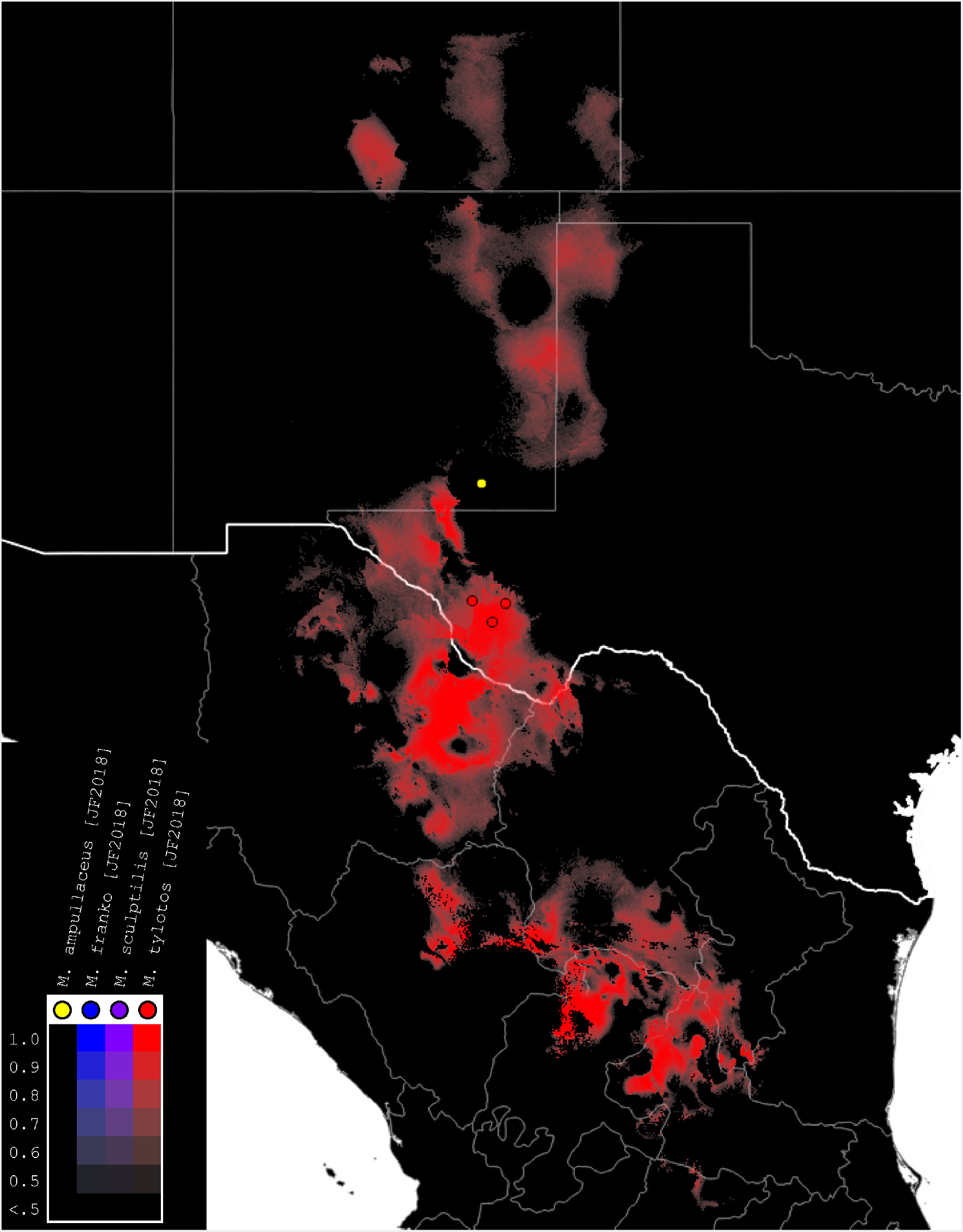
Distributions of *M. ampullaceus* [JF2018] and *M. tylotos* [JF2018]. Combined occurrence record and Maxent habitat modeling map for *M. ampullaceus* [JF2018] and *M. tylotos* [JF2018], as indicated in the legend.

**Figure 37.**
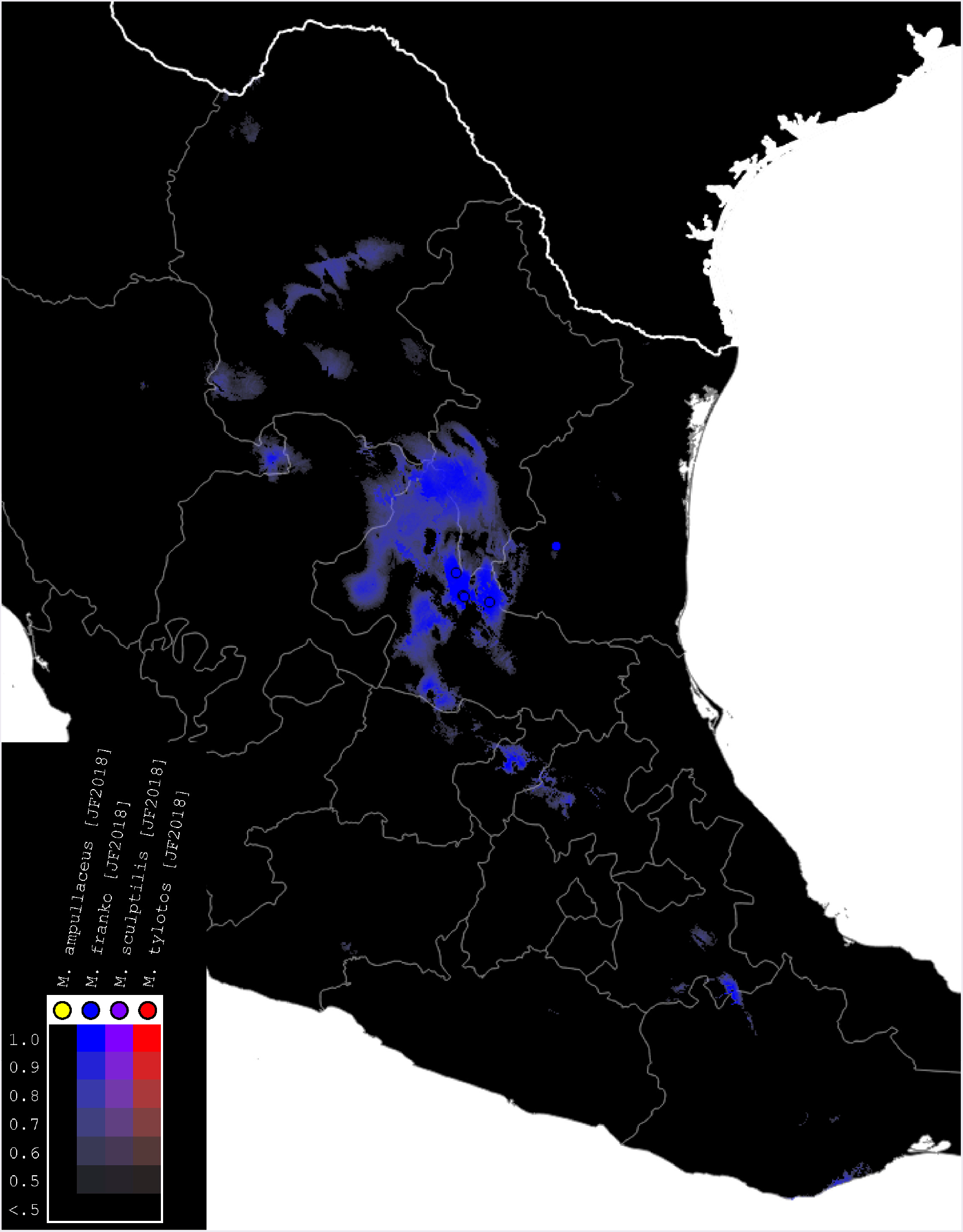
Distributions of *M. franko* [JF2018]. Combined occurrence record and Maxent habitat modeling map for *M. franko* [JF2018], as indicated in the legend.

**Figure 38.**
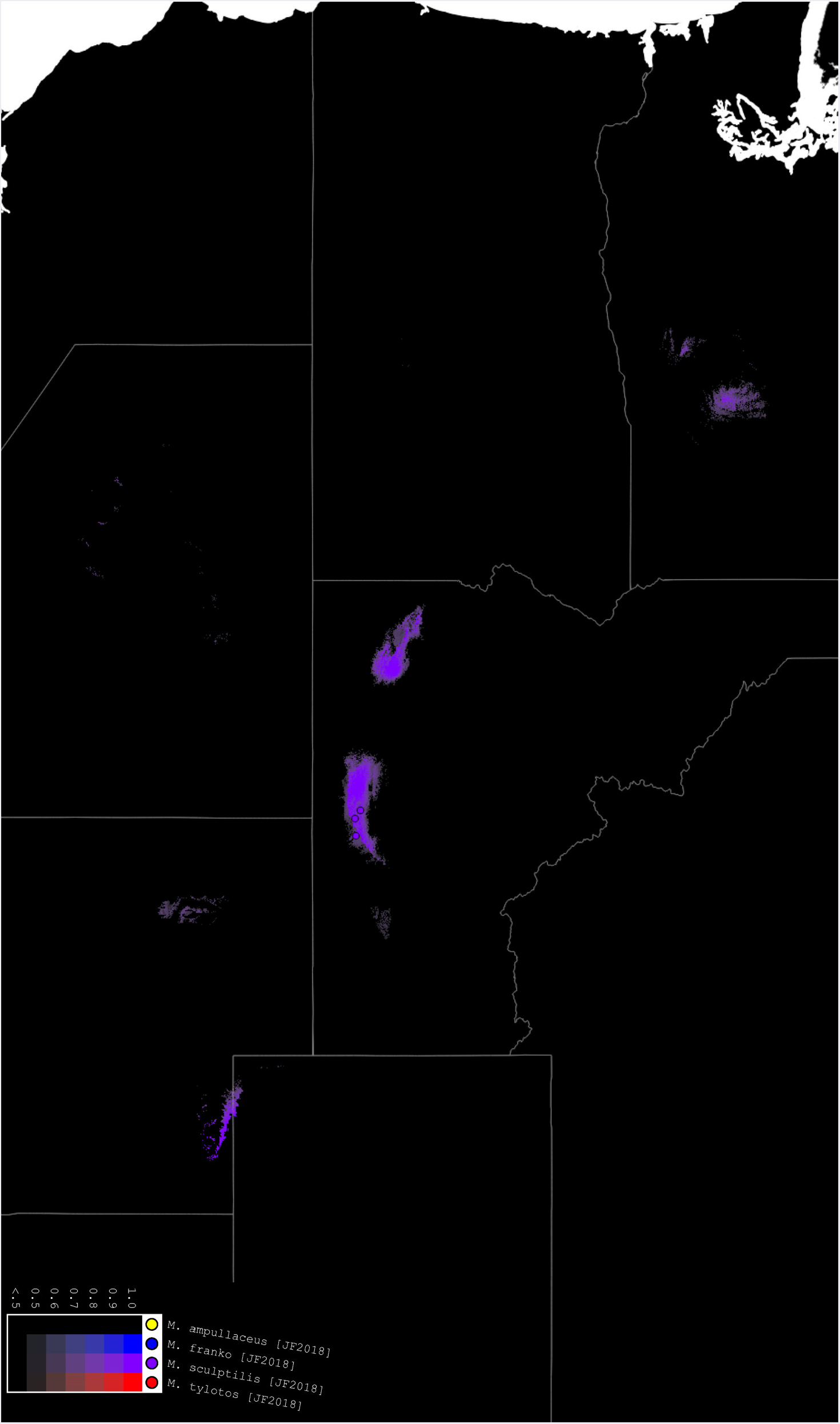
Distributions of *M. sculptilis* [JF2018]. Combined occurrence record and Maxent habitat modeling map for *M. sculptilis* [JF2018], as indicated in the legend.

#### Diagnosis

*Minyomerus tylotos* [JF2018] is most readily distinguished from other congenerics by a combination of characters, as follows. The nasal plate lacks distinct impressions, having instead a poorly defined anteromesal convexity completely and evenly covered with white scales. The frons is protuberant and moderately punctate. The entire body, including the legs, head, and venter, are clothed with brown, linear to minutely apically expanded setae, which are of similar length throughout and appear distinctly undifferentiated and uniform across body regions. The body is somewhat bulky, with the pronotum protuberant laterally and globular in dorsal view. The setae lining the anterodorsal margin of the pronotum uniquely apically explanate, with a longitudinal, medial, ridge-like portion that tapers to either side apicolaterally (visible at high magnification). The lateral margins of the elytra are protuberant anteriorly and sub-parallel along the between anterior 1/5 and posterior 1/3 of their length. The spermatheca has the corpus narrow throughout, equal in thickness to the collum. The ramus is basally stalked and apically bulbous. The collum exhibits a double-bend, and is recurved.

#### Description of female

##### Habitus

Length 3.46-3.62 mm, width 1.42-1.54 mm, length/width ratio 2.35-2.44, widest at anterior 1/6 of elytra. Integument orange-brown to black. Scales with variously interspersed colors ranging from slightly off-white or beige to manila/tan to dark coffee brown, in some specimens appearing semi-translucent (in others opaque). Setae linear to apically explanate, appearing minutely spatulate, sub-recumbent to sub-erect, tan to brown in color.

##### Mandibles

Covered with white scales, with 2-3 longer setae, and 1-3 shorter setae between these.

##### Maxillae

Cardo bifurcate at base with an inner angle of ca. 90°, arms roughly equal in length and width, arms of bifurcation equal in length to apically outcurved arm. Stipes sub-rectangular, 1.5× wider than long, roughly equal in width to inner arm of bifurcation of cardo, glabrous. Galeo-lacinial complex nearly extending to apex of maxillary palpomere I; complex mesally membranous, laterally sclerotized, with sharp demarcation of sclerotized region separating palpiger from galeo-lacinial complex; setose in membranous area just adjacent to sclerotized region, setae covering 1/2 of dorsal surface area; dorsally with 5 apicomesal lacinial teeth; ventrally with 3 reduced lacinial teeth. Palpiger with a single lateral seta, otherwise glabrous, anterior 1/2 membranous, posteriorly sclerotized.

##### Maxillary palps

I apically oblique, apical end forming a 45° angle with base, with 2 apical setae; II sub-cylindrical, with 1 apical seta.

##### Labium

Prementum roughly pentagonal; apical margins arcuate, medially angulate; lateral margins feebly incurved; basal margin arcuate. Labial palps 3-segmented, I with apical 1/2 projecting beyond margin of prementum, reaching apex of ligula; III slightly longer than II.

##### Rostrum

Length 0.49-0.50 mm, anterior portion 2.25-2.5 × broader than long, rostrum/pronotum length ratio 0.58-0.62, rostrum length/width ratio 1.26-1.32. Separation of rostrum from head generally obscure. Dorsal outline of rostrum nearly square, anterior half of dorsal surface feebly impressed, posterior half coarsely but shallowly punctate to rugose. Rostrum in lateral view nearly square; apical margin strongly bisinuate and emarginate, appearing medially notched, with 2 large vibrissae. Nasal plate lacking distinct impressions, having instead a poorly defined anteromesal convexity, integument completely and evenly covered with white scales. Margins of mandibular incision directed ca. 25-30° outward dorsally in frontal view. Ventrolateral sulci weakly defined as a broad concavity dorsad of insertion point of mandibles, running parallel to scrobe, becoming flatter posteriorly and disappearing ventrally. Dorsal surface of rostrum with median fovea short and linear, or punctate. Rostrum ventrally with sub-parallel sulci beginning at corners of oral cavity and continuing halfway to back of head.

##### Antennae

Minute tooth formed by overhanging dorsal margin of scrobe anterior to margin of eye by 1/3 of length of eye. Scape extending to posterior margin of eye. Terminal funicular antennomere lacking appressed scales, having instead a covering of apically-directed pubescence with interspersed sub-erect setae. Club nearly 3 × as long as wide.

##### Head

Eyes globular and somewhat elongate, strongly impressed, slanted ca. 45° antero-ventrally; eyes separated in dorsal view by 4× their anterior-posterior length, set off from anterior prothoracic margin by 1/4 of their anterior-posterior length. Head between eyes punctate and protuberant.

##### Pronotum

Length/width ratio 0.88-0.89; widest near anterior 2/5; somewhat globular. Anterior margin arcuate, but feebly incurved mesally, lateral margins evenly curved and widening into a bulge just anteriad of midpoint of pronotum, posterior margin straight, with a slight mesal incurvature. Pronotum in lateral view with transverse ventrolateral sulci strongly excavated and distinctly sculptured; with short, recumbent to sub-erect setae that barely attain or reach just beyond anterior margin; these setae becoming shorter and more erect laterally, reaching a maximum length nearly equal to length of eye; dorsally, these setae become uniquely apically explanate, with a longitudinal, medial, ridge-like portion that tapers to either side apicolaterally. Anterolateral margin with a single ocular vibrissa present, emerging near ventral margin of eye; vibrissa achieving a maximum length of 2/5 of anterior-posterior length of eye.

##### Scutellum

Not exposed.

##### Pleurites

Metepisternum nearly hidden by elytron except for triangular extension.

##### Thoracic sterna

Mesocoxal cavities separated by 1/3 × width of mesocoxal cavity. Metasternum with transverse sulcus not apparent; metacoxal cavities widely separated by ca. 3× their width.

##### Legs

Profemur/pronotum length ratio 0.90-0.96; profemur with distal 1/5 produced ventrally as a sub-rectangular projection covering tibial joint; condyle of tibial articulation occupying 4/5 of distal surface and 1/5 length of femur. Protibia/profemur length ratio 0.86-0.91; protibial apex with ventral setal comb recessed in a subtly incurved groove; mucro present as an acute, medially-projected tooth, which is approximately equal in length to surrounding setae. Protarsus with tarsomere III 2× as long as II; wider than long. Metatibial apex with weakly projecting, poorly defined, narrow convexity laterally flanged by 5 short, spiniform setae.

##### Elytra

Length/width ratio 3.03-3.21; widest at anterior 1/6; anterior margins jointly 1.5-2× wider than posterior margin of pronotum; lateral margins nearly straight and sub-parallel after anterior 1/5, converging in posterior 1/3. Posterior declivity angled at 70-75° to main body axis. Elytra with 10 complete striae; striae broadly sculpted; punctures broad and faint beneath appressed scales, separated by 4-5 × their diameter; intervals elevated.

##### Abdominal sterna

Ventrite III anteromesally incurved around a fovea located mesally on anterior margin, posterior margin elevated and set off from IV along lateral 3/8s of its length. Sternum VII mesally 2/3 × as long as wide; setae slightly lengthening, and becoming medially directed in posterior 1/3; anterior margin weakly curved; posterior margin distinctly incurved mesally, appearing broadly notched; surface of sternite concave, appearing broadly foveate, immediately anteriad of marginal incurvature.

##### Tergum

Tergum VII mesally incurved. Pygidium sub-cylindrical; medial 1/3 of anterior 2/3 of pygidium less sclerotized, with a patch of very short, fine setae.

##### Sternum VIII

Anterior laminar edges each incurved forming a 130° angle with lateral margin; slightly less sclerotized medially between arms; posterior margin medially incurved.

##### Ovipositor

Coxites as long as broad; styli with 3 setae near the base.

##### Spermatheca

?-shaped; collum short, apically with a large, angulate, hood-shaped projection angled at 45° to corpus, sub-equal in length to ramus and contiously aligned with curvature of bulb of ramus; collum sub-contiguous with, and angled at ca. 60° to ramus; ramus basally elongate and constricted, forming a stalk, 1/3 × length of collum, bulbous apically, 3 × thicker than stalk; corpus not swollen, of equal thickness to collum and cornu; cornu elongate, apically, gradually narrowed, strongly recurved in basal 1/3, straight along mesal 1/3, and curved near apical 1/3 such that apex is parallel to collum and corpus.

#### Description of male

Not available or known.

#### Etymology

Named in reference to the short, apically explanate setae interspersed throughout the dorsum, which give this species a distinctly “knobbed” appearance; *tylotos* – knobby; Greek adjective (Brown 1956).

#### Material examined

##### Holotype

♀ “H. O. Canyon,; Davis Mts., Texas; Jeff Davis County; VII-20-1968, 6200’; J. E. Hafernik” (**TAMU**).

##### Paratypes

“24 mi. wsw. Ft. Davis; Jeff Davis Co., Texas; August 17, 1969; Board & Hafernik” (**TAMU**: 1 ♀); “USA Texas Jeff Davis Co.; 4.1 mi. S. Fort Davis; sweeping grasses-weeds; 4750’. 19.VII.82; R.S. Anderson” (**CMNC**: 1 ♀)

#### Distribution

This species has been found in three localities near the Davis Mountains in Jeff Davis County and in nearby Presidio County, Texas (USA). Habitat models (Figs. 36) predict that this represents the northeastern extent of its range, indicating a strong likelihood that it is present in other parts of the northern Chihuahuan desert, especially in the state of Chihuahua (Mexico).

#### Natural history

No host plant associations have been documented. It is unknown whether this species is parthenogenetic.

## CHECKLIST OF SPECIES

RCC-5 articulations are provided in bold font. See Jansen & Franz (2015) for alignments of *Minyomerus* concepts published from 1831 to 2015.

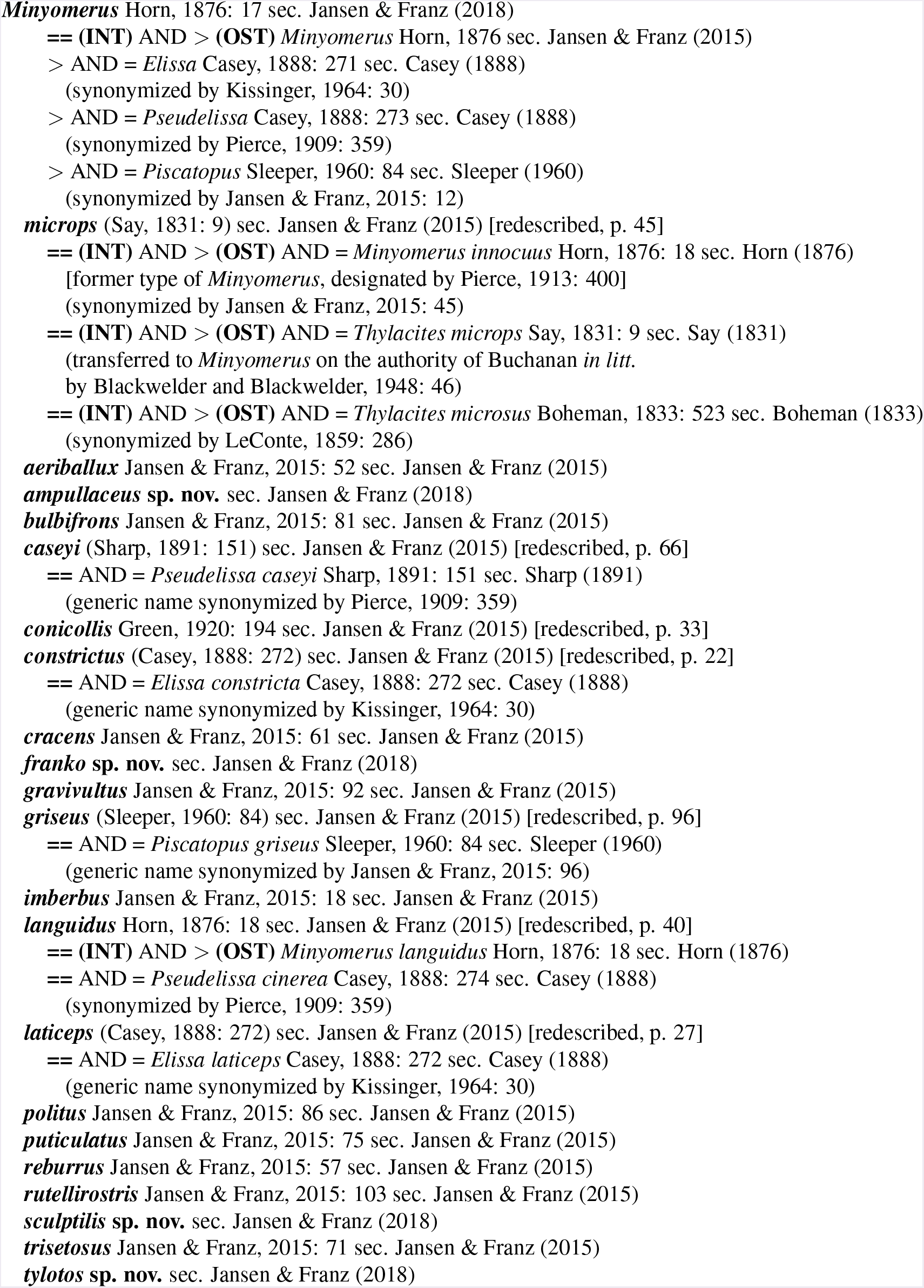

## SPECIES IDENTIFICATION KEY

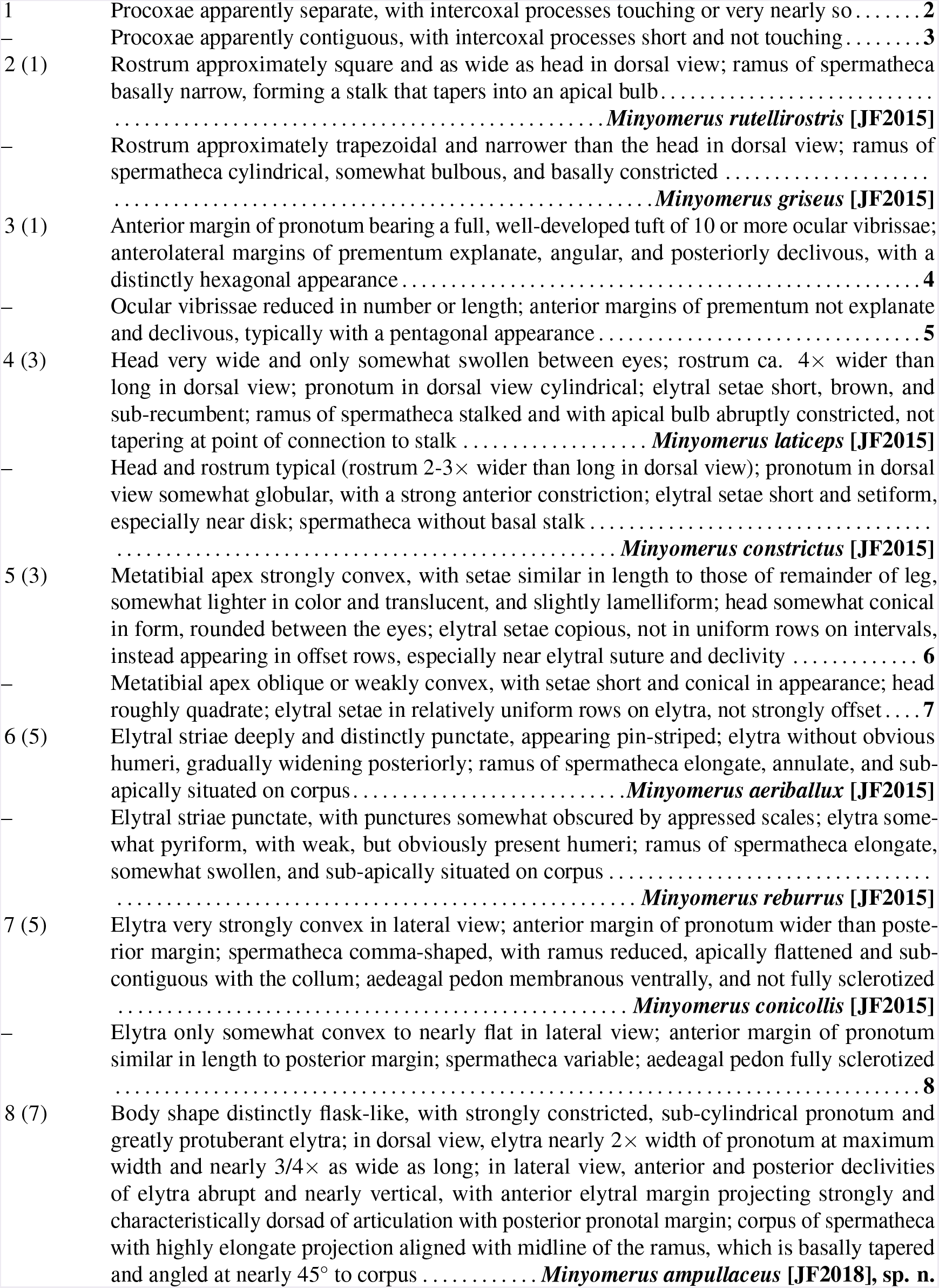
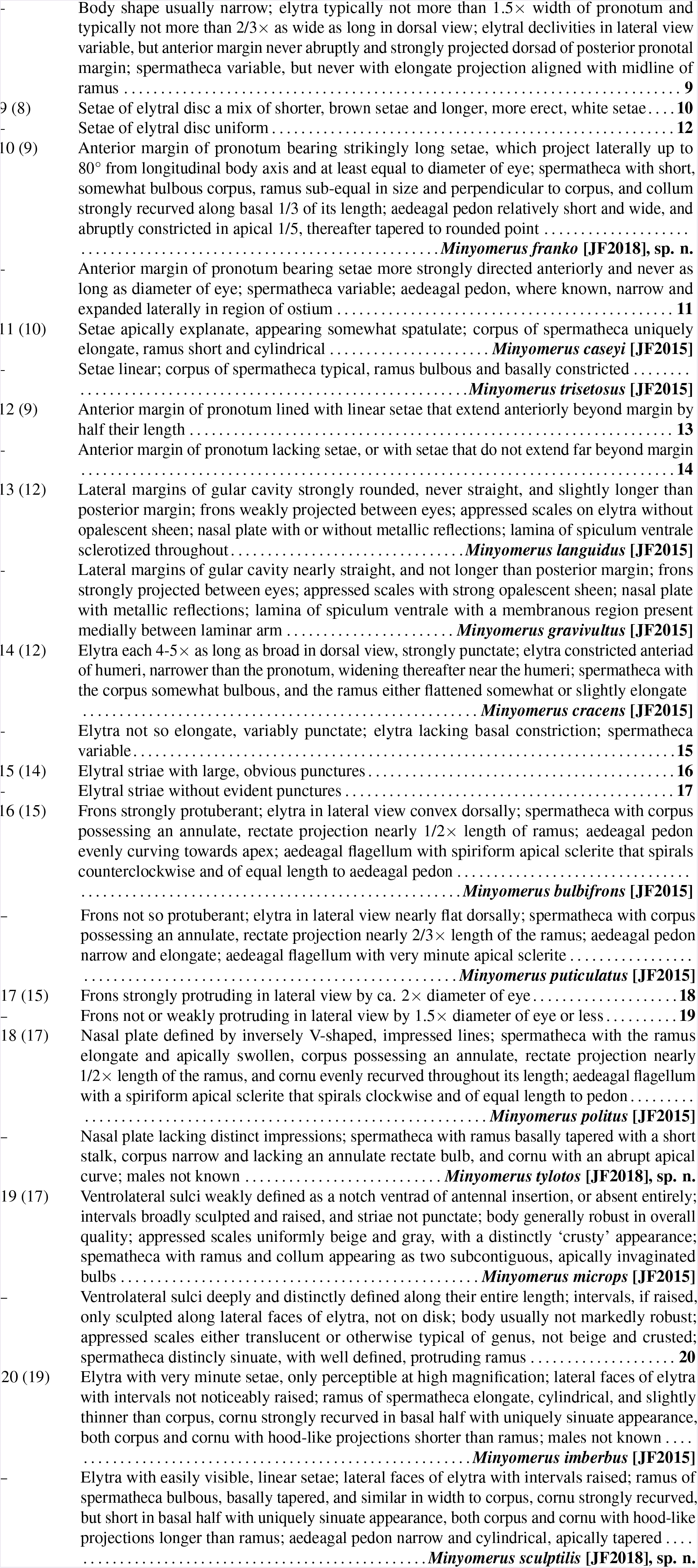

## PHYLOGENETIC RESULTS

A matrix of 52 characters was assembled for the 26 terminal taxa (Tab. 1). These characters are comprised of all 46 characters included in the revision of *Minyomerus* [JF2015], plus an additional 6 characters intended to identify putative sister taxa to the newly described species. Parsimony analysis returned a single, most-parsimonious cladogram (henceforth MPT) with a length (L) of 99 steps, a consistency index (CI) of 60 and a retention index (RI) of 80 (Farris 1989); see Figs. 30-31. TNT (Tree Analysis Using New Technology) was used to confirm that the shortest tree had been found (Goloboff et al. 2008). The most-parsimonious cladogram is shown in Fig. 30, with relative and absolute Bremer support values (see also **Materials and Methods: Phylogenetic analysis**) mapped along the left side of each branch; nodes with bootstrap support above 0.95 are marked with a “*” symbol to the right of each node. In a complementary graph, we show the herein used clade concept labels (Fig. 31).

The characters, states, and preferred optimizations are described in this section. Characters relating to placement of the herein described taxa are discussed in detail in the **Discussion** section, along with changes in species group composition and tree topology from Jansen & Franz (2015). For all characters not resolved as unreversed synapomorphies, both the individual consistency (ci) and retention (ri) indices are provided.

**Table 1.**
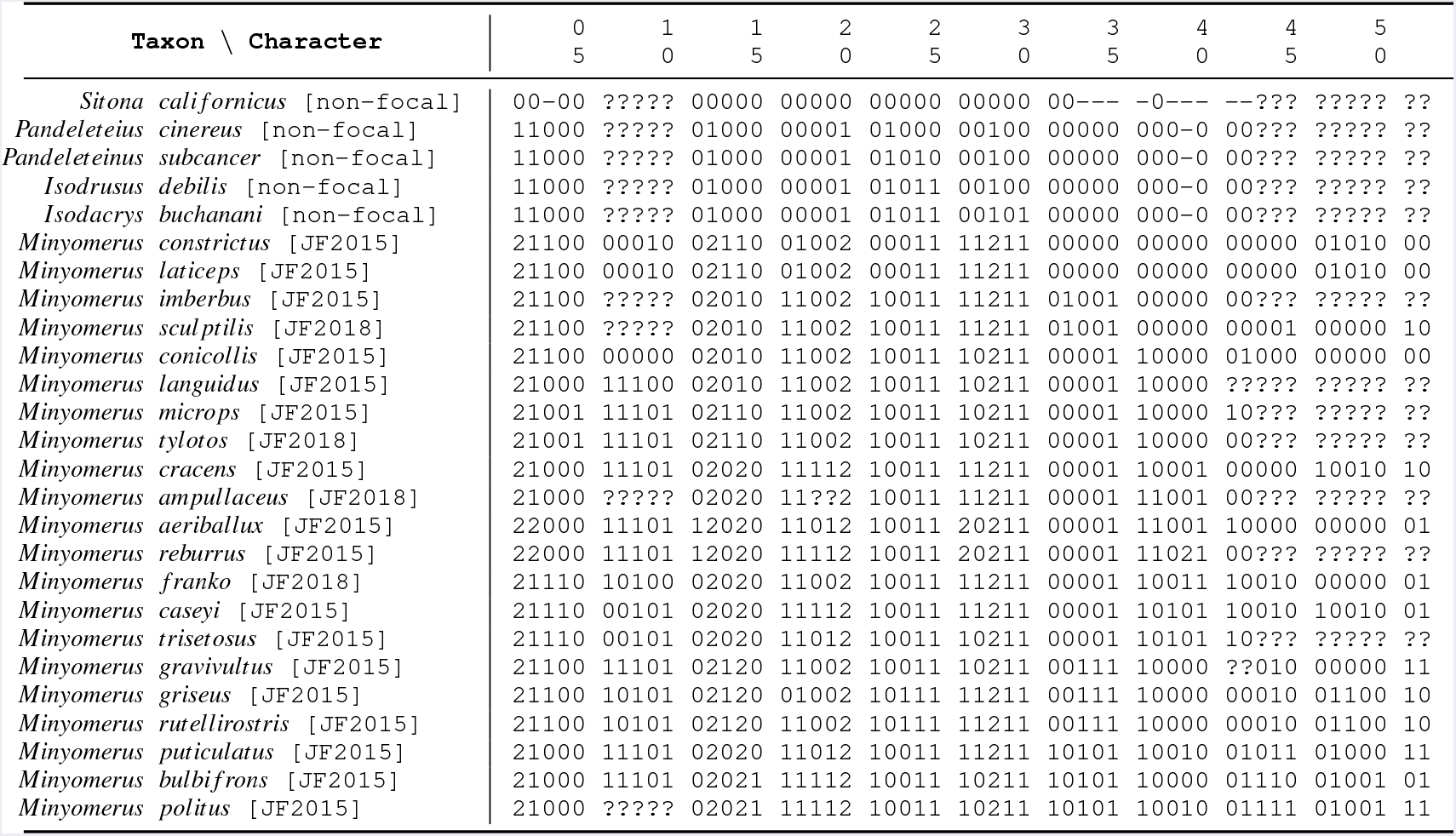
Taxon/character matrix used for for cladistic analysis. Includes all species of *Minyomerus* [JF2015], newly designated species, and select outgroup taxa. All multi-state characters coded as additive, except for character 33. The symbol “–” denotes inapplicable character states, whereas “?” denotes missing information (see also text).

1. Habitus, form of appressed scales: (0) elongate pyriform, not overlapping; (1) sub-circular to polygonal, variously overlapping non-linearly; (2) sub-circular and only overlapping posteriorly. Coded as additive due to alignment of character states with the preferred phylogeny. Coding as non-additive in isolation or in unison with other additive multi-state characters does not affect polarization of the character/states or alter the phylogeny. State 1 is a synapomorphy for the tanymecine clade [non-focal], whereas state 2 is a synapomorphy for *Minyomerus* [JF2018].
2. Habitus, arrangement of elytral setae: (0) variously interspersed; (1) arranged in single-file rows on elytral intervals; (2) arranged non-uniformly on elytral intervals. Coded as additive due to alignment of character states with the preferred phylogeny. Coding as non-additive in isolation or in unison with other additive multi-state characters does not affect polarization of the character/states or alter the phylogeny. State 1 is a synapomorphy for the tanymecine clade [non-focal], whereas state 2 is a synapomorphy the *M. aeriballux–M. reburrus* clade [JF2015].
3. Habitus, lateral elytral setae and ventral setae differentiated from setae of elytral disc: (0) absent; (1) present. Homoplasy for *Minyomerus* [JF2018], with a reversal (state 0) in the *M. aeriballux-M. languidus* clade [JF2015], subsequent convergent gain (state 1) in the *M. bulbifrons-M. caseyi* clade [JF2018], and convergent reversal (state 0) in the *M. bulbifrons-M. puticalutus* clade [JF2015] (ci = 25; ri = 70).
4. Habitus, rows of elytral setae with larger white setae randomly interspersed among smaller brown setae: (0) absent; (1) present. Synapomorphy for the *M. caseyi-M. franko* clade [JF2018]. Changed from Jansen & Franz (2015), where *M. rutellirostris* [JF2015] was previously coded as having this character; however, the white elytral setae of this species are not randomly interspersed, but follow a distinct, and uniquely derived, pattern where every other interval contains a row of such setae.
5. Habitus, elytra and pronotum generally large, protuberant, and sculpted in appearance along dorsal and lateral faces: (0) absent; (1) present. Synapomorphy for the *M. microps-M. tylotos* clade [JF2018].
6. 6. Prementum, anterior margin medially with a distinct facet, rather than a single edge edge, that continues to lateral margins: (0) absent; (1) present. Synapomorphy for the *M. aeriballux–M. languidus* clade [JF2015], with a single reversal in the *M. caseyi–M. trisetosus* clade [JF2015] (ci = 50; ri = 75).
7. Prementum, strongly ligulate and with margins nearly straight, appearing pentagonal: (0) absent; (1) present. Synapomorphy for the *M. aeriballux–M. languidus* clade [JF2015], with independent reversals in the *M. caseyi–M. franko* clade [JF2018] and *M. griseus–M. rutellirostris* clade [JF2015], respectively (ci = 33; ri = 71).
8. Prementum, anterolateral margins simple, unexpanded: (0) absent; (1) present. Synapomorphy for the*M. aeriballux–M. languidus* clade [JF2015].
9. Prementum, anterolateral margins explanate, angular, and posteriorly declivous, with a distinctly hexagonal appearance: (0) absent; (1) present. ACCTRAN optimization preferred (see Agnarsson & Miller 2008), therefore inferred as a synapomorphy for the *M. constrictus–M. laticeps* clade [JF2015].
10. Prementum, exposure of palpomere I: (0) exposed, visible beyond ligula and anterior margin of prementum in ventral view; (1) hidden, fully covered or only minutely exposed beyond ligula and anterior margin of prementum in ventral view. Synapomorphy for the *M. aeriballux–M. microps* clade [JF2015], with a single reversal in *M. franko* [JF2018] (ci = 50; ri = 75).
11. Rostrum, form in dorsal view: (0) approximately quadrate; (1) somewhat conical, medially convex. Synapomorphy for the *M. aeriballux–M. reburrus* clade [JF2015].
12. Rostrum, form of nasal plate and demarcation of epistoma: (0) with three parallel, longitudinal cari-nae, and surface planar between these; (1) with a sharp, narrow, chevron-shaped carina demarcating epistoma; (2) with a broad, scale-covered, chevron-shaped carina demarcating epistoma. Coded as additive due to alignment of character states with preferred phylogeny. Coding as non-additive in isolation or in unison with other additive multi-state characters does not affect polarization of the character/states or alter the phylogeny. State 1 is a synapomorphy for the tanymecine clade [non-focal], whereas state 2 is a synapomorphy for *Minyomerus* [JF2018].
13. Rostrum, sulcus posteriad of nasal plate weakly impressed: (0) absent; (1) present. Convergently present in the *M. constrictus–M. laticeps* clade [JF2015], the *M. microps–M. tylotos* clade [JF2018], and the *M. gravivultus–M. griseus* clade [JF2015] (ci = 33; ri = 60).
14. Rostrum, form of sulcus posteriad of nasal plate: (0) absent; (1) sulcus present, broad, and weakly punctate; (2) sulcus present, more strongly punctate. Coded as additive due to alignment of character states with preferred phylogeny. Coding as non-additive in isolation or in unison with other additive multi-state characters does not affect polarization of the character/states or alter the phylogeny. Synapomorphy for *Minyomerus* [JF2018] (state 1) and the *M. aeriballux–M. cracens* clade [JF2015] (state 2), respectively.
15. Head, frons very strongly projected beyond anterior margin of eye, by 2× anterior-posterior length of eye: (0) absent; (1) present. Synapomorphy for the *M. bulbifrons–M. politus* clade [JF2015].
16. Head, frons with posterior transverse constriction: (0) absent; (1) present. Synapomorphy for the *M. aeriballux–M. languidus* clade [JF2015], with a single reversal in *M. griseus* [JF2015] (ci = 50, ri = 85).
17. Antenna, length of scrobe relative to funicle and club: (0) scrobe shorter than funicle and club combined; (1) scrobe subequal in length to funicle and club combined. Synapomorphy for *Minyomerus* [JF2018].
18. Antenna, terminal funicular segment entirely without thin, nearly setiform scales: (0) absent; (1) present. Convergently present in *M. cracens* [JF2015], *M. reburrus* [JF2015], *M. caseyi* [JF2015], and the *M. bulbifrons–M. politus* clade [JF2015] (ci = 25; ri = 25).
19. Antenna, terminal funicular segment at least partially clothed with broad scales: (0) absent; (1) present. Synapomorphy for the *M. aeriballux–M. cracens* clade [JF2018] with independent reversals in *M.franko* [JF2018] and the *M. gravivultus–M. griseus* clade [JF2015] (ci = 33; ri = 71).
20. Head, angle of base in relation to prothorax: (0) directed anteriorly, in line with main body axis; (1) directed strongly ventrally; (2) directed slightly ventrally. Coded as additive due to alignment of character states with preferred phylogeny. Coding as non-additive in isolation or in unison with other additive multi-state characters does not affect polarization of the character/states or alter the phylogeny. State 1 is a synapomorphy for the tanymecine clade [non-focal], whereas state 2 is a synapomorphy for *Minyomerus* [JF2018].
21. Pronotum, condition of post-ocular vibrissae: (0) present in a well-developed tuft of 10 or more setae; (1) present in a reduced tuft of 3-7 setae. Synapomorphy for the *M. aeriballux–M. imberbus* clade [JF2018].
22. Prosternum, intercoxal process complete, undivided: (0) absent; (1) present. Synapomorphy for the tanymecine clade [non-focal], with a single reversal for *Minyomerus* [JF2018] (ci = 50; ri = 66).
23. Prosternum, intercoxal process divided at midpoint between coxae, but both anterior and posterior processes extending completely between procoxae and contiguous with each other: (0) absent; (1) present. Synapomorphy for the *M. griseus–M. rutellirostris* clade [JF2015].
24. Legs, fore femora not swollen in comparison to other legs: (0) absent; (1) present. Synapomorphy for the *M. aeriballux-P. subcancer* clade [non-focal].
25. Legs, sculpture of ventral surface of protibiae: (0) evenly convex throughout; (1) with a longitudinal groove or concavity. Synapomorphy for the *M. aeriballux-I. debilis* clade [non-focal].
26. Legs, setation of metatibial apex: (0) bristles at least as long as surrounding setae and setiform; (1) bristles shorter than surrounding setae and conical; (2) bristles sub-equal in length to surrounding setae and somewhat lamelliform. Coded as additive due to alignment of character states with preferred phylogeny, and the appearance of being a transformation series. Coding as non-additive in isolation or in unison with other additive multi-state characters does not affect polarization of the character/states or alter the phylogeny. Synapomorphy for *Minyomerus* [JF2018] (state 1) and the *M. aeriballux–M. reburrus* clade [JF2015] (state 2), respectively.
27. Legs, curvature of metatibial apex: (0) convex; (1) oblique. ACCTRAN optimization preferred (see Agnarsson & Miller 2008), therefore inferred as a synapomorphy for *Minyomerus* [JF2018] with a reversal (state 0) in the *M. aeriballux–M. conicollis* clade [JF2015], then a convergent gain (state 1) in the *M. aeriballux–M. bulbifrons* clade [JF2018], with independent reversals (state 0) in the *M. aeriballux–M. reburrus* clade [JF2015], *M. gravivultus* [JF2015], *M. trisetosus* [JF2015], and the *M. bulbifrons–M. politus* clade [JF2015] (ci = 14; ri = 40).
28. Legs, relative length of mesotarsi to mesotibiae: (0) tarsi less than 3/4× length of tibiae; (1) tarsi at least equal in length to tibiae; (2) tarsi shorter than tibiae, but longer than 3/4 × length of tibiae. Coded as additive due to alignment of character states with preferred phylogeny. Coding as non-additive in isolation or in unison with other additive multi-state characters does not affect polarization of the character/states or alter the phylogeny. State 1 is a synapomorphy for the tanymecine clade [non-focal], whereas state 2 is a synapomorphy for *Minyomerus* [JF2018].
29. Legs, tarsi ventrally spinose: (0) absent; (1) present. Synapomorphy for *Minyomerus* [JF2018].
30. Elytra, humeral angle rounded, not projected: (0) absent; (1) present. Synapomorphy for the *M. aeriballux-I. buchanani* clade [non-focal].
31. Female terminalia, spermatheca with apical cylindrical bulb on corpus: (0) absent; (1) present. Synapomorphy for the *M. bulbifrons–M. puticulatus* clade [JF2015].
32. Female terminalia, corpus of spermatheca sinuate: (0) absent; (1) present. Synapomorphy for the *M. imberbus–M. sculptilis* clade [JF2018].
33. Female terminalia, lamina of spiculum ventrale less sclerotized between laminar arms: (0) absent; (1) present. Coded as inapplicable for *S. californicus* [non-focal], as laminar arms are not apparent. Synapomorphy for the *M. gravivultus–M. griseus* clade [JF2015].
34. Female terminalia, lamina of spiculum ventrale with laminar arms bifurcating around a membranous region: (0) absent; (1) present. Coded as inapplicable for *S. californicus* [non-focal], as laminar arms are not apparent. Synapomorphy for the *M. gravivultus–M. griseus* clade [JF2015].
35. Female terminalia, lamina of spiculum ventrale with style basally divided or obscured, not mesally intact: (0) absent; (1) present. Coded as inapplicable for *S. californicus* [non-focal], as laminar arms are not apparent. Synapomorphy for the *M. aeriballux–M. imberbus* clade [JF2015].
36. Female terminalia, lamina of spiculum ventrale with laminar arms clearly bifurcating. (0) absent; (1) present. Coded as inapplicable for *S. californicus* [non-focal], as laminar arms are not apparent. Synapomorphy for the *M. aeriballux–M. conicollis* clade [JF2015].
37. Female terminalia, laminar arms narrowly bifurcating basally, thereafter sub-parallel mesally: (0) absent; (1) present. Synapomorphy for the *M. aeriballux–M. ampullaceus* clade [JF2018].
38. Female terminalia, coxites of ovipositor with a lateral, anteriorly-directed, recurved, alate process: (0) absent; (1) present. Coded as inapplicable for *S. californicus* [non-focal], as coxites of ovipositor are not apparent. Synapomorphy for the *M. caseyi–M. trisetosus* clade [JF2015].
39. Female terminalia, relative length of styli to coxites of ovipositor: (0) Similar in size; (1) distinctly shortened; (2) highly reduced, appearing minute. Coded as non-additive, due to strong differences in structure of coxites and styli in state 2; inapplicable for outgroup taxa, as styli of ovipositor are not apparent. ACCTRAN optimization preferred (see Agnarsson & Miller 2008), therefore inferred as convergent gains in *M. franko* [JF2018] and the *M. bulbifrons–M. puticulatus* clade [JF2015] (state 1), with a single reversal in *M. bulbifrons* [JF2015] (state 0). Autapomorphy for *M. reburrus* [JF2015] (state 2) (ci = 50, ri = 0).
40. Female terminalia, condition of medial, anteriorly-directed, sclerotized process of coxites of ovipositor: (0) fully developed; (1) reduced and inapparent. Coded as inapplicable for *S. californicus* [non-focal], as coxites of ovipositor are not apparent. Synapomorphy for the *M. aeriballux–M. cracens* clade [JF2015], with a single reversal in the *M. gravivultus–M. griseus* clade [JF2015] (ci = 50, ri = 83).
41. Female terminalia, anterior margin of tergum VII entirely free of sclerotized band: (0) absent; (1) present. Coded as inapplicable for *S. californicus* [non-focal], as the tergum VII is evenly sclerotized throughout. Convergently present in *M. aeriballux* [JF2015], *M. microps* [JF2015], and the *M. caseyi–M. trisetosus* clade [JF2018] (ci = 33; ri = 50).
42. Female terminalia, anterior margin of tergum VII sclerotized fully, appearing as an obviously complete band: (0) absent; (1) present. Coded as inapplicable for *S. californicus* [non-focal], as the tergum VII is evenly sclerotized throughout. Convergently present in *M. conicollis* [JF2015] and the *M. bulbifrons–M. puticulatus* clade [JF2015] (ci = 50; ri = 66).
43. Male terminalia, apical sclerite of aedeagal flagellum elongate-spiriform: (0) absent; (1) present. Synapomorphy for the *M. bulbifrons–M. politus* clade [JF2015].
44. Male terminalia, style of spiculum gastrale with an anterior ventral flange: (0) absent; (1) present. Synapomorphy for the *M. bulbifrons–M. caseyi* clade [JF2018].
45. Male terminalia, lamina of spiculum gastrale longer than broad and anteriorly extended along syle: (0) absent; (1) present. ACCTRAN optimization preferred (see Agnarsson & Miller 2008), therefore inferred as convergent gains in the *M. imberbus–M. sculptilis* clade [JF2018] and the *M. bulbifrons–M. puticulatus* clade [JF2015], with a reversal in *M. bulbifrons* [JF2015] (ci = 33; ri = 0).
46. Male terminalia, sub-triangular sclerites of sternum VIII with a medial process: (0) absent; (1) present. ACCTRAN optimization preferred (see Agnarsson & Miller 2008), therefore inferred as convergent gains in *M. cracens* [JF2015] and the *M. caseyi–textitM*. trisetosus clade [JF2015] (ci = 50, ri = 0).
47. Male terminalia, curvature of posterior margin of tergum VII: (0) evenly arcuate; (1) medially incurved. ACCTRAN optimization preferred (see Agnarsson & Miller 2008), therefore convergently present in the *M. constrictus–M. laticeps* clade [JF2015] and the *M. bulbifrons–M. gravivultus* clade [JF2015] with a reversal in *M. gravivultus* [JF2015] (ci = 33; ri = 66).
48. Male terminalia, tergum VII approximately 4× as long as broad: (0) absent; (1) present. Synapomorphy for the *M. griseus–M. rutellirostris* clade [JF2015].
49. Male terminalia, aedeagal pedon expanded laterally around ostium: (0) absent; (1) present. ACCTRAN optimization preferred (see Agnarsson & Miller 2008), therefore convergently present in the *M. constrictus–M. laticeps* clade [JF2015], *M. cracens* [JF2015], and the *M. caseyi–M. trisetosus* clade [JF2015] (ci = 33; ri = 33).
50. Male terminalia, aedeagal pedon broad basally, evenly tapering toward apex: (0) absent; (1) present. Synapomorphy for the *M. bulbifrons–M. politus* clade [JF2015].
51. Male terminalia, aedeagal pedon medially sclerotized along dorsum: (0) absent; (1) present. ACCTRAN optimization preferred (see Agnarsson & Miller 2008), therefore convergently present in the *M. imberbus–M. sculptilis* clade [JF2015], *M. cracens* [JF2015], and the *M. bulbifrons–M. gravivultus* clade [JF2015], with a reversal in *M. bulbifrons* [JF2015] (ci = 25; ri = 50).
52. Male terminalia, width of connection between apodemes of aedeagal tegmen: (0) wider than base of apodeme; (1) narrower than base of apodeme. Synapomorphy for the *M. aeriballux–M. bulbifrons* clade [JF2018], with a single reversal in the *M. griseus–M. rutellirostris clade* [JF201] (ci = 50; ri = 83).

## RCC-5 ALIGNMENTS

Details of our RCC-5 alignment approached are given in free text form in the **Supplemental Information SI1**, which also describes the content of the data input and output files. The latter, in turn, are appended in .txt, .csv, and .pdf format in the **Supplemental Information SI2 to S4**. All shown alignments are *intensional* in the sense of Franz & Peet (2009), and thus maximize high-level concept congruence where indicated, and in spite of non-congruent lower-level concept sampling.

The first, classification-based alignment (Fig. 32) is simple and straightforward to interpret (see also **Supplemental Information SI2**). We obtain high-level congruence among the concepts *Minyomerus* [JF2018] and *Minyomerus* [JF2015], where 17 species-level concepts are retained from Jansen & Franz (2015) and four species-level concepts are added in the current review. The coverage constraint is relaxed for *Minyomerus* [JF2015], thus allowing the four new species-level concepts to be subsumed under this parent. This is based on our assertion that they fall under the generic character circumscription of Jansen & Franz (2015).

The following two Figs. 33-34 show fully bifurcated, multi-phylogeny alignments of the same reasoner toolkit input, but resolved as whole concepts versus split concepts, respectively. In Fig. 33 (**Supplemental Information SI3**), we observe that the phylogenetic placements of *two* of the four new species-level concepts cause significant non-congruence in the alignment, resulting in seven overlapping RCC-5 articulations. *Minyomerus franko* [JF2018] is subsumed under the *M. caseyi–M. franko* clade [JF2018], which is intensionally congruent with the *M. caseyi–M. trisetosus* clade [JF2015]. In other words, this placement is not the source of non-congruence in the alignment. Similarly, the placement of *M. tylotos* [JF2018] into the new *M. microps–M. tylotos* clade [JF2018] is not conflicting in an intensional sense. At the next, more inclusive level, this addition “resolves” into the congruent *M. aeriballux–M. microps* clade [JF2018]/[JF2015].

In contrast, the placement of *M. ampullaceus* [JF2018] “inside” of *M. cracens* [JF2015] in the current phylogeny, generates five overlapping articulations among as many (five) non-congruent concept regions positioned 1-2 levels above these species-level concepts. The conflict is resolved in the next, more inclusive and congruent region of the *M. aeriballux–M. cracens* clade [JF2018] == *M. aeriballux–M. bulbifrons* clade [JF2015]

The placements of the previously circumscribed *M. imberbus* [JF2015] and the new species-level concept *M. sculptilis* [JF2018] – in relation to the congruent clade *M. constrictus–M. laticeps* [JF2018]/[JF2015] - cause two additional instances of overlap (Fig. 33). In the current phylogeny, *M. imberbus* [JF2015] is sister to *M. sculptilis* [JF2018], and placed “inside” ofthe *M. constrictus–M. laticeps* clade [JF2018]/[JF2015]. However, in the preceding phylogeny sec. Jansen & Franz (2015), *M. imberbus* [JF2015] is non-congruently included in the *M. constrictus–M. imberbus* clade [JF2015]. This conflict is only resolved at the level of *Minyomerus* [JF2018]/[JF2015].

Figure 34 (**Supplemental Information SI4**) shows that the inclusion of the four new species-level concepts in the *Minyomerus* [JF2018] phylogeny generates five split-concept regions for which there are no adequate labels in either input phylogeny. These labels correspond to the overlapping articulations mentioned above; in particular the non-congruent assignments of *M. ampullaceus* [JF2018], *M. cracens* [JF2018], and *M. sculptilis* [JF2018]. The phylogenetic character evidence for these placements and relationships are discussed in the following sections.

## DISCUSSION

### Relationships to the previous revision

The differences of the current phylogeny (Figs. 30-31) in relation to that of Jansen & Franz (2015) are in large part due to the unique character combinations present in the newly added species (Rieppel 2007, Franz 2014). Nonetheless, three main clades are resolved with strong support, and further corroborate the topology of Jansen & Franz (2015), as follows:

1. *Minyomerus* [JF2018] is strongly supported by the same eight synapomorphies identified in Jansen & Franz (2015). These are reiterated in the Introduction (Bremer support value [henceforth: bsv] = 10, relative fit difference [henceforth: rfd] = 95; Bootstrap [henceforth: boot] = 100).
2. *Minyomerus griseus* [JF2015] forms a well-supported clade with *M. rutellirostris* [JF2015] (bsv = 4, rfd = 77, boot = 96). These taxa jointly share the same two synapomorphies (chars. 23:1 and 48:1) provided in Jansen & Franz (2015): (1) the intercoxal process is divided at the midpoint between the coxae, but has both the anterior and posterior processes extending completely between the procoxae and contiguous with each other; and (2) the male tergum VII is nearly 4× as long as broad, respectively. In addition, the *M. gravivultus–M. griseus* [JF2015] clade (bsv = 3, rfd = 60), as resolved in the current cladogram, is congruent with that of Jansen & Franz (2015).
3. *Minyomerus* [JF2018] is nested within a well-supported clade of Tanymecini [non-focal] (boot = 100). However, further work is needed to assess the phylogenetic relationships between all genera presently assigned to the Tanymecini [non-focal] (Alonso-Zarazaga & Lyal 1999).

### Intrageneric relationships

Within *Minyomerus* [JF2018], beginning at the earliest-bifurcating node and proceeding towards the leaves, the first major incongruence with *Minyomerus* [JF2015] is the placement of *M. imberbus* [JF2015]. This species was sister to the *M. constrictus–M. laticeps* [JF2015] clade, which in turn was sister to the *M. aeruballux–M. conicollis* clade [JF2015]. The present analysis places *M. imberbus* [JF2015] in a clade with *M. sculptilis* [JF2018] (see **Placement of newly described species**). The *M. aeriballux–M. imberbus* clade [JF2018] (bsf = 2, rfd = 50) is supported by three synapomorphies: (1) presence of a transverse constriction across the posterior of the frons (char. 16: 1); (2) presence of a reduced tuft of post-ocular vibrissae (char. 21: 1); and (3) a mesally obscure lamina of the spiculum ventrale in the female (char. 35: 1).

We resolve *M. cracens* [JF2015] as sistertothe *M. aeriballux–M. bulbifrons* [JF2018] clade, inclusively supported by three synapomorphies: (1) presence of a strongly punctate sulcus posteriad of the nasal plate (char. 14: 2); (2) presence of broad scales on the terminal funicular segment of the antennae(char. 19: 1); and (3) absence of a medial, anteriorly-directed, sclerotized process on the coxites of the ovipositor (char. 40: 1).

The *M. aeriballux–M. bulbifrons* [JF2018] clade is weakly supported by a single synapomorphy: the width of the connection between the apodemes of the aedeagal tegmen is narrower than the base of the apodeme (char. 52: 1). Within this clade, the position of the *M. bulbifrons–M. caseyi* clade [JF2018] clade as separate from, and sister to, the *M. aeriballux–M. ampullaceus* clade [JF2018], is supported by one synapomorphy and one homoplasious character, namely: (1) presence of an anterior ventral flange on the style of the spiculum gastrale (char. 44: 1 – synapomorphic), and (2) differentiation of the setae on the lateral portion of the elytra and on the venter from the setae on the elytral disc (char. 3: 1 – homoplasious).

### Placement of newly described species

Clades within *Minyomerus* [JF2018] not addressed in the preceding section are identical in topology and composition to those of *Minyomerus* [JF2015], except for the addition of newly described species. Here we assess the phylogenetic placements of these species. We also discuss similarities in the biogeographic range of each species, in relation to the putative sister taxa, based on the results of species distribution modeling (see Figs. 35-38).

#### *Minyomerus sculptilis* [JF2018]

*Myniomerus sculptilis* [JF2018] is inferred as sister to *M. imberbus* [JF2015]. The *M. imberbus–M. sculptilis* clade [JF2018] (bsv = 3, rfd = 72) is supported by a single synapomorphy and two homoplasious characters: (1) corpus of spermatheca sinuate (char. 32: 1 – synapomorphic); (2) lamina of spiculum gastrale in male longer than broad and anteriorly extended along style (char. 45: 1 – homoplasious); and (3) aedeagal pedon medially sclerotized along dorsum (char. 51:1-homoplasious). In addition to these characters, *M. imberbus* [JF2015] and *M. sculptilis* [JF2018] share a general external gestalt, which makes separating these two species difficult, especially in damaged or worn specimens.

Whereas *M. sculptilis* [JF2018] is associated with big sagebrush (*Artemisia tridentata* [non-focal], tumbleweed (*Salsola tragus* [non-focal], and tall tumblemustard (*Sisymbrium altissimum* [non-focal]; its sister taxon *M. imberbus* [JF2015] is associated with budsage (*Artemisia spinescens* [non-focal]. The divergence of these two species may have been driven in part by differences in host-plant use. However, this is less likely considering the generalist feeding habits of *Minyomerus* [JF2018] congenerics. Conversely, their divergence may have resulted from a vicariance event, based on their present-day biogeographic distributions, which are separated by the eastern extension of the Columbia Plateau. *Minyomerus sculptilis* [JF2018] appears to be endemic to the Snake River Plain to the north, whereas *M. imberbus* [JF2015] has been found in the Great Basin Desert to the south.

#### *Minyomerus tylotos* [JF2018]

*Minyomerus tylotos* [JF2018] is sister to *M. microps* [JF2015]. The *M. microps–M. tylotos* clade [JF2018] (bsv = 3, rfd = 73) is supported by a single synapomorphy and a single homoplasious character: (1) elytra and pronotum generally large, protuberant, and sculpted in appearance along dorsal and lateral faces (char. 5: 1 – synapomorphic); and (2) sulcus posteriad of nasal plate broad and weakly punctate (char. 13: 1 – homoplasius). In addition to these characters, the two species share a similar gestalt and uniform setation.

*Minyomerus tylotos* [JF2018] appears to be endemic to northern Chihuahuan Desert, whereas *M. microps* [JF2015] is widely distributed to the north throughout the Great Plains and along the Missouri River. We consider it likely that *M. microps* [JF2015] represents a northern radiation of the common ancestor of this clade. Conversely, *M. tylotos* [JF2018] may represent the ancestral distribution to the south, based on the hypothesized origin of *Minyomerus* [JF2018] in the Chihuahuan Desert; see Jansen & Franz (2015) and Wilson & Pitts 2010.

#### *Minyomerus ampullaceus* [JF2018]

*Minyomerus ampullaceus* [JF2018] is sister to the *M. aeriballux–M. reburrus* clade [JF2015]. The *M. aeriballux–M. ampullaceus* clade [JF2018] (bsv = 1, rfd = 50) is supported by a single synapomorphy: lamina of spiculum ventrale with laminar arms basally bifurcating and sub-parallel mesally thereafter (char. 37: 1). The placement of this species is tentative and based on the characteristics of a single, worn specimen.

Nonetheless, the biogeographic distributions of the species in the *M. aeriballux–M. ampullaceus* clade [JF2018] exhibit overlap. *Minyomerus ampullaceus* [JF2018] is documented from Carlsbad, New Mexico, in the western parts of the distributions of *M. aeriballux* [JF2015] and *M. reburrus* [JF2015]. The divergence of the latter two species is thought to be a result of their habitat and host plant preference, given their overlapping ranges. *Minyomerus aeriballux* [JF2015] is found in very sandy soils and on dune systems, whereas *M. reburrus* [JF2015] prefers arid grasslands. Without additional distributional or host plant data for *M. ampullaceus* [JF2018], we cannot assess whether the single documented locality for this species represents the center or edge of its range. However, this locality does overlap with the known range of its sister clade, suggesting that the divergence of *M. ampullaceus* [JF2018] from the *M. aeriballux–M. ampullaceus* clade [JF2018] was not a vicariance event.

#### *Minyomerus franko* [JF2018]

*Minyomerus franko* [JF2018] is sister to the *M. caseyi–M. trisetosus* clade [JF2015]. The *M. caseyi–M. franko* clade [JF2018] (bsv = 4, rfd = 63) is supported by a single synapomorphy and two homoplasious characters: (1) rows of setae on elytral intervals comprised of larger white setae randomly interspersed among smaller brown setae(char. 4: 1 – synapomorphic); (2) prementum lacking strong ligula and straight margins, not appearing pentagonal (char. 7: 0 – homoplasious); and (3) anterior margin of female tergum VII entirely free of sclerotized band (char. 41:1-homoplasious). In addition to these characters, members of this clade share a generally similar gestalt, especially regarding the head and rostrum, and the articulation between the pronotum and elytra in dorsal and lateral view. The interspersed, white elytral setae of these three species exhibit varying degrees of apical expansion, and can appear moderately to greatly explanate or spatulate in at least some, but not all, specimens.

*Minyomerus franko* [JF2018] has been documented on spear globemallow *Sphaeralcea hastulata* [non-focal]. *Minyomerus trisetosus* [JF2015] is associated with broomweed *Xanthocephalum* [non-focal], creosote bush *Larrea tridentata* [non-focal] and snakeweed *Gutierrezia* [non-focal]. *Minyomerus caseyi* has no known plant associations. It is therefore possible that the divergence of *M. franko* [2018] was facilitated by differences in host-plant preference. However, this remains unlikely given the generalist feeding habits of congenerics.

Alternatively, the speciation sequence in the *M. caseyi–M. franko* clade [JF2018] may correspond to vicariance events. *Minyomerus trisetosus* [JF2015] inhabits a broad swath of the northern Chihuahuan Desert, whereas *M. franko* [JF2018] and *M. caseyi* [JF2015] are exclusively encountered in the southern Chihuahuan Desert. MaxEnt predicts overlapping species distributions for the latter two species. However, the *documented* localities of these two species pertain to distinct biogeographic regions. *Minyomerus franko* [JF2018] has only been collected in the valleys of the Sierra Madre Oriental range, whereas *M. caseyi* [JF2015] is found along the western edge of this range, in the eastern portion of the Central Mexican Plateau. Additional occurrence records are needed to clarify the spatial extents of these species’ distributions, and thus draw more robust inferences regarding their endemicity.

## CONCLUSIONS

Through addition of four herein described species, the entimine [non-focal] genus *Minyomerus* [JF2018] is expanded to include 21 species. We predict that additional undescribed species of *Minyomerus* [JF2018] exist throughout the North American deserts, given the narrow endemicity patterns of many members of the genus. Furthermore, we believe that sampling in poorly-sampled locales, particularly in the northwestern United States and in northern Mexico, will yield new evolutionary insights for this group.

New molecular data can strengthen phylogenetic hypotheses and provide estimates regarding the timing of diversification of *Minyomerus* [JF2018], thereby testing our current inference of an origin in central Mexico. Another research direction should focus on the reproductive behavior of certain species suspected to be parthenogenetic; including rearing and karyotyping. Finally, the validity of the genus *Minyomerus* [JF2018] as a member of the Tanymecini [non-focal], and its relationships to other Entiminae [non-focal], remain uncertain.

## ACKNOWLEDGMENTS

The authors are grateful to Robert Anderson (**CMNC**), Ed Riley and John Oswald (**TAMU**), and Lourdes Chamorro (**USNM**) for their assistance and provision of specimens used in this study. The authors also thank Salvatore Anzaldo, Andrew Johnston, Sangmi Lee and other ASUHIC members for their assistance in procuring, treating, and maintaining specimen loans upon entry into the ASUHIC.

## SUPPLEMENTAL INFORMATION

**SIl** Explanation of the RCC-5 alignment approach. File format: .pdf

SI2A Input constraints for the *Minyomerus* [JF2018]/[JF2015] rank-only classification alignment. File format: .txt

**SI2B** Input visualization for the SI2A input. File format: .pdf

SI2C Set of 114 *Maximally Informative Relations* (MIR) for the SI2A input. File format: .csv SI2D Alignment visualization for the SI2A input. File format: .pdf

**SI3A** Input constraints for the *Minyomerus* [JF2018]/[JF2015] phylogeny alignment – whole-concept resolution with overlap. File format: .txt SI3B Input visualization for the SI3A input. File format: .pdf

**SI3C** Set of 925 *Maximally Informative Relations* (MIR) for the SI3A input. File format: .csv SI3D Alignment visualization for the SI3A input. File format: .pdf

**SI4A** Input constraints for the *Minyomerus* [JF2018]/[JF2015] phylogeny alignment – split-concept resolution. File format: .txt SI4B Input visualization for the SI4A input. File format: .pdf

**SI4C** Set of 925 *Maximally Informative Relations* (MIR) for the SI4A input. File format: .csv

**SI4D** Alignment visualization for the SI4A input. File format: .pdf

## FUNDING STATEMENT

This research was funded in part by the National Science Foundation (DEB-1155984) and the United States Department of Agriculture – Agricultural Research Service (Agreement 58-1275-1-335). There was no additional external funding received for this study.

